# Exploring phenotypic diversity: a comparative analysis of cichlid convergence

**DOI:** 10.1101/2024.01.23.576830

**Authors:** Mariana Leal-Cardín, Seraina E. Bracamonte, Javier Aldegunde, Alan Hudson, Isabel S. Magalhaes, Claudia Patricia Ornelas-García, Marta Barluenga

## Abstract

Convergent evolution of similar phenotypes suggests some predictability in the evolutionary trajectories of organisms, due to strong and repeated selective pressures, and/or developmental constraints. In adaptive radiations, particularly in cichlid fish radiations, convergent phenotypes are commonly found within and across geographical settings. There are some main axes of cichlid morphological diversification. Recurrent changes in body patterns reveal adaption to alternative habitats, and modifications of the trophic apparatus respond to the exploitation of different food resources. Here we compare two Neotropical cichlid assemblages, the Mexican desert cichlid, and the Nicaraguan Midas cichlid, with similar polymorphic body and trophic apparatus patterns despite their independent evolution and evaluate morphological and gene expression convergence and divergence. We found a single morphological axis of differentiation in pharyngeal jaws with equivalent papilliform and molariform morphotypes in both cichlid radiations. In contrast, we found two different axes of differentiation in the shape of the body, defining two alternative limnetic body patterns. Genetic differences between morphotypes seem to be specific to each radiation, with no clear patterns of convergence.

## Introduction

Convergent evolution is an outstanding process pointing to some predictability in the evolutionary trajectories of organisms under similar selective pressures. Convergence, which is defined as the development of similar features in non-related taxa, often occurs as a response to common ecological patterns (Endler, 1986; Schluter, 2000; Losos, 2011). The phenomenon of phenotypic convergence has been extensively studied over the years, revealing that it frequently arises when organisms are subjected to similar selective pressures (Hulsey, 2009; McGee & Wainwright, 2013; Rosenblum *et al., 2014;* Hulsey *et al*., 2018; Feiner *et al*., 2020). Whether convergent traits arise from shared genotypic changes is gaining increased attention. Recent genomic analyses have explored the molecular architecture of convergence, and the concept of the predictability of genetic evolution is emerging as a fascinating idea (Conte *et al*., 2012; Stern, 2013; Chabrol *et al*., 2018; Sackton & Clark, 2019; Moran *et al*., 2023). While convergent phenotypes may result from divergent evolution of genetic mechanisms (e.g., Cooper *et al*., 2014), data is increasingly indicating that convergence often stems from the recurrent utilization of the same genes (Cresko *et al*., 2004; Protas *et al*., 2006; Gross *et al*., 2009; Chan *et al*., 2010; Poore *et al*., 2023). In the line of this thought, convergent evolution can result from the development of similar or identical mutations in independent lineages by the repetitive sorting of pre-existing alleles (Barret & Schluter, 2008), or the incorporation of foreign alleles through hybridization (as discussed in Stern, 2013). Additionally, genomic alterations in regulatory regions have been demonstrated to significantly contribute to convergent phenotypic evolution (Cresko *et al.,* 2004; Gross *et al.,* 2009; Stern, 2013).

Adaptive radiations involve a burst of diversification within an evolutionary lineage in response to the emergence of many vacant ecological niches (Givnish, 1997). Evolutionary radiations often provide compelling examples of phenotypic convergence, with similar traits emerging in geographically distant yet analogous environments (Losos, 2011). Anole lizards, for instance, have developed analogous eco-morphs on different Caribbean islands (e.g., Losos *et al.,* 1998; Losos, 2011; Mahler *et al*., 2013). Hawaiian spiders have also evolved equivalent eco-morphs on another archipelago (Gillespie, 2004). Similarly, three-spine sticklebacks have repeatedly given rise to trophically similar species pairs in temperate freshwaters (e.g., Schluter & Nagel, 1995; Kaeuffer *et al*., 2012; Poore *et al*., 2023). Convergent evolution is also evident in cichlid fish across the Great East African lakes (Kocher *et al*., 1993; Rüber & Adams, 2001; Muschick *et al*., 2012).

Cichlid fish adaptive radiations serve as an excellent model for evolutionary and ecological research, as evidenced by numerous studies (e.g., Kocher, 2004; Seehausen, 2006; Barluenga *et al*., 2006; Salzburger, 2018; Elmer *et al*., 2014; Franchini *et al*., 2014; Vanhove *et al.,* 2016; Meier *et al*., 2018; Ronco *et al*., 2021; Lozano-Martin *et al*., 2023). One of the primary drivers of diversity in cichlids is the ecological specialization to various habitats and food resources (e.g., Fryer & Iles, 1972; Salzburger & Meyer, 2004; Barluenga *et al*., 2006; Muschick *et al*., 2012; Kusche *et al*., 2014). The process of diversification in cichlids while occupying alternative habitats has been proposed as the initial stage of cichlid differentiation in adaptive radiations (Danley & Kocher, 2001; Streelman & Danley, 2003; Ronco *et al*., 2021), often associated with variation in body plans (Klingenberg *et al*., 2003; Clabaut *et al.,* 2007; Burns & Sidlauskas, 2018). Similar evidence in other fish taxa suggests that habitat shifts linked to body shape differentiation commonly represent the first step in ecological speciation (Schluter & McPhail, 1992; Robinson & Wilson, 1994; Rutschmann *et al*., 2011). The prevalent pattern observed in both freshwater and marine fishes involves recurrent transitions between littoral and pelagic habitats (e.g., three-spine sticklebacks *Gasterosteus aculeatus*, arctic chars *Salvelinus alpinus*, sunfishes *Lepomis spp*., whitefishes *Coregonus spp*.; Schluter & McPhail, 1992; Robinson & Wilson, 1994; Rogers & Bernatchez, 2007; Vonlanthen *et al*., 2012; Skoglund *et al*., 2015). These shifts have functional implications on body form, with littoral species repeatedly evolving deeper bodies for increased maneuverability (i.e., benthic), while pelagic species develop a more elongate, slender body shape, thought to be adapted for steady locomotion in open water (i.e., limnetic) (Robinson & Wilson, 1994). Divergence in body shape related to environmental features stands out as one of the most common patterns of diversification in fishes, with its genetic basis attributed to multiple genes of moderate to small effect (Gagnaire *et al*., 2013; Rogers & Bernatchez, 2007; Feulner & Seehausen 2019, De Lorenzo *et al*., 2023).

Diversification in cichlid species often undergoes a second stage associated with distinct diets (Danley & Kocher, 2001; Streelman & Danley, 2003; Ronco *et al.,* 2021). Cichlids showcase a diverse array of trophic phenotypes, characterized by significant variations in their feeding apparatus (Fryer & Iles, 1972; Rüber *et al*., 1999; Muschick *et al*., 2012). The evolution of a distinctive pharyngeal jaw apparatus (Fryer & Iles, 1972) underlies this diversity in feeding structures, enabling heightened specialization of the oral and pharyngeal jaws—an innovation crucial for their evolutionary success. Secondary divergence related to microhabitat partitioning and associated trophic resources is a recurring pattern not only in cichlids but also in other vertebrate adaptive radiations, such as Galapagos finches and anoles lizards (Streelman & Danley, 2003; Grant & Grant, 2009; Losos, 2009).

Most cichlid evolutionary studies have focus on the species rich African lacustrine assemblages in the Great East African lakes (Fryer & Iles, 1972; Seehausen, 2006; Salzburger, 2018; Ronco *et al.,* 2021). The Neotropical region holds several relatively recent cichlid radiations that are increasingly gaining interest due to the possibilities they offer to study the early stages of diversification, and sometimes incomplete processes of complete species delimitations (Barlow, 1976; Kornfield & Taylor, 1983; Barluenga *et al*., 2006; Hulsey *et al*., 2006; Elmer & Meyer, 2011; Elmer *et al*., 2014; Kusche *et al*., 2014; Magalhaes *et al.,* 2015; Burres, 2016; Kautt *et al.,* 2020). Two Neotropical cichlid assemblages well-known for their diversity and polymorphism are those of the Midas cichlid (*Amphilophus spp*.) in the Nicaraguan lakes (Barlow, 1976), and the desert cichlid (*Herichthys minckleyi*) in the Cuatro Ciénegas Valley in Mexico (Kornfield & Taylor, 1983).

In this study, we investigated morphological and gene expression convergence in two phylogenetic and geographically distant Neotropical cichlid radiations, the Cuatro Ciénegas desert cichlid, and the Nicaraguan crater lake Midas cichlid. We focused on the known primary axes of morphological variation in cichlid fish, body shape and pharyngeal jaw, a key trophic trait. We employed geometric morphometrics and genomic tools to explore the hypothesis that under similar environmental conditions, fish evolve similar phenotypic adaptations, a phenomenon well-established in cichlid fish (e.g., Hulsey, 2009; Muschick *et al*., 2012; Härer *et al*., 2018). We assessed the extent of convergence between these two systems and examined similarity in the underlying expression profiles of comparable eco-morphotypes across cichlid assemblages.

## Material and Methods

### Study area and species sampling

We collected adult specimens of the desert cichlid from several pools in the Cuatro Ciénegas Valley, Mexico, during 2011 and 2012 (Fig. 1A), and of the Midas cichlid species complex in two Nicaraguan crater lakes (CL), Xiloá and Apoyo, between years 2009 and 2016 (Fig. 1B). The collection method involved the use of gill nets, and fish were euthanized with an overdose of MS222 on ice. All fish were measured, weighted, and photographed in a standardized position in the field for subsequent morphometric analyses. We dissected the spleen and conserved it in RNAlater.

**Figure 1.**
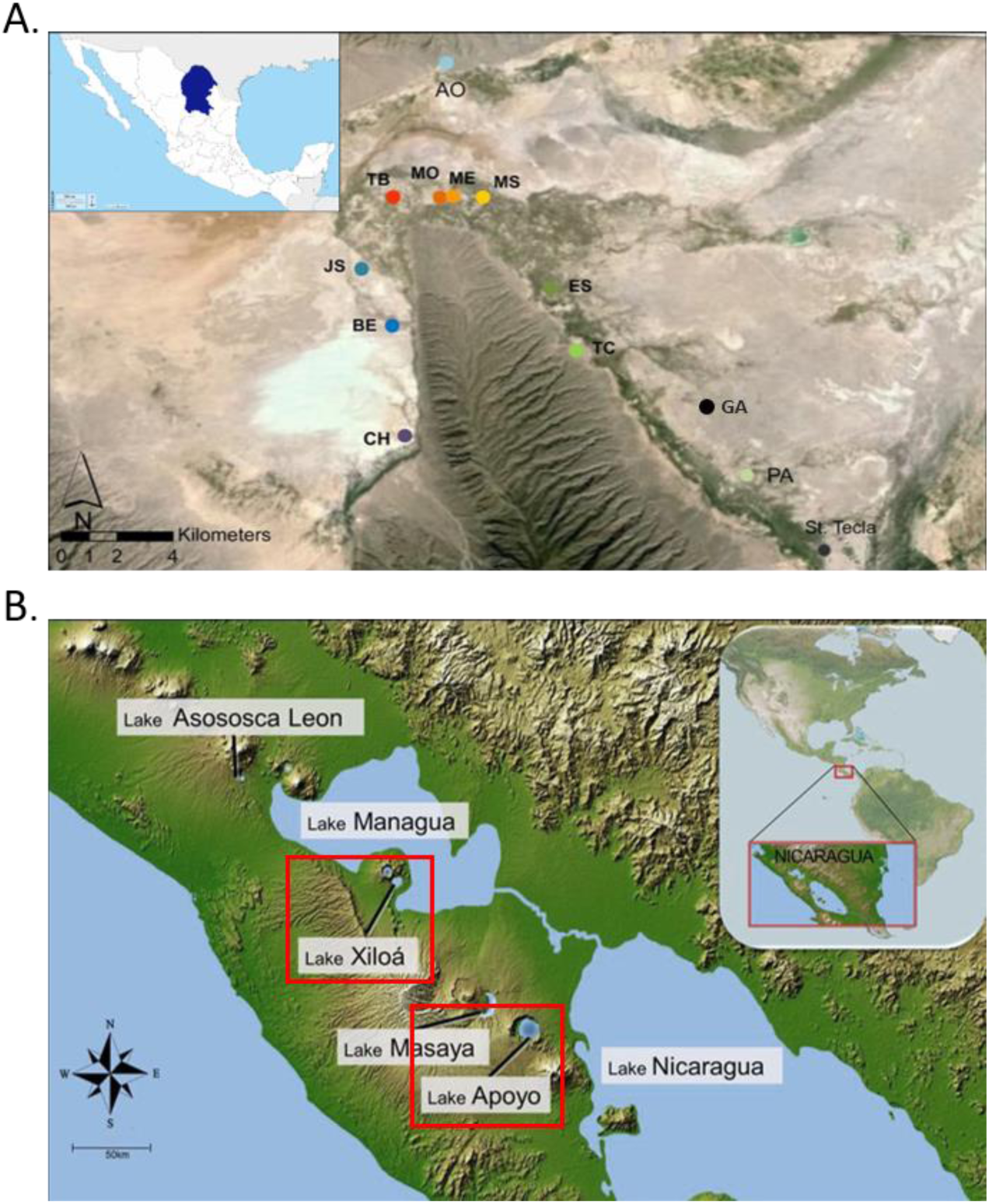
Map of the Cuatro Ciénegas Valley in Mexico (A) and the Pacific Coast of Nicaragua (B). Samples sites in Mexico included lakes Churince (CH), Becerra (BE), Juan Santos (JS), Tierras Blancas, (TB), Mojarral Oeste (MO), Mojarral Este (ME), Anteojo (AO), Escobedo (ES), Tío Cándido (TC), Pozas Azules (PA), Santa Tecla (St. Tecla) and rivers Mesquítez (MS) and Garabatal (GA). Crater lakes Apoyo and Xiloá in Nicaragua are framed in red.

Lower pharyngeal jaws (LPJ) were dissected in the field and preserved dry. In the laboratory, the jaws were cleaned, and the bone was scanned using an HP Scanjet G4050 scanner. Individuals were classified according to body and pharyngeal jaw morphological features. We analysed a total of 1008 fish, 463 from Mexico and 545 from Nicaragua (see Supplementary file 1 for detailed numbers and locations and information about phenotypic features).

The Mexican desert cichlid *Herichthys minckleyi* is phenotypically polymorphic in body shape, featuring benthic and limnetic fish, each with two distinct and divergent pharyngeal jaw types (molariform and papilliform) (Fig. 2; Kornfield & Taylor, 1983; Magalhaes *et al.,* 2015). Our sample included specimens representing all described morphotypes of the desert cichlid and are those included in Magalhaes *et al*. (2015). The Midas cichlid is also polymorphic in body shape and pharyngeal jaw shows recurrent morphotypes. In several Nicaraguan crater lakes there are recurrent eco-morphotypes of the Midas cichlid. In the two studied crater lakes, there is a littoral species preying on hard diets (CL Xiloá, *Amphilophus amarillo*; CL Apoyo, *A. astorquii*), a benthic profundal species (CL Xiloá, *A. xiloaensis*; CL Apoyo, *A. chancho*), and a pelagic species (CL Xiloá *A. sagittae*, CL Apoyo *A. zaliosus*). We included all six species in this study (Fig. 2).

**Figure 2.**
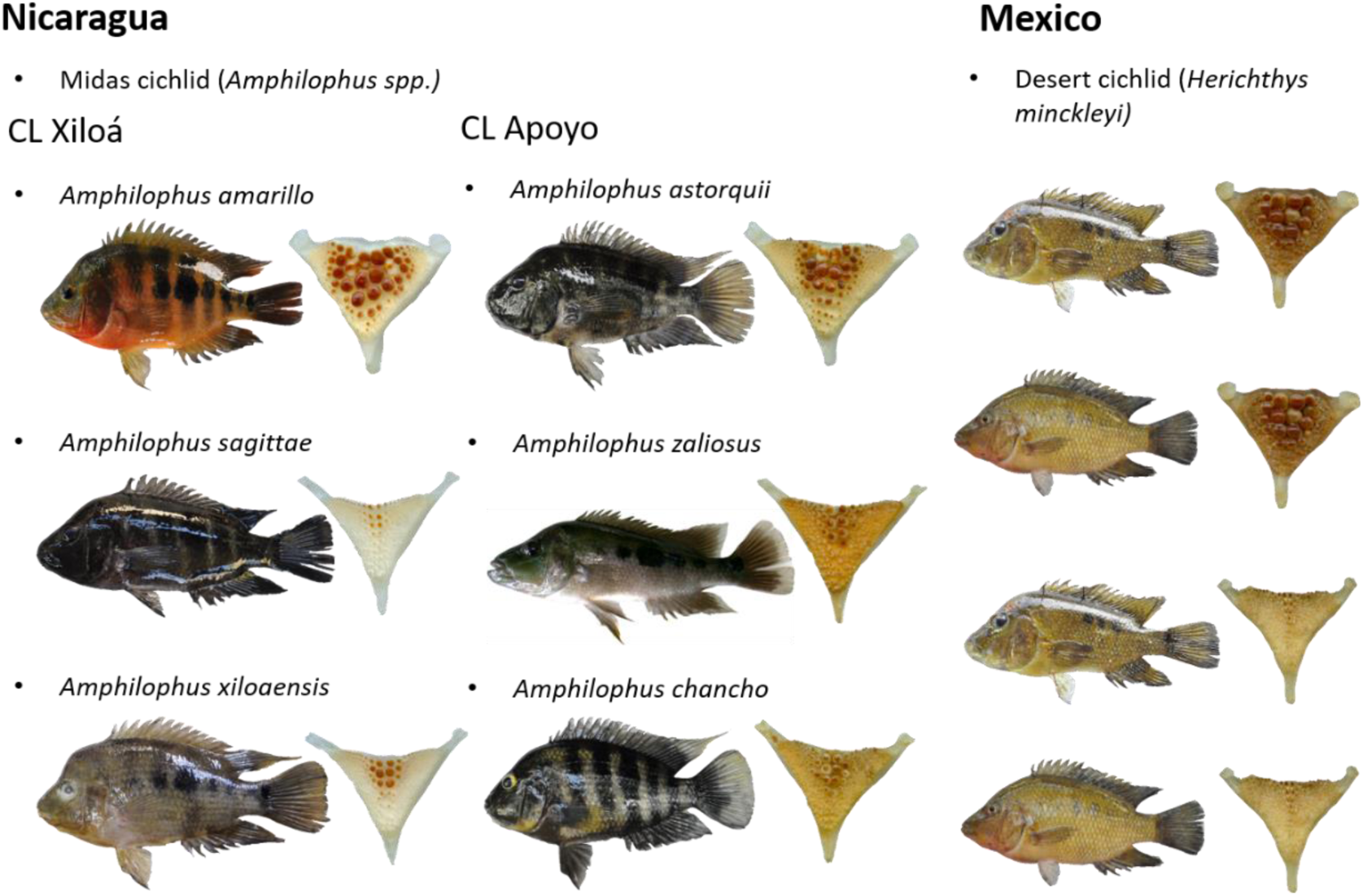
Polymorphisms of body and lower pharyngeal shapes in the two cichlid systems studied.

### Morphological analyses

We used landmark-based geometric morphometrics to evaluate the body and LPJ shape of fish. We transformed .jpg images to .tps images using TPSUTIL version 1.74 (Rohlf, 2015). We recorded eight landmarks describing the outline of the LPJ, and 15 landmarks across the fish body with TPSDIG2 version 2.30 (Rohlf, 2015). We used standard landmarks previously employed in similar studies (see Magalhaes *et al*., 2015; Sowersby *et al*., 2021; Fig.3).

**Figure 3.**
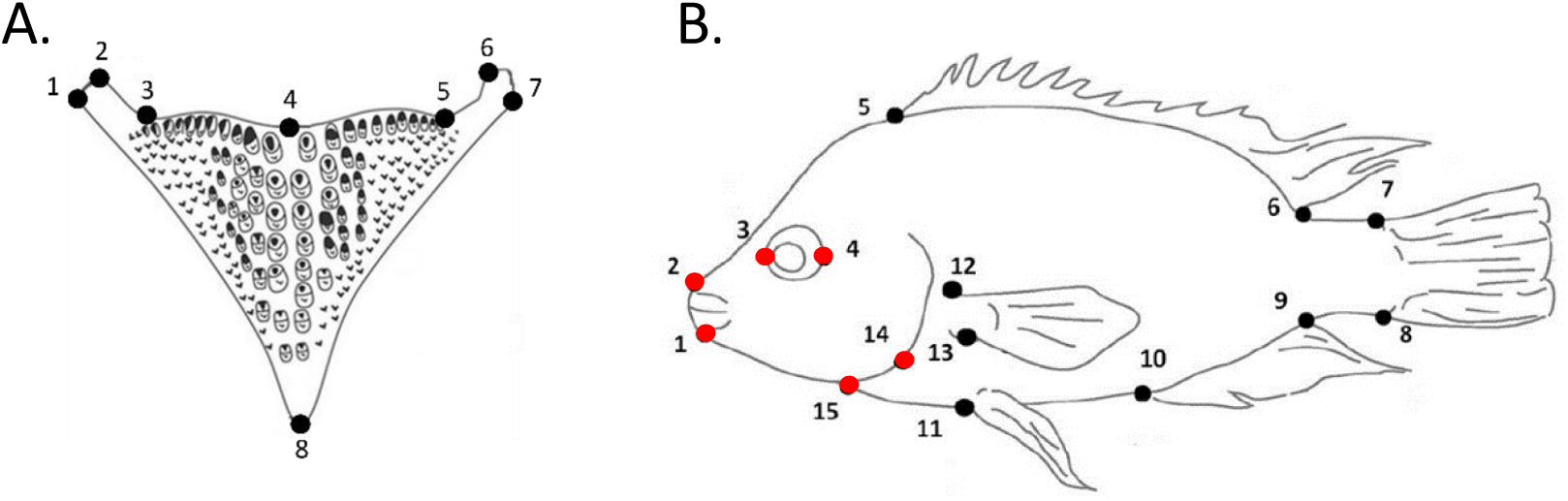
Landmark position for LPJ (A) and body shape (B) analyses. The different colours of the dots in the body shape figure indicate the different modules used for the modularity analysis.

We used the R package geomorph version 4.0.4 (Adams *et al.,* 2022) to analyse the configuration of the landmarks. To eliminate differences due to the scale, position, and orientation of the images we performed a Procrustes superimposition (Dryden & Mardia, 1998) with the *gpagen* function. This process aimed to extract a configuration of landmarks that retained only biologically meaningful information. The landmark coordinates were standardized to a unit centroid size, translated to a common position, and rotated to minimize Procrustes distances between all landmark configurations (Dryden & Mardia, 1998; Rohlf, 1999; Zelditch *et al*., 2004).

To assess allometric effects, we conducted multivariate regression analyses of individual Procrustes coordinates on centroid size for both body and LPJ shape using the *procD.lm* function. We retained the residuals of the regression for further analyses when the allometric effects were significant.

To characterize the variation in body and LPJ shape among individuals, we conducted principal component analyses (PCA) using the *gm.prcomp* function and constructed thin plate spline (TPS) diagrams. Phenotypic eco-morphotypes have been previously defined in Mexican and Nicaraguan cichlid radiations (Kornfield & Taylor, 1983; McKaye *et al.,* 2002; Stauffer *et al*., 2008; Magalhaes *et al*., 2015). To re-evaluate these clusters, we employed the k-means clustering algorithm with the *fviz.nbclust* function from the R package factoextra version 1.0.7 (Kassambara & Mundt, 2020), opting for the silhouette method (Kaufman & Rosseeuw, 2009) to determine the optimal number of clusters (k). In order to perform linear discriminant analyses (LDA), a model was constructed using a machine learning algorithm, with a 10-fold cross-validation method. For this approach, we used 80% of the dataset as training and the remaining 20% was dataset as test. The cross-validation was performed to determine the number of principal components (PCs) that maximized the variation. The LDA was constructed using the number of PCs obtained, and the posterior probabilities for each individual was calculated to be reassigned to their original group. We visualized the first linear discriminant function using density plots.

### Modularity of body shape

Modularity is a fundamental feature of organisms. Modules are the occurrence of highly integrated traits that are relatively independent from other modules (Klingenberg, 2008). Modularity is hypothesised to facilitate evolvability (Bonner, 1988; Raff, 1996; Wagner *et al*., 1996) because individual modules can change without compromising other modules (Hansen, 2003, 2006) and this has been proposed as one of the key factors to explain morphological diversity in ray-finned fishes (Ornelas-García et al. 2017; Larouche *et al*., 2015). With this reasoning, we defined two modules on the shape of fish, head, and trunk, and investigated their integration and relative variation.

We used two landmark configurations to test the morphological integration of the two module hypothesis in the body (head and trunk, see in Fig. 3 the landmarks defining head and trunk) with geomorph. We used the function *modularity.test* to quantify the degree of modularity of head and trunk Procrustes shape variables by calculating the covariance ratio (CR) coefficient (Adams & Collyer, 2019). A modular signal is revealed by values of CR coefficient lower than 1.

We also used six meristic measurements on the head and trunk taken from each fish image the size of the head and trunk with six additional measurements with TPSdig2. We used ANOVAs in R to evaluate differences in the ratio of head vs trunk, relative height, and relative length of the body.

### Gene expression analyses

#### RNA extraction and sequencing

We isolated RNA from spleen tissues obtained from seven individuals in CL Xiloá, eight individuals in CL Apoyo, and ten individuals from the Mexican pools. The spleen tissue was cut into small pieces, homogenized in 1 ml of TRIzol using a TissueLyser for 5 minutes at 25 Hz, and then centrifuged for 2 minutes at 13000 rpm at room temperature (RT). Following this, 200 µl of chloroform was added to the lysate, incubated for 2 minutes at RT, and centrifuged for 15 minutes at 4000 rpm at 4 °C. This step was repeated, with the final centrifugation lasting 5 minutes at RT.

Next, 500 µl of isopropanol were added to the aqueous phase containing the RNA, incubated for 30 minutes at RT, and centrifuged for 15 minutes at 13000 rpm at 4 °C. The resulting pellet was washed by resuspending it in 1 ml of 75% ethanol and centrifuging for 5 minutes at 13000 rpm at 4 °C. After removing the ethanol, the pellet was air-dried for 5-10 minutes. Subsequently, the pellet was resuspended in 100 µl of RNase-free water and dissolved at 55 °C.

The RNA was further purified using the RNeasy MinElute Cleanup kit (Qiagen) following the manufacturer’s recommendations with minor modifications. Specifically, the RNA bound to the spin column membrane was washed twice with RPE buffer instead of once with RPE and once with ethanol. The RNA was eluted in 30 µl of RNase-free water and incubated for 1 minute at RT before centrifugation.

To assess the quality and quantity of the RNA, an Agilent 2100 Bioanalyzer was used. Subsequently, libraries were prepared using the TruSeq RNA Sample Prep kit version 2 and sequenced on an Illumina HiSeq 2000 at GenePool at the University of Edinburgh. The reads were demultiplexed using bcl2fastq.

#### Read filtering and mapping

We used Trimmomatic version 0.39 (Bolger *et al*., 2014) to clip Illumina adapters, allowing for a maximum of 2 mismatches and extending to a score of 30. Reads were further processed by removing leading and trailing bases with a quality of less than 3. Additionally, reads were trimmed when the average quality within a 4-base sliding window fell below 15, and any reads shorter than 50 bp were excluded.

The trimmed reads were then mapped to the *Amphilophus citrinellus* reference genome (accession no. JACBYM000000000, annotation: Kautt *et al.,* 2020) using STAR RNA-seq aligner version 2.7.9a (Dobin *et al*., 2013) in multi-sample 2-pass mode. In the first mapping pass, newly identified splice junctions from all samples were collected and supplied to the second mapping pass. The strand was derived from the intron motif, and multi-mapping reads were aligned in random order.

Finally, we assembled the mapped reads from all samples into a consensus transcriptome using StringTie version 2.1.7 (Kovaka *et al*., 2019).

#### Differential gene expression

We quantified reads with featureCounts (Liao, Smyth, & Shi, 2014) implemented in the R package Rsubread version 2.4.2 (Liao, Smyth, & Shi, 2019), ignoring reads that mapped to multiple genomic regions and reads that overlapped multiple genes. We removed one sample from Mexico from the analysis due to a high number of unassigned reads. We identified differentially expressed genes (DEG) between jaw morphs and body morphs within and between systems with the Bioconductor package DESeq2 version 1.36.0 (Love, Huber, & Andres, 2014). We compared papilliforms vs molariforms, and limnetics vs benthics in the desert cichlid, in the Midas cichlid species complex, and across systems. We controlled for the lake of origin in Midas cichlid comparisons and geographic regions in cross-system comparisons. We further compared papilliform, molariform, benthic, and limnetic Midas cichlids between lakes. For this, we identified DEG between individuals from both lakes, irrespective of morphotype, to account for differences attributable to the lake of origin (i.e. between-lake DEG). For each morphotype, we then identified DEG between lakes and removed between-lake DEG to obtain a list of genes that can be attributed to differences within morphotype (i.e. within-morphotype DEG). Similarly, we identified within-morphotype DEG between systems accounting for between-system DEG.

#### Functional annotation

We extracted transcript sequences from the Amphilophus citrinellus genome with AGAT version 0.8.1 (Dainat, 2022) and predicted peptides with TransDecoder version 5.5.0 (Haas, 2018). We blasted the transcripts and the predicted peptides against the UniProtKB database Swiss-Prot. We accepted only hits with an e-value threshold < 0.001. We first excluded hits spanning < 50% of the query or subject length and selected the hit with the lowest e-value. We inferred Gene Ontology (GO) terms for each gene with Trinotate and used these for GO enrichment analyses with the R package topGO (Alexa & Rahnenführer, 2022). We performed Fisher’s exact tests with the elim algorithm (Alexa et al, 2006). We only considered GO terms with at least 5 annotated genes. We clustered enriched GO terms by semantic similarity with GOSemSim (Yu et al., 2010) for visualization.

## Results

### Pharyngeal jaw variation

Pharyngeal jaws exhibit polymorphism in both cichlid systems studied. In the Mexican system, a single species has been described with polymorphic LPJ (see Magalhaes *et al*. 2015). In the Nicaraguan system, we examined two lakes, each hosting different species with either papilliform or molariform LPJ. We observed a size-related effect on LPJ shape in all populations (*p* < 0.05) and consequently applied an allometric correction to all subsequent analyses. For all LPJ PCAs, 12 PCs were required to account for the entirety of shape variation, as shown below.

#### Pharyngeal jaw variation in the desert cichlid

Statistical analyses of morphometric data, including PCA and k-means, revealed the presence of two distinct types of LPJs in the desert cichlid (Figs. 4, S1A, S2A). Shape variation between the two jaw types, as indicated by PC1, ranged from LPJs with wider horns and a round, convex upper line (molariform) to LPJs with narrower horns and a concave upper line (papilliform), as illustrated in the TPS diagrams (Fig. 4).

**Figure 4.**
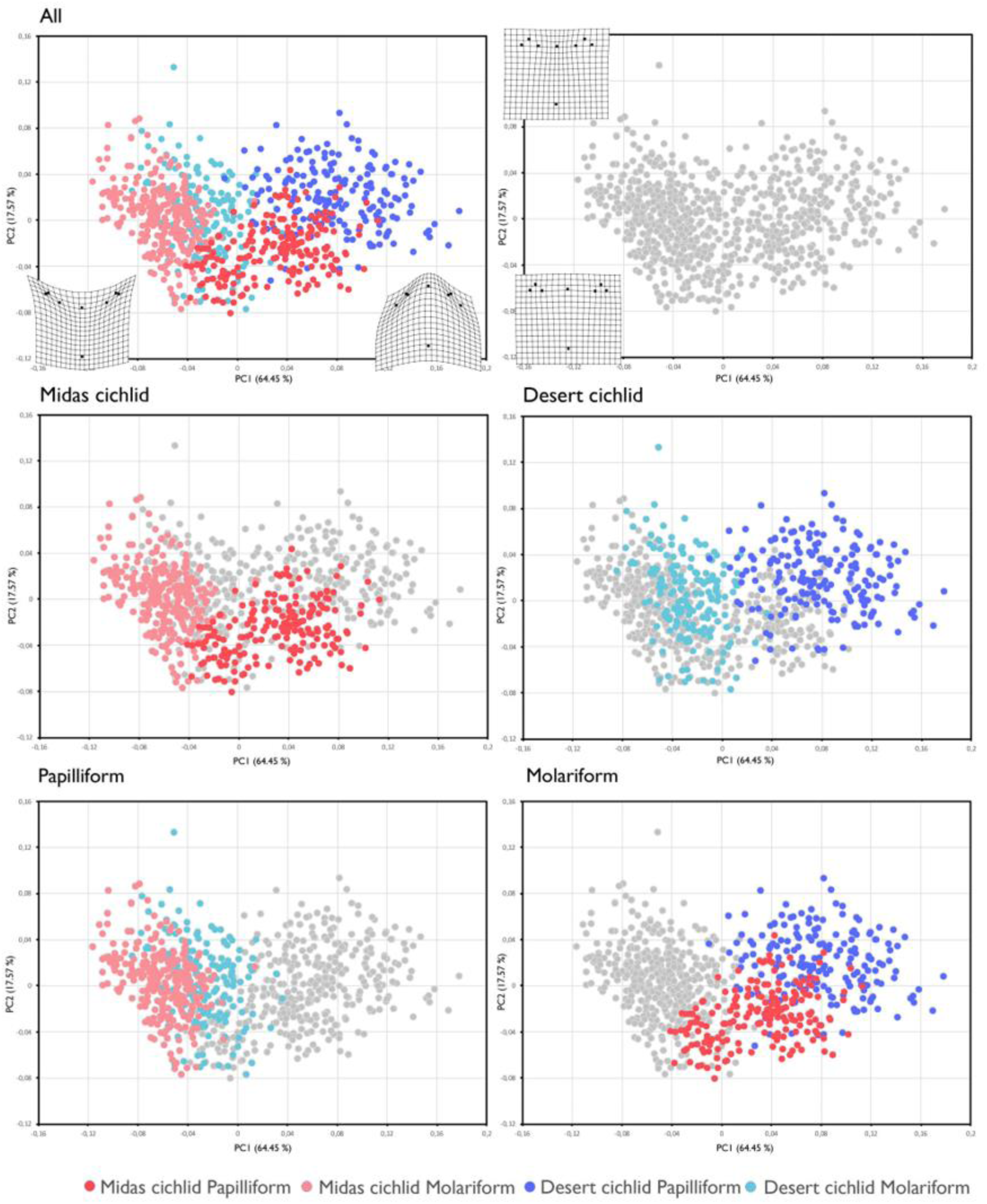
PCA showing the distribution of LPJ shape morphospace including both geographical regions together. PC1 separates morphotypes and geographical regions within each morphotype cluster. In Nicaragua, Midas cichlid shows differentiation between the two morphotypes, the same as what happens with the desert cichlid in Mexico. The papilliform type shows more overlap between regions than the molariform type.

For the LDA, we retained the first 9 PCs, resulting in a high overall proportion of correctly reassigned specimens (94%), with minimal overlap between morphotypes (Fig. 5A). Disparity analysis indicated that both morphotypes occupied an equivalent portion of morphospace (Procrustes variance, molariform: 0.004, papilliform: 0.005; *p* < 0.05).

**Figure 5.**
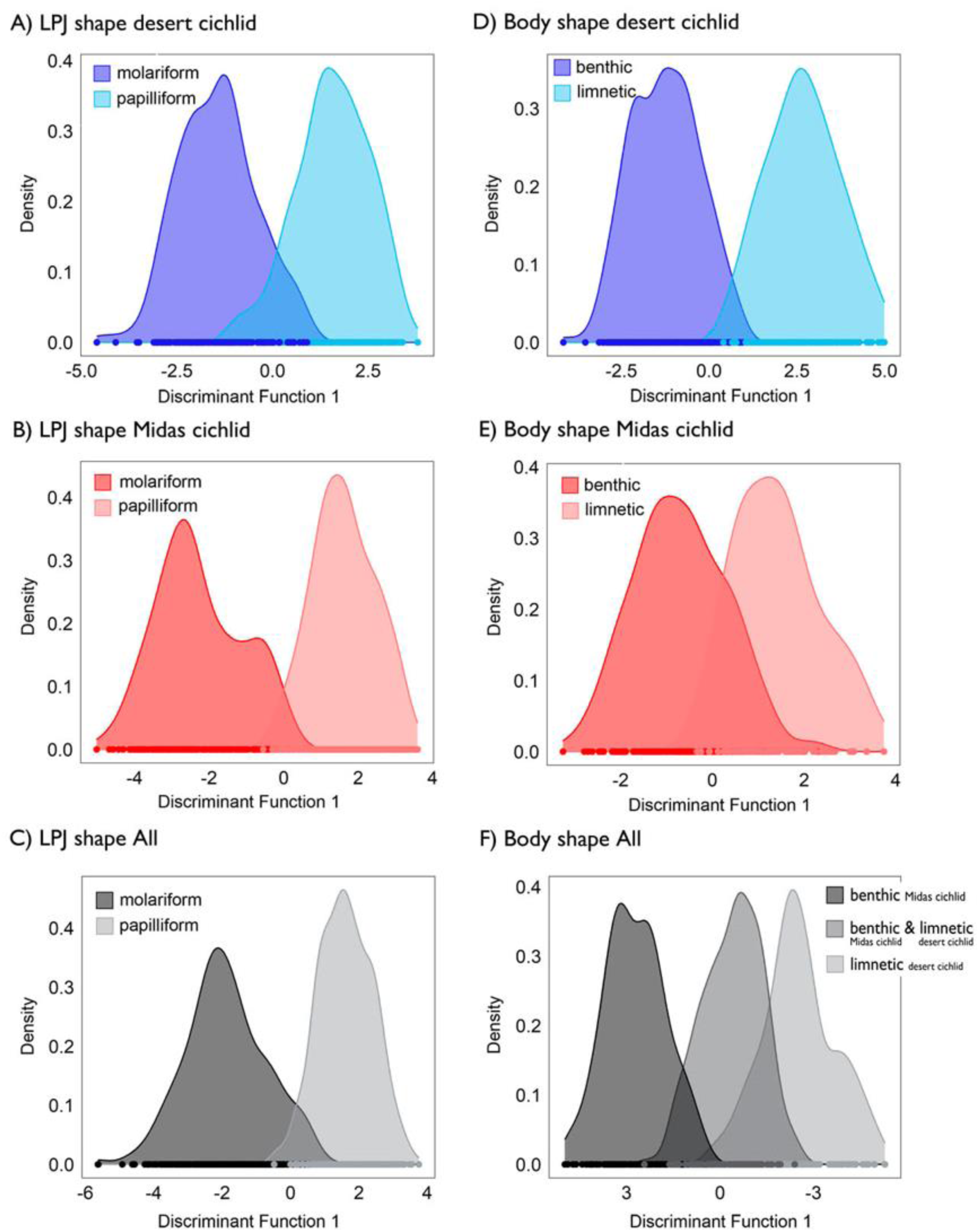
Linear discriminant function analyses differentiate two LPJ phenotypes in the two cichlid radiations desert cichlid (A), Midas cichlid (B), and also when all fish are analysed together (C). Two body shape phenotypes are differentiated in each cichlid radiation desert cichlid (D), Midas cichlid (E), but when all fish are analysed together three clusters are recovered by the k-means algorithm (F), differentiating the group of limnetic desert cichlids, from a group of limnetic Midas cichlids and benthic desert cichlids which overlapped, and from the group formed by all benthic Midas cichlids. Density plots show little overlap between jaw morphotypes in all comparisons and a larger overlap between body morphotypes.

#### Pharyngeal jaw variation in the Midas cichlid

The comprehensive examination of the LPJ shape across six Nicaraguan cichlid species inhabiting the two crater lake systems and the geometric morphometric analysis conducted to compare morphotypes within and between lakes confirm shape distinctions among species and evaluate the presence of intermediate morphotypes.

In CL Xiloá, morphometric analyses (PCA and k-means) revealed two distinct LPJ types: one molariform, including *A. amarillo*, and another papilliform, including all *A. xiloaensi*s and *A. sagittae*, consistent with previous descriptions (Figs. S1B, S2B). The LDA, with the first 3 PCs retained, achieved a remarkable 99% overall correct specimen reassignment, clearly differentiating the two morphotypes (Fig. 6A).

**Figure 6.**
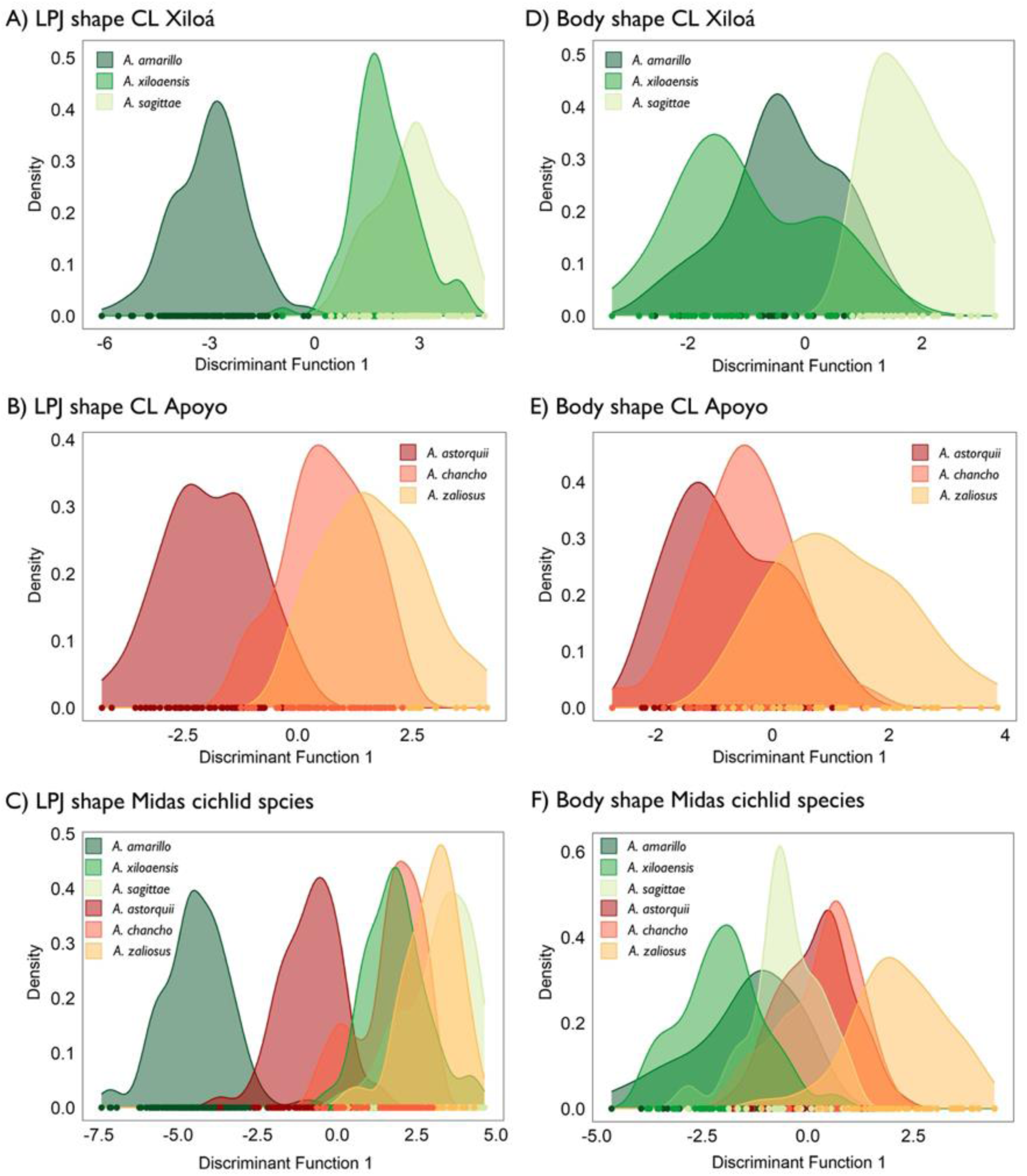
Linear discriminant function analyses differentiate eco-morphotypes within CL Xiloá and CL Apoyo in Nicaragua. Density plots show very marked differentiation between molariform and papilliform species in CL Xiloá (A), while this differentiation is less clear in CL Apoyo (B). The analysis including all species confirms the two jaw morphotypes (C), and shows stronger molarization in CL Xiloá. Density plots show differentiation between limnetic and benthic species in CL Xiloá (D) and CL Apoyo (E) with again more overlap among species in the later. The analysis including all species finds marked overlap in body shape between morphotypes (F).

The results from CL Apoyo were less clear. K-means clustering suggested an optimal number of three clusters, indicating LPJ shape differences among the three species (Figs. S1C, S2C). The LDA, with 7 retained PCs, resulted in an 81% overall correct specimen reassignment. However, a significant overlap was observed between the two papilliform LPJs (Fig. 6B).

Disparity analyses revealed greater morphological diversity among Midas cichlid species in CL Xiloá compared to those in CL Apoyo (Procrustes variance, CL Xiloá = 0.005, CL Apoyo = 0.003; *p* < 0.05), primarily due to the pronounced molarization of LPJ in *A. amarillo* (Fig. 4).

Comparing the morphospace within each crater lake, *A. amarillo* and *A. sagittae* occupied equivalent spaces in CL Xiloá, while *A. xiloaensis* occupied a significantly smaller portion (Procrustes variance, *A. amarillo*: 0.006, *A. xiloaensis*: 0.003, *A. sagittae*: 0.005). Similar trends were observed in CL Apoyo, where *A. astorquii* and *A. zaliosus* occupied equivalent spaces, while *A. chancho* occupied a significantly smaller portion (Procrustes variance, *A. astorquii*: 0.003, *A. chancho*: 0.002, *A. zaliosus*: 0.004).Comparing Midas cichlid species from both lakes, two differentiated LPJ clusters were identified, corresponding to each morphotype (Figs. 4, S1D, S2D). The LDA, with the first 6 PCs retained, achieved a high 97% overall correct specimen reassignment, displaying minimal overlap between the two morphotypes (Fig. 5B). Disparity analysis indicated that molariform jaws were more morphologically diverse than papilliform jaws (Procrustes variance, molariform: 0.006, papilliform: 0.004; *p* < 0.05).

Comparing papilliform LPJs from both Nicaraguan lakes revealed considerable shape overlap (see Fig. 6C), but k-means clustering suggested an optimal number of clusters of k = 2 (Figs. S1E, S2E). The LDA, with 10 retained PCs, achieved a 75% overall correct specimen reassignment. Disparity analysis indicated a larger papilliform morphospace in CL Apoyo compared to that in CL Xiloá (Procrustes variance, CL Apoyo: 0.003, CL Xiloá: 0.002; *p* < 0.05).

Comparing molariform LPJs from both lakes revealed significant differences between molariforms in each lake (see Fig. 6C), supported by k-means clustering analysis (k = 2; Figs. S1F, S2F). The LDA, with 10 retained PCs, achieved a 93% overall correct specimen reassignment. Disparity analysis indicated that the morphospace used by molariforms in CL Apoyo was larger than that used by molariforms in CL Xiloá (Procrustes variance, CL Apoyo = 0.003, CL Xiloá = 0.002; *p* < 0.05).

#### Pharyngeal jaw shape variation across radiations

We compared the LPJ shape of fish from the two different geographical regions to explore the degree of convergence between morphotypes and to identify potential geographical differences. A PCA through the two major axes (PC1 = 64.45%, PC2 = 17.57%; Fig. 4) revealed a single cloud of points without major discontinuities, but with morphotypes arranged by geographical region. PC1 ordered samples according to morphotype, ranging from most molariform with short and wide horns and highly convex upper lines in Mexico to most papilliform with long and narrow horns and concave upper lines in Nicaragua. Notably, some overlap was observed between Midas cichlid molariform jaws and desert cichlid papilliform jaws. PC2 ordered samples from longer LPJs in Mexico to shorter and wider LPJs in Nicaragua (Fig. 4).

The k-means clustering assigned individuals into two major morphological clusters (k = 2; papilliform *vs*. molariform), largely corresponding to previously defined molariform and papilliform jaws (Figs. S1G, S2G). The LDA for k = 2 retained 6 PCs, achieving a very high overall proportion of correctly reassigned specimens (95%), with minimal overlap between the two morphotypes (Fig. 5C). When considering all morphotype/region combinations, the LDA retained 8 PCs, resulting in an overall proportion of correctly assigned specimens of 80%. Papilliform eco-morphotypes clustered together, while molariform desert cichlids and Midas cichlids from CL Xiloá formed separate clusters, with molariforms from CL Apoyo placed in an intermediate position (Fig. S3A).

Disparity analysis revealed that the morphospace occupied by desert cichlid jaws was larger than that occupied by Midas cichlid jaws (Procrustes variance, Mexican cichlids: 0.007, Nicaraguan cichlids: 0.005; *p* < 0.05). Furthermore, each eco-morphotype utilized a distinct relative portion of the morphospace (Procrustes variance in desert cichlid molariform: 0.010, desert cichlid papilliform: 0.004, Midas cichlid molariform: 0.005, Midas cichlid papilliform: 0.006; *p* < 0.05).

To discern regional differences in LPJ shape, we separately compared molariform and papilliform LPJs from both geographical regions. In both cases, clustering analyses identified two distinct groups corresponding to each region, with less differentiation in papilliform LPJs (Figs. S1H, S1I, S2H, S2I). The degree of molarization was more pronounced in Mexico, where LPJs were generally longer, narrower, and had shorter horns compared to Nicaraguan cichlids.

### Body shape variation

Our analyses confirmed the polymorphic nature of body shape in both cichlid systems under study. We observed a size-related effect on body shape in all populations (*p* < 0.05) and consequently applied an allometric correction to all subsequent analyses. In all PCA examinations, 26 PCs were required to account for the totality of shape variation.

#### Body shape variation in the desert cichlid

Statistical analyses of morphometric data (PCA and k-means) identified two main body forms in the desert cichlid (Figs. 7, S1J, S2J). The two morphotypes were mainly distinguished by variations in body length and depth. PC1 revealed an array of shape variations, ranging from deeper benthic fish to slender limnetic fish with larger heads and more prominent snouts (Fig. 7). In the LDA, we retained the first 22 PCs, achieving a remarkably high overall proportion of correctly reassigned specimens (98%), with almost no overlap between morphotypes (Fig. 5D). Disparity analysis further indicated that limnetic fish exhibited greater morphological diversity (Procrustes variance, benthic: 0.0017, limnetic: 0.0025; *p* < 0.05).

**Figure 7.**
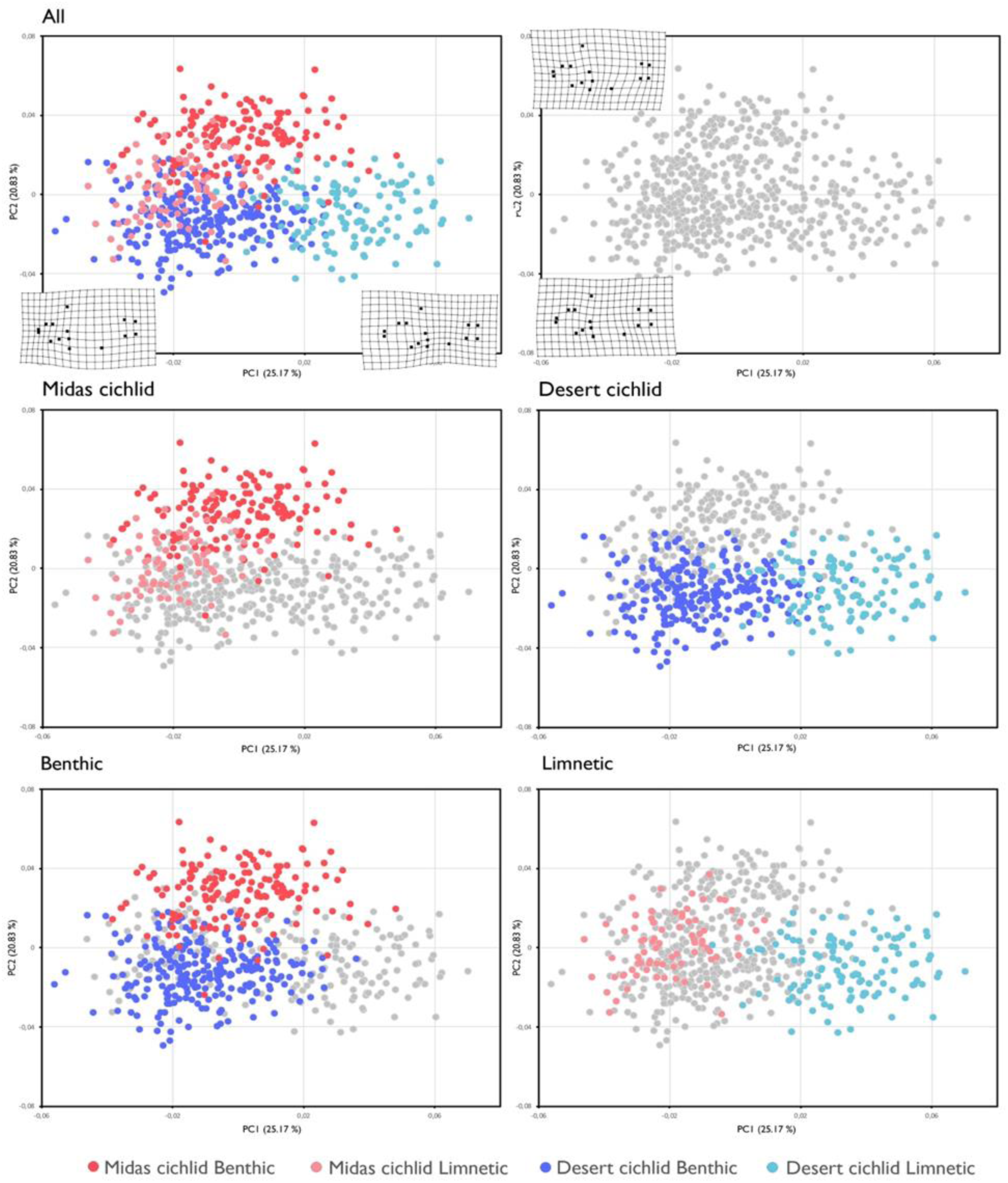
PCA showing the distribution of body shape morphospace including both geographical regions together. PC1 distinguishes two clouds of dots, one corresponding to desert cichlid limnetic and other that groups the rest of the ecotypes. In Nicaragua, Midas cichlid shows differentiation between the two morphotypes according to PC2. In Mexico, the desert cichlid shows differentiation between morphotypes according to PC1. Each morphotype differs between Mexico and Nicaragua but with the benthic type showing considerable overlap in PC1.

#### Body shape variation in the Midas cichlid

The different Midas cichlid species are known to inhabit distinct areas within the lakes, exhibiting variations in body shape. In CL Xiloá, *A. amarillo,* and *A. xiloaensis* are benthic, residing near the substrate in shallow and deeper areas, respectively, while *A. sagittae* is limnetic, occupying the water column (McKaye *et al*., 2002). Similarly, in CL Apoyo, *A. astorquii* and *A. chancho* are benthic in shallow and deeper areas, respectively, while *A. zaliosus* is limnetic (Stauffer *et al*., 2008; Elmer *et al*., 2014). The body shape analyses of all six Nicaraguan species using geometric morphometric tools aimed to compare morphotypes across lakes and to evaluate the extent of repeated evolution in body shape.

Our results confirmed the presence of two distinct body shapes in CL Xiloá, benthic (*A. amarillo* and *A. xiloaensis*) and limnetic (*A. sagittae*) fish (Figs. S1K, S2K). For the LDA, 6 PCs were retained, and the overall proportion of correctly assigned specimens was 74%, with considerable overlap between morphotypes and species (Fig. 6D). In CL Apoyo, two differentiated body shapes were also identified, benthic (*A. chancho* and *A. astorquii*) and limnetic (*A. zaliosus*) fish (Figs. S1L, S2L). For the LDA, 5 PCs were retained, and the overall proportion of correctly assigned specimens was 68%, again with considerable overlap between morphotypes and species (Fig. 6E). The morphological disparity was similar in both lakes (Procrustes variance, CL. Xiloá = 0.0016, CL. Apoyo = 0.0016, *p* > 0.05). Within CL. Xiloá, the benthic shallow *A. amarillo* was morphologically less diverse than the other two species (Procrustes variance, *A. amarillo*: 0.001, *A. sagittae*: 0.002, *A. xiloaensis*: 0.002), while within CL. Apoyo, morphological disparity was similar in all species (Procrustes variance, *A. astorquii*: 0.0014, *A zaliosus*: 0.0016, *A. chancho*: 0.0017; *p* > 0.05).

When comparing fish from both lakes, the analyses distinguished two body shape clusters (Figs. S1M, S2M). PC2 differentiated the two morphotypes within the Midas cichlid, similar to the desert cichlid, where benthic fish had higher bodies, while limnetic fish had slenderer bodies and more prominent snouts (Fig. 7). Unlike the desert cichlid, morphotypes did not differ in head size. For the LDA, the first 3 PCs were retained, and the overall proportion of correctly assigned specimens was 86%, with some overlap among morphotypes (Fig. 6F). The disparity analysis did not show significant differences in the morphospace occupied by each morphotype (Procrustes variance, benthic: 0.0016, limnetic: 0.0017; *p* < 0.05).

The comparison of benthic species across lakes showed considerable overlap, although the k-means clustering analysis recovered an optimal number of clusters of k = 2 (Figs. S1N, S2N). For the LDA, 25 PCs were retained, and the overall proportion of correctly reassigned specimens was 87%. The disparity analysis found that the morphospace occupied by species in each lake did not differ (Procrustes variance, CL Apoyo: 0.0013, CL Xiloá: 0.0014; *p* > 0.05).

When comparing the two limnetic Midas cichlid species, analyses resolved two different body shape clusters (Figs. S1O, S2O; see also Fig. 6F). For the LDA, 24 PCs were retained, and the overall proportion of correctly reassigned specimens was 99%. The disparity analysis showed no differences between the two species (Procrustes variance, CL Apoyo: 0.0010, CL Xiloá: 0.0011; *p* > 0.05).

#### Body shape variation across cichlid radiations

We conducted a comprehensive analysis of fish from two geographical regions, aiming to explore the level of convergence between morphotypes and identify any geographical differences. The PCA on two major axes (PC1 = 25.17%, PC2 = 20.83%) highlighted significant differences between limnetic fish from the two radiations, with considerable overlap among all benthic fish (Fig. 7).

With the k-means clustering, we identified three distinct clusters (k = 3): one for desert cichlid limnetic fish, another for Midas cichlid benthic fish, and a third cluster comprising a mix of desert cichlid benthic and Midas cichlid limnetic individuals (Figs. S1P, S2P). The LDA with k = 3, retaining 25 PCs, resulted in a 97% correct reassignment rate (Fig. 5F). Considering all morphotype/region combinations (6 groups), the discriminant analysis with 23 retained PCs achieved a 94% overall correct assignment rate. Specifically, desert cichlid limnetic fish differentiated from the rest, while Midas cichlid limnetic fish from both lakes overlapped with each other and with desert cichlid benthic fish. Midas cichlid benthic fish exhibited significant overlap as well (Fig. S3B).

The disparity analysis showed no differences in the morphospace occupied by desert cichlid and Midas cichlid (Procrustes variance, desert cichlid: 0.002, Midas cichlid: 0.002; *p* > 0.05), nor in the morphospace occupied by each morphotype (benthic desert cichlid: 0.002, limnetic desert cichlid: 0.003, benthic Midas cichlid: 0.002, limnetic Midas cichlid: 0.002; *p* > 0.05).

To understand regional differences in body shape, we separately compared benthic and limnetic bodies from both geographical regions. Clustering analyses in both cases revealed two differentiated groups corresponding to each region (Figs. S1Q, S1R, S2Q, S2R). Disparity analysis indicated equivalent morphospace occupied by each region (Procrustes variance, Mexico: 0.002, Nicaragua: 0.002; *p* > 0.05).

#### Modularity analyses

The analysis of modularity that was performed with geomorph supported independent modules for the head and trunk in both cichlid lineages (desert cichlid, CR = 0.91, Midas cichlid, CR = 0.92; p < 0.05).

We compared the relative length of the head and trunk of all species and morphotypes based on meristic measurements. The factorial-ANOVA revealed statistically significant differences between benthic and limnetic species in both Mexico and Nicaragua. Benthic species across systems did not differ, but limnetic species from Nicaragua and Mexico did show differences. We performed a regression analysis to understand the relationship between head and trunk. The slope of the regression followed a similar trend in Mexican individuals and benthic species from Nicaragua but limnetic species from Nicaragua showed a steeper slope.

### Gene expression

We analysed gene expression patterns in 25 samples (see Table S1 for details). A total of 11,180 genes exhibited differential expression between the desert cichlid and the Midas cichlid, regardless of their morphological characteristics. Additionally, in Nicaragua, 960 genes showed differential expression between fish sampled from CL Xiloá and those from CL Apoyo.

#### Gene expression differentiation associated with LPJ shape

In the desert cichlid, we only found three DEGs between molariform and papilliform fish (Fig. S4A, S5A). The PCA of these DEGs distinctly separated the two jaw morphotypes on PC1, explaining 91% of the variation (Fig. 8A). It is important to note that this result is based on a limited set of three genes. In contrast, the Midas cichlid exhibited 688 DEG between molariform and papilliform fish (Fig. S4B, S5B). The PCA showed the separation of the two morphotypes on PC1, which explained 41% of the variation (Fig. 8B). Interestingly, fish from the two lakes did not form distinct clusters. The comparison of fish with the same LPJ morphotype between Nicaraguan lakes revealed that 594 genes were differentially expressed between papilliform fish between the two lakes (Fig. S4C) and only 63 genes between the two molariform species (Fig. S4D). For both papilliform and molariform fish, the PCAs separated fish by the lake on PC1 (Fig. S6A, S6B). For papilliform fish, PC1 explained 57% of the variation, and for molariform fish, PC1 explained 82% of the variation.

**Figure 8.**
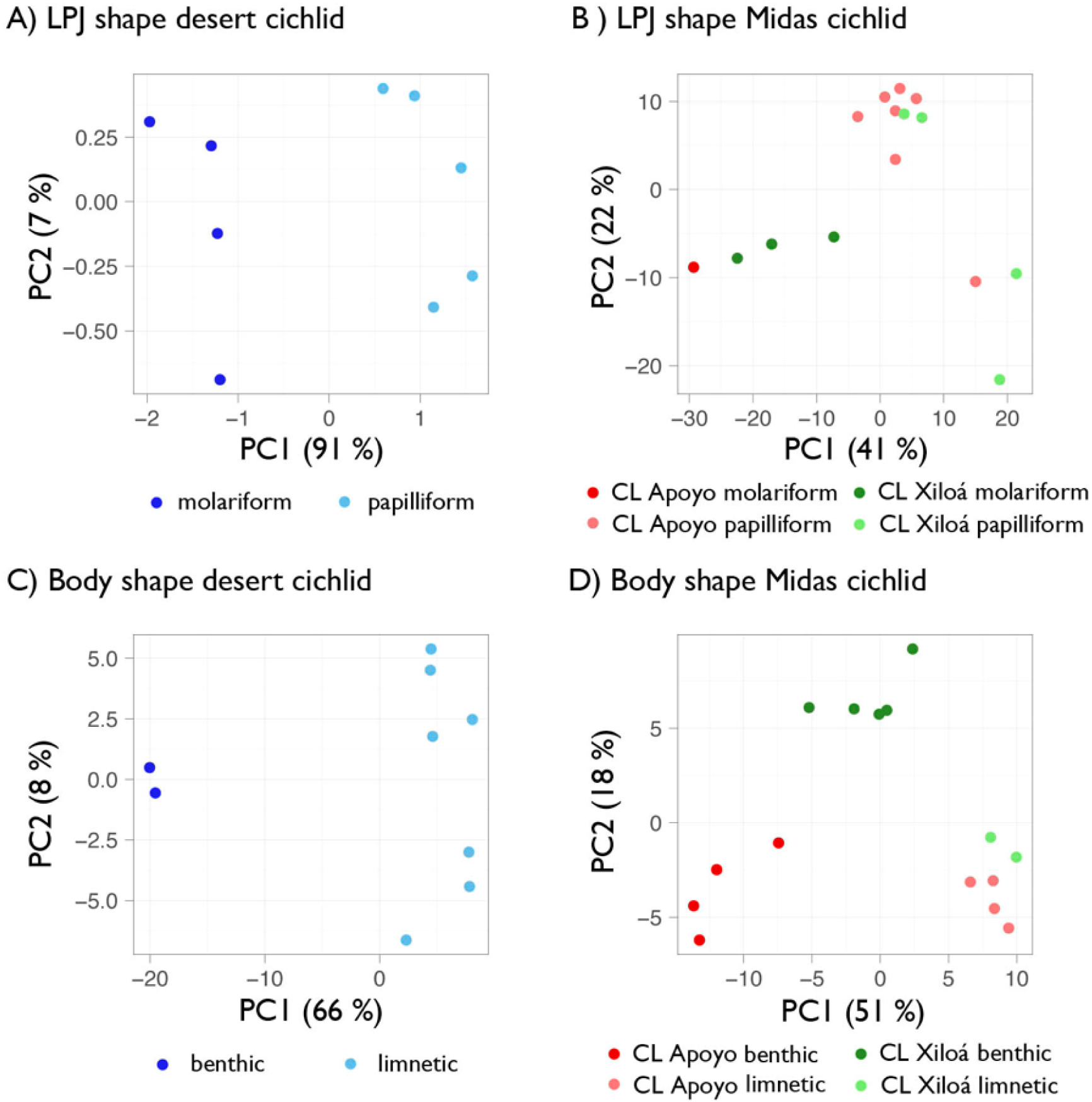
PCA of gene expression levels in the comparisons between morphotypes within each geographical region. Dots represent individuals. Axes represent the first two PCs, with the percentage of the variance explained by each of them indicated. (A) LPJ shape in the desert cichlid. (B) LPJ shape in the Midas cichlid. (C) Body shape in the desert cichlid. (D) Body shape in the Midas cichlid.

Further comparisons between geographic regions revealed 872 differentially expressed genes for papilliform fish (Fig. S4E, S5C). The PCA separated fish from both regions on PC1, albeit with low support (31% of variation; Fig. S7A), and Nicaraguan fish did not distinctly cluster by lake. Molariform fish in different geographic regions showed 481 differentially expressed genes (Fig. S4F, S5D), with PC1 separating fish from both regions (64%; Fig. S7B). Once again, Nicaraguan fish did not cluster distinctly by lake, although it is important to note that the number of individuals was very low.

For all comparisons, GO enrichment analyses indicated that differentially expressed genes were primarily involved in various metabolic and cellular processes, cell signalling, and mRNA regulation, both between papilliform and molariform fish within geographic regions and between fish with similar jaws between geographic regions and between lakes in Nicaragua. Several immune-associated processes were also enriched.

#### Gene expression differentiation associated with body shape

We identified 286 DEG between benthic and limnetic desert cichlids (Fig. S8A, S9A). The PCA separated the two morphotypes on PC1 (66%; Fig. 8C); however, the benthic morphotype was only represented by two individuals.

For the Midas cichlid, we found fewer DEG between the two morphotypes, with only 81 genes differentially expressed (Fig. S8B, S9B). The PCA separated benthic from limnetic fish on PC1 (51%; Fig. 8D). PC1 and PC2 separated benthic fish from the two lakes, and PC2 differentiated limnetic fish (18%). Differentiation was more pronounced between benthic fish. Accordingly, we found 290 DEG between benthic fish from both lakes and 151 DEG between the two limnetic species (Fig. S8C, S8D). Within benthic and limnetic species, PCAs separated fish from the two lakes on PC1 (Fig. S6C, S6D), explaining 64% and 77% of the variation, respectively. However, limnetic fish in CL. Xiloá were only represented by two samples.

The comparison of morphotypes between the desert and Midas cichlids identified 139 DEG between benthic fish and 707 DEG between limnetic fish of each geographic region (Fig. S8E, S9C, S8F, S9D). This aligns with the more pronounced shape differentiation between limnetic fish than among benthic. Benthic morphotypes in both regions separated on PC1, although it only explained 49% of the variation, and Mexican fish were only represented by two samples (Fig. S7C). Fish from the two Nicaraguan lakes were not differentiated by this analysis. Limnetic fish from the two geographic regions separated on PC1, explaining 41% of the variation (Fig. S7D). Fish from the Nicaraguan lakes separated on PC2 (13%).

Similar to LPJ differentiation, many metabolic and cellular processes were enriched between benthic and limnetic fish, and in similar body types within and between geographic regions. This was most pronounced between limnetic fish from different regions, consistent with the more pronounced shape differences. Additionally, GO terms associated with tissue structure and development were enriched in some comparisons. Within each geographic region, several enriched GO terms were associated with an immune or other defence response. In particular, limnetic fish from the two Nicaraguan lakes differed strongly in immune processes.

## Discussion

Adaptive radiations could be responsible for much of the ecological and morphological diversity we recognize. Major axes of differentiation likely govern the diversification patterns in taxon-rich groups like cichlid fish (Barluenga *et al*., 2006; Muschick *et al*., 2012; Kusche *et al*., 2014; Ronco *et al*., 2021). Different studies reveal intriguing patterns of parallel or convergent evolution in cichlid fish, where equivalent morphologies emerge in response to similar ecological adaptive peaks (Seehausen, 2006; Muschick *et al*., 2012; Ronco *et al*., 2021). Environmental factors impose limitations on available ecological niches, coupled with genetic or physiological constraints determining the range of morphologies achievable by organisms.

Repeated patterns of eco-morphological variation in cichlid fish, particularly those related to body shape and jaw morphology, demonstrate convergence within and among cichlid assemblages (Kocher *et al*., 1993; Winemiller *et al*., 1995; Rüber & Adams, 2001; Elmer & Meyer, 2011; Muschick *et al*., 2012; Brawand *et al*., 2014; Ronco *et al*., 2021). This study provides strong evidence of convergence in the shape of the lower pharyngeal jaw in two independent Neotropical cichlid radiations when adapting to exploit hard prey. The LPJ evolves along the same morphospace trajectory in the two independent Neotropical cichlid radiations. On the contrary, we found divergent evolutionary trajectories in the body shape of fish adapting to pelagic habits in the different cichlid lineages, evidencing different strategies. We generally found gene expression patterns specific of each radiation, with no evident sings of convergent genetic processes underlying similar eco-morphotypes.

### Convergence in trophic morphology

Despite the evolutionary and ecological differences between the two adaptive radiations studied, a common feature in both cases was a subset of fish specializing in feeding on snails, displaying trophic adaptations aimed at enhancing the crushing force required to break the shells of molluscs. This specialization is frequent in most cichlid adaptive radiations (e.g., Burress, 2016), and the evolution of the key innovation in the form of pharyngeal jaws is argued to have facilitated the explosive diversification of cichlid fish (Fryer & Iles, 1972; Salzburger & Meyer, 2004; Salzburger, 2018).

In the desert cichlid (*H. minckleyi*), our results agree with previous findings (Magalhaes *et al*., 2015), revealing papilliform and molariform jaw morphotypes. Within Nicaragua, specifically in CL Xiloá, we observed two distinct types of LPJ shapes, confirming that *Amphilophus amarillo* exhibits molariform jaws while *A. sagittae* and *A. xiloaensis* have papilliform jaws. In CL Apoyo, our results indicated three different groups of LPJ shapes corresponding to each of the three species, with *A. chancho* showing an intermediate shape. When comparing all species from Nicaragua together, our analysis revealed a separation between molariform and papilliform morphotypes. Interestingly, in CL Apoyo, two types of papilliform LPJ were identified, as all papilliform species clustered together without an intermediate third group.

Further analyses within and between lakes for papilliform species did not yield significant differences. However, when specifically comparing molariform species, each lake clustered together. Notably, molariform jaws of fish from CL Xiloá exhibited wider horns and a more convex upper line compared to fish from CL Apoyo. This, coupled with clustering analysis results grouping *A. astorquii* with papilliform species rather than *A. amarillo*, suggests a lower degree of molarization in fish from CL Apoyo.

A strong correspondence in adaptation for crushing was found in both systems, with molariform jaws generally having wider horns to anchor stronger muscles for crushing snails and a broader teeth area accommodating rounder and larger teeth. In contrast, papilliform jaws exhibited narrower horns and teeth in both radiations.

When combining samples from both regions, each radiation’s samples clustered together, indicating structural similarity between the two groups. However, some differences in equivalent morphotypes were observed due to geographic factors, suggesting that adaptation to crushing may vary based on phylogenetic constraints or ecological pressures in each region.

Notably, Mexican fish displayed relatively longer jaws, wider horns, and more space for teeth compared to Nicaraguan papilliform jaws, which were narrower than the equivalent Mexican morphotype. While the adaptive significance of these differences remains unclear, the degree of molarization is linked to the consumption of hard prey, such as snails. One possible explanation for the greater degree of molarization in Mexico is the desertic conditions of the Cuatro Ciénegas valley, where snails may represent a more abundant and profitable food source (Magalhaes *et al*., 2015). The clustering analysis revealed four groups that mixed samples and morphotypes from different geographical regions, confirming convergence between the two regions in LPJ shape. Moreover, it suggests that what we define as one morphotype in one region could transition into the other morphotype in the other region.

### Convergence and divergence in body shape

A common feature to many cichlid fish radiations is the evolution of pelagic forms exploiting the water column, particularly in the deep rift valley African lakes (e.g., Shaw *et al*. 2000; Munyandamutsa *et al*., 2021). This feature is common to other cichlid radiations, even in environments without clear profundal regions (Kornfield & Taylor, 1983; Magalhaes *et al.,* 2015). In both neotropical radiations studied here, littoral and pelagic forms are found, with benthic and limnetic body forms. However, a detailed study of their body plans evidences that each system has a fairly distinct body design for pelagic fish, while littoral populations remain majorly conserved in their body form. Our analyses suggest that body shape morphology has diverged along different axes of differentiation, likely depending on the different environmental conditions in which the fishes live. The Midas cichlid limnetic species exhibit a dorsal fin closer to the head, while in Mexico, the bodies are more fusiform, characterized by longer heads and more pointed snouts. Benthic fish in Nicaragua have larger heads compared to benthic forms in Mexico.

According to previous studies (McKaye *et al*., 2002; Stauffer & McKaye, 2002), *A. amarillo*, *A. xiloaensis*, *A. chancho*, and *A. astorquii* represent the benthic morphotypes in crater lakes Xiloá and Apoyo, respectively, while *A. sagittae* and *A. zaliosus* are the limnetic forms. Principal Component Analysis revealed two distinct groups corresponding to each morphotype, although there is considerable overlap. The primary difference between benthic and limnetic forms lies in body height, marked by the straight-line distance between the beginning of the dorsal and pelvic fins. Limnetic forms generally have shorter bodies than the other morphotype. Notably, our analysis did not reveal significant morphological differences between benthic morphotypes.

In CL Xiloá, the main differences among benthic morphotypes are observed in coloration (*A. amarillo* has a yellow throat) and jaw morphotype (*A. xiloaensis* is always papilliform, while *A. amarillo* is always molariform). Similarly, in CL Apoyo, the primary difference between benthic species lies in the trophic apparatus, with all *A. astorquii* individuals being molariform, and all *A. chancho* individuals being papilliform. These differences reflect alternative ecological preferences. Despite morphological similarities, individual species can be accurately distinguished.

### Gene expression patterns related to body and pharyngeal jaw morphotypes

Phenotypic diversification is linked to different genomic variants, and in recent evolutionary events often also or mostly related to gene regulation patterns. In the recent Neotropical radiations analysed here, we explored patterns of differential gene expression in different eco-morphotypes, and patterns of convergence in similar ones. We found clear differences between jaw types and body forms between radiations. We also found differential gene expression between crater lake morphotypes in the Midas cichlid. This corroborates not only morphotypes-specific gene regulation, but also regional and lake-specific gene regulation.

With the limitation of the samples compared, LPJ morphotypes showed very different gene regulation variation, with only three differentially expressed genes in the desert cichlid, while almost 700 genes were differentially expressed genes in the Midas cichlid. Although these differences could be due to the sample used and the considerably higher population densities in the Midas cichlid populations, this result could also indicate different genomic background for this equivalent morphological trait, which could be more complex in the Midas cichlid. An alternative explanation could be that fish with different jaw types in the Midas cichlid correspond to different species with other fixed genetic differences, in contrast with the polymorphism of the desert cichlid. We further evaluated each LPJ morphotype across radiations. Fish with papilliform jaws presented more differentially expressed genes than fish with molariform jaws, potentially suggesting a common genetic background for the derived molariform morphotype. Interestingly, larger genetic differences between papilliform LPJs were associated with lower morphological differentiation between radiations.

The analysis of gene expression differentiation of body types between radiations, in contrast, provided almost the opposite result. The derived limnetic body type showed more genes differentially expressed (over 300 DEG) than the benthic forms (less than 100 DEG). Also, the two morphotypes differed in fewer genes in the Midas cichlid (less than 50 DEG) than in the desert cichlid (ca. 100 DEG). This is an interesting result that could be at the root of the genetic background of a more complex trait. Body shape is likely governed by multiple genes of moderate to small effect (Gagnaire et al., 2013; De Lorenzo *et al*., 2023). In contrast, the molecular basis of pharyngeal jaw could be regulated by a few genes of major additive effect (Albertson & Kocher, 2006). Additional studies including more samples and a larger sequencing coverage could provide a better understanding of these processes.

### Conclusions

Cichlid fish adaptive radiations repeatedly show phenotypic convergence, often involving body shape and trophic structures. Here we provide strong evidence of convergence in the shape of a trophic structure, the pharyngeal jaw, in two independent Neotropical cichlid radiations, the desert and the Midas cichlids. However, we also evidence phenotypic divergence in body shape despite apparent morphological similarities, probably due to local differences in environmental conditions in each geographical setting. We investigated gene regulation similarities and differences of these convergent and divergent phenotypic traits, but found mostly divergent patterns. Our results agree with overall convergent phenotypic evolution, driven by non-convergent genetic mechanisms, although a careful examination of complex phenotypes has revealed also non-convergent phenotypic patterns.

## Acknowledgements

We are grateful to Roger Butlin, MARENA, and the Universidad Centro Americana in Nicaragua for their support. Funding was provided by Spanish Ministry of Science and Innovation (MCIN)/Spanish Research Agency (AEI) and European Regional Development Fund (ERDF) “A way to make Europe” through projects CGL2010-16103, CGL2013-42462-P, CGL2017-82986-C2-1-P to MB.

