## Supplementary File 1 for "Exploring phenotypic diversity: a comparative analysis of cichlid convergence"

Detailed list of the samples included in this chapter. Blank cells indicate that the sample was not used for the analysis referred by each column. Ben and Lim refer to benthic and limnetic body shape morphotypes respectively. Pap and Mol refer to papilliform and molariform LPJ morphotypes respectively.

| ID | System | Lake | Species | Body | Jaw |
|---|---|---|---|---|---|
| 11Meba12 | Mexico | Anteojo | *H. minckleyi* | Ben | Pap |
| 11Meba13 | Mexico | Anteojo | *H. minckleyi* | Ben | Pap |
| 11Meba14 | Mexico | Anteojo | *H. minckleyi* | Ben | Pap |
| 11Meba23 | Mexico | Anteojo | *H. minckleyi* | Ben | Pap |
| 11Meba27 | Mexico | Anteojo | *H. minckleyi* | Ben | Pap |
| 11Meba28 | Mexico | Anteojo | *H. minckleyi* | Ben | Pap |
| 11Meba30 | Mexico | Anteojo | *H. minckleyi* | Ben | Pap |
| 11Meba32 | Mexico | Anteojo | *H. minckleyi* | Ben | Pap |
| 11Meba33 | Mexico | Anteojo | *H. minckleyi* | Ben | Pap |
| 11Meba35 | Mexico | Anteojo | *H. minckleyi* | Ben | Pap |
| 11Meba40 | Mexico | Anteojo | *H. minckleyi* | Ben | Pap |
| 11Meba44 | Mexico | Anteojo | *H. minckleyi* | Ben | Pap |
| 11Meba46 | Mexico | Anteojo | *H. minckleyi* | Ben | Pap |
| 12Meda72 | Mexico | Anteojo | *H. minckleyi* | Ben | Pap |
| 12Meda73 | Mexico | Anteojo | *H. minckleyi* | Ben | Pap |
| 12Meda74 | Mexico | Anteojo | *H. minckleyi* | Ben | Pap |
| 12Meda77 | Mexico | Anteojo | *H. minckleyi* | Ben | Pap |
| 12Meda80 | Mexico | Anteojo | *H. minckleyi* | Ben | Pap |
| 12Meda81 | Mexico | Anteojo | *H. minckleyi* | Ben | Pap |
| 12Medb01 | Mexico | Anteojo | *H. minckleyi* | Ben | Pap |
| 12Medb03 | Mexico | Anteojo | *H. minckleyi* | Ben | Pap |
| 12Medb05 | Mexico | Anteojo | *H. minckleyi* | Ben | Pap |
| 12Medb12 | Mexico | Anteojo | *H. minckleyi* | Ben | Pap |
| 12Medb14 | Mexico | Anteojo | *H. minckleyi* | Ben | Pap |
| 11Meba17 | Mexico | Anteojo | *H. minckleyi* | Ben | |
| 11Meba18 | Mexico | Anteojo | *H. minckleyi* | Ben | |
| 11Meba19 | Mexico | Anteojo | *H. minckleyi* | Ben | |
| 11Meba20 | Mexico | Anteojo | *H. minckleyi* | Ben | |
| 11Meba21 | Mexico | Anteojo | *H. minckleyi* | Ben | |
| 11Meba22 | Mexico | Anteojo | *H. minckleyi* | Ben | |
| 11Meba24 | Mexico | Anteojo | *H. minckleyi* | Ben | |
| 11Meba26 | Mexico | Anteojo | *H. minckleyi* | Ben | |
| 11Meba29 | Mexico | Anteojo | *H. minckleyi* | Ben | |
| 11Meba31 | Mexico | Anteojo | *H. minckleyi* | Ben | |
| 11Meba34 | Mexico | Anteojo | *H. minckleyi* | Ben | |
| 11Meba36 | Mexico | Anteojo | *H. minckleyi* | Ben | |
| 11Meba37 | Mexico | Anteojo | *H. minckleyi* | Ben | |
| 11Meba39 | Mexico | Anteojo | *H. minckleyi* | Ben | |
| 11Meba41 | Mexico | Anteojo | *H. minckleyi* | Ben | |
| 11Meba43 | Mexico | Anteojo | *H. minckleyi* | Ben | |
| 11Meba47 | Mexico | Anteojo | *H. minckleyi* | Ben | |
| 11Meba49 | Mexico | Anteojo | *H. minckleyi* | Ben | |
| 12Meda75 | Mexico | Anteojo | *H. minckleyi* | Ben | |
| 12Meda76 | Mexico | Anteojo | *H. minckleyi* | Ben | |
| 12Meda78 | Mexico | Anteojo | *H. minckleyi* | Ben | |
| 12Meda79 | Mexico | Anteojo | *H. minckleyi* | Ben | |
| 12Medb02 | Mexico | Anteojo | *H. minckleyi* | Ben | |
| 12Medb04 | Mexico | Anteojo | *H. minckleyi* | Ben | |

| | | | | | |
|---|---|---|---|---|---|
| 12Medb07 | Mexico | Anteojo | *H. minckleyi* | Ben | |
| 12Medb08 | Mexico | Anteojo | *H. minckleyi* | Ben | |
| 12Medb09 | Mexico | Anteojo | *H. minckleyi* | Ben | |
| 12Medb10 | Mexico | Anteojo | *H. minckleyi* | Ben | |
| 12Medb13 | Mexico | Anteojo | *H. minckleyi* | Ben | |
| 12Medb15 | Mexico | Anteojo | *H. minckleyi* | Ben | |
| 12Medb16 | Mexico | Anteojo | *H. minckleyi* | Ben | |
| 12Meda71 | Mexico | Anteojo | *H. minckleyi* | Lim | Pap |
| 12Medb06 | Mexico | Anteojo | *H. minckleyi* | Lim | |
| 12Medb11 | Mexico | Anteojo | *H. minckleyi* | Lim | |
| 11Meba25 | Mexico | Anteojo | *H. minckleyi* | | Pap |
| 11Mebb20 | Mexico | Becerra | *H. minckleyi* | Ben | Pap |
| 12Meda32 | Mexico | Becerra | *H. minckleyi* | Ben | |
| 12Meda40 | Mexico | Becerra | *H. minckleyi* | Ben | |
| 12Meda60 | Mexico | Becerra | *H. minckleyi* | Ben | |
| 11Meba77 | Mexico | Becerra | *H. minckleyi* | Lim | Mol |
| 11Meba78 | Mexico | Becerra | *H. minckleyi* | Lim | Mol |
| 11Meba81 | Mexico | Becerra | *H. minckleyi* | Lim | Mol |
| 11Mebb04 | Mexico | Becerra | *H. minckleyi* | Lim | Mol |
| 11Mebb05 | Mexico | Becerra | *H. minckleyi* | Lim | Mol |
| 11Mebb07 | Mexico | Becerra | *H. minckleyi* | Lim | Mol |
| 11Mebb08 | Mexico | Becerra | *H. minckleyi* | Lim | Mol |
| 11Mebb12 | Mexico | Becerra | *H. minckleyi* | Lim | Mol |
| 11Mebb17 | Mexico | Becerra | *H. minckleyi* | Lim | Mol |
| 11Mebb23 | Mexico | Becerra | *H. minckleyi* | Lim | Mol |
| 11Mebb24 | Mexico | Becerra | *H. minckleyi* | Lim | Mol |
| 11Mebb26 | Mexico | Becerra | *H. minckleyi* | Lim | Mol |
| 11Mebb27 | Mexico | Becerra | *H. minckleyi* | Lim | Mol |
| 11Mebb28 | Mexico | Becerra | *H. minckleyi* | Lim | Mol |
| 11Mebb29 | Mexico | Becerra | *H. minckleyi* | Lim | Mol |
| 11Mebb31 | Mexico | Becerra | *H. minckleyi* | Lim | Mol |
| 11Mebb34 | Mexico | Becerra | *H. minckleyi* | Lim | Mol |
| 11Mebb35 | Mexico | Becerra | *H. minckleyi* | Lim | Mol |
| 12Meda28 | Mexico | Becerra | *H. minckleyi* | Lim | Mol |
| 12Meda31 | Mexico | Becerra | *H. minckleyi* | Lim | Mol |
| 12Meda36 | Mexico | Becerra | *H. minckleyi* | Lim | Mol |
| 12Meda42 | Mexico | Becerra | *H. minckleyi* | Lim | Mol |
| 12Meda43 | Mexico | Becerra | *H. minckleyi* | Lim | Mol |
| 12Meda44 | Mexico | Becerra | *H. minckleyi* | Lim | Mol |
| 12Meda45 | Mexico | Becerra | *H. minckleyi* | Lim | Mol |
| 12Meda47 | Mexico | Becerra | *H. minckleyi* | Lim | Mol |
| 12Meda50 | Mexico | Becerra | *H. minckleyi* | Lim | Mol |
| 12Meda51 | Mexico | Becerra | *H. minckleyi* | Lim | Mol |
| 12Meda52 | Mexico | Becerra | *H. minckleyi* | Lim | Mol |
| 12Meda53 | Mexico | Becerra | *H. minckleyi* | Lim | Mol |
| 12Meda54 | Mexico | Becerra | *H. minckleyi* | Lim | Mol |
| 12Meda55 | Mexico | Becerra | *H. minckleyi* | Lim | Mol |
| 12Meda56 | Mexico | Becerra | *H. minckleyi* | Lim | Mol |
| 12Meda58 | Mexico | Becerra | *H. minckleyi* | Lim | Mol |
| 12Meda61 | Mexico | Becerra | *H. minckleyi* | Lim | Mol |
| 12Meda62 | Mexico | Becerra | *H. minckleyi* | Lim | Mol |
| 12Meda63 | Mexico | Becerra | *H. minckleyi* | Lim | Mol |
| 11Meba79 | Mexico | Becerra | *H. minckleyi* | Lim | Pap |
| 11Meba80 | Mexico | Becerra | *H. minckleyi* | Lim | Pap |
| 11Mebb01 | Mexico | Becerra | *H. minckleyi* | Lim | Pap |
| 11Mebb02 | Mexico | Becerra | *H. minckleyi* | Lim | Pap |
| 11Mebb03 | Mexico | Becerra | *H. minckleyi* | Lim | Pap |
| 11Mebb06 | Mexico | Becerra | *H. minckleyi* | Lim | Pap |

| | | | | | |
|---|---|---|---|---|---|
| 11Mebb09 | Mexico | Becerra | *H. minckleyi* | Lim | Pap |
| 11Mebb10 | Mexico | Becerra | *H. minckleyi* | Lim | Pap |
| 11Mebb22 | Mexico | Becerra | *H. minckleyi* | Lim | Pap |
| 11Mebb25 | Mexico | Becerra | *H. minckleyi* | Lim | Pap |
| 11Mebb30 | Mexico | Becerra | *H. minckleyi* | Lim | Pap |
| 11Mebb33 | Mexico | Becerra | *H. minckleyi* | Lim | Pap |
| 11Mebb36 | Mexico | Becerra | *H. minckleyi* | Lim | Pap |
| 12Meda24 | Mexico | Becerra | *H. minckleyi* | Lim | Pap |
| 12Meda25 | Mexico | Becerra | *H. minckleyi* | Lim | Pap |
| 12Meda26 | Mexico | Becerra | *H. minckleyi* | Lim | Pap |
| 12Meda27 | Mexico | Becerra | *H. minckleyi* | Lim | Pap |
| 12Meda30 | Mexico | Becerra | *H. minckleyi* | Lim | Pap |
| 12Meda35 | Mexico | Becerra | *H. minckleyi* | Lim | Pap |
| 12Meda39 | Mexico | Becerra | *H. minckleyi* | Lim | Pap |
| 12Meda46 | Mexico | Becerra | *H. minckleyi* | Lim | Pap |
| 12Meda49 | Mexico | Becerra | *H. minckleyi* | Lim | Pap |
| 12Meda57 | Mexico | Becerra | *H. minckleyi* | Lim | Pap |
| 11Meba75 | Mexico | Becerra | *H. minckleyi* | Lim | |
| 11Meba76 | Mexico | Becerra | *H. minckleyi* | Lim | |
| 11Mebb11 | Mexico | Becerra | *H. minckleyi* | Lim | |
| 11Mebb15 | Mexico | Becerra | *H. minckleyi* | Lim | |
| 11Mebb19 | Mexico | Becerra | *H. minckleyi* | Lim | |
| 11Mebb21 | Mexico | Becerra | *H. minckleyi* | Lim | |
| 11Mebb32 | Mexico | Becerra | *H. minckleyi* | Lim | |
| 12Meda29 | Mexico | Becerra | *H. minckleyi* | Lim | |
| 12Meda33 | Mexico | Becerra | *H. minckleyi* | Lim | |
| 12Meda34 | Mexico | Becerra | *H. minckleyi* | Lim | |
| 12Meda37 | Mexico | Becerra | *H. minckleyi* | Lim | |
| 12Meda38 | Mexico | Becerra | *H. minckleyi* | Lim | |
| 12Meda48 | Mexico | Becerra | *H. minckleyi* | Lim | |
| 12Meda67 | Mexico | Becerra | *H. minckleyi* | | Mol |
| 12Meda68 | Mexico | Becerra | *H. minckleyi* | | Mol |
| 11Mebb04 | Mexico | Becerra | *H. minckleyi* | | Pap |
| 11Mebb14 | Mexico | Becerra | *H. minckleyi* | | Pap |
| 12Meda23 | Mexico | Becerra | *H. minckleyi* | | Pap |
| 12Meda41 | Mexico | Becerra | *H. minckleyi* | | Pap |
| 12Meda65 | Mexico | Becerra | *H. minckleyi* | | Pap |
| 12Meda70 | Mexico | Becerra | *H. minckleyi* | | Pap |
| 11Meba01 | Mexico | Churince | *H. minckleyi* | Ben | Mol |
| 11Meba02 | Mexico | Churince | *H. minckleyi* | Ben | Mol |
| 11Meba06 | Mexico | Churince | *H. minckleyi* | Ben | Mol |
| 11Meba07 | Mexico | Churince | *H. minckleyi* | Ben | Mol |
| 11Mebc34 | Mexico | Churince | *H. minckleyi* | Ben | Mol |
| 11Mebc37 | Mexico | Churince | *H. minckleyi* | Ben | Mol |
| 11Mebc41 | Mexico | Churince | *H. minckleyi* | Ben | Mol |
| 11Mebc42 | Mexico | Churince | *H. minckleyi* | Ben | Mol |
| 11Mebc43 | Mexico | Churince | *H. minckleyi* | Ben | Mol |
| 11Mebc44 | Mexico | Churince | *H. minckleyi* | Ben | Mol |
| 11Mebc45 | Mexico | Churince | *H. minckleyi* | Ben | Mol |
| 11Mebc48 | Mexico | Churince | *H. minckleyi* | Ben | Mol |
| 11Mebc51 | Mexico | Churince | *H. minckleyi* | Ben | Mol |
| 11Mebc52 | Mexico | Churince | *H. minckleyi* | Ben | Mol |
| 12Meda02 | Mexico | Churince | *H. minckleyi* | Ben | Mol |
| 12Meda03 | Mexico | Churince | *H. minckleyi* | Ben | Mol |
| 12Meda04 | Mexico | Churince | *H. minckleyi* | Ben | Mol |
| 12Meda06 | Mexico | Churince | *H. minckleyi* | Ben | Mol |
| 12Meda08 | Mexico | Churince | *H. minckleyi* | Ben | Mol |
| 12Meda09 | Mexico | Churince | *H. minckleyi* | Ben | Mol |

| | | | | | |
|---|---|---|---|---|---|
| 12Meda11 | Mexico | Churince | *H. minckleyi* | Ben | Mol |
| 12Meda12 | Mexico | Churince | *H. minckleyi* | Ben | Mol |
| 12Meda15 | Mexico | Churince | *H. minckleyi* | Ben | Mol |
| 12Meda17 | Mexico | Churince | *H. minckleyi* | Ben | Mol |
| 12Meda19 | Mexico | Churince | *H. minckleyi* | Ben | Mol |
| 12Meda21 | Mexico | Churince | *H. minckleyi* | Ben | Mol |
| 11Meba03 | Mexico | Churince | *H. minckleyi* | Ben | Pap |
| 11Meba05 | Mexico | Churince | *H. minckleyi* | Ben | Pap |
| 11Meba09 | Mexico | Churince | *H. minckleyi* | Ben | Pap |
| 11Meba10 | Mexico | Churince | *H. minckleyi* | Ben | Pap |
| 11Mebc38 | Mexico | Churince | *H. minckleyi* | Ben | Pap |
| 11Mebc39 | Mexico | Churince | *H. minckleyi* | Ben | Pap |
| 11Mebc46 | Mexico | Churince | *H. minckleyi* | Ben | Pap |
| 11Mebc54 | Mexico | Churince | *H. minckleyi* | Ben | Pap |
| 12Meda01 | Mexico | Churince | *H. minckleyi* | Ben | Pap |
| 12Meda05 | Mexico | Churince | *H. minckleyi* | Ben | Pap |
| 12Meda07 | Mexico | Churince | *H. minckleyi* | Ben | Pap |
| 12Meda13 | Mexico | Churince | *H. minckleyi* | Ben | Pap |
| 12Meda18 | Mexico | Churince | *H. minckleyi* | Ben | Pap |
| 12Meda22 | Mexico | Churince | *H. minckleyi* | Ben | Pap |
| 11Meba04 | Mexico | Churince | *H. minckleyi* | Ben | |
| 11Meba08 | Mexico | Churince | *H. minckleyi* | Ben | |
| 11Meba11 | Mexico | Churince | *H. minckleyi* | Ben | |
| 11Mebc35 | Mexico | Churince | *H. minckleyi* | Ben | |
| 12Meda10 | Mexico | Churince | *H. minckleyi* | Ben | |
| 12Meda16 | Mexico | Churince | *H. minckleyi* | Ben | |
| 12Meda20 | Mexico | Churince | *H. minckleyi* | Ben | |
| 11Mebc50 | Mexico | Churince | *H. minckleyi* | Lim | Mol |
| 12Meda14 | Mexico | Churince | *H. minckleyi* | Lim | Pap |
| 11Mebc49 | Mexico | Churince | *H. minckleyi* | | Mol |
| 11Mebc53 | Mexico | Churince | *H. minckleyi* | | Mol |
| 11Mebc36 | Mexico | Churince | *H. minckleyi* | | Pap |
| 11Mebb57 | Mexico | Escobedo | *H. minckleyi* | Ben | Mol |
| 12Medb17 | Mexico | Escobedo | *H. minckleyi* | Ben | Mol |
| 11Mebb56 | Mexico | Escobedo | *H. minckleyi* | Ben | Pap |
| 11Mebb62 | Mexico | Escobedo | *H. minckleyi* | Ben | Pap |
| 12Medb27 | Mexico | Escobedo | *H. minckleyi* | Ben | Pap |
| 12Medb28 | Mexico | Escobedo | *H. minckleyi* | Ben | Pap |
| 12Medb33 | Mexico | Escobedo | *H. minckleyi* | Ben | Pap |
| 12Medb39 | Mexico | Escobedo | *H. minckleyi* | Ben | Pap |
| 12Medb41 | Mexico | Escobedo | *H. minckleyi* | Ben | Pap |
| 11Mebb55 | Mexico | Escobedo | *H. minckleyi* | Ben | |
| 12Medb29 | Mexico | Escobedo | *H. minckleyi* | Ben | |
| 11Mebb58 | Mexico | Escobedo | *H. minckleyi* | Lim | Mol |
| 11Mebb59 | Mexico | Escobedo | *H. minckleyi* | Lim | Mol |
| 11Mebb60 | Mexico | Escobedo | *H. minckleyi* | Lim | Mol |
| 11Mebb61 | Mexico | Escobedo | *H. minckleyi* | Lim | Mol |
| 12Medb18 | Mexico | Escobedo | *H. minckleyi* | Lim | Mol |
| 12Medb21 | Mexico | Escobedo | *H. minckleyi* | Lim | Mol |
| 12Medb23 | Mexico | Escobedo | *H. minckleyi* | Lim | Mol |
| 12Medb25 | Mexico | Escobedo | *H. minckleyi* | Lim | Mol |
| 12Medb36 | Mexico | Escobedo | *H. minckleyi* | Lim | Mol |
| 12Medb37 | Mexico | Escobedo | *H. minckleyi* | Lim | Mol |
| 12Medb19 | Mexico | Escobedo | *H. minckleyi* | Lim | Pap |
| 12Medb22 | Mexico | Escobedo | *H. minckleyi* | Lim | Pap |
| 12Medb24 | Mexico | Escobedo | *H. minckleyi* | Lim | Pap |
| 12Medb35 | Mexico | Escobedo | *H. minckleyi* | Lim | Pap |
| 12Medb45 | Mexico | Escobedo | *H. minckleyi* | Lim | Pap |

| | | | | | |
|---|---|---|---|---|---|
| 12Medb20 | Mexico | Escobedo | *H. minckleyi* | Lim | |
| 12Medb26 | Mexico | Escobedo | *H. minckleyi* | Lim | |
| 12Medb34 | Mexico | Escobedo | *H. minckleyi* | | Mol |
| 12Medb43 | Mexico | Escobedo | *H. minckleyi* | | Mol |
| 12Medb31 | Mexico | Escobedo | *H. minckleyi* | | Pap |
| 12Medb40 | Mexico | Escobedo | *H. minckleyi* | | Pap |
| 11Mebc05 | Mexico | Garabatal River | *H. minckleyi* | Ben | Pap |
| 11Mebc07 | Mexico | Garabatal River | *H. minckleyi* | Ben | Pap |
| 11Mebc08 | Mexico | Garabatal River | *H. minckleyi* | Ben | Pap |
| 11Mebc03 | Mexico | Garabatal River | *H. minckleyi* | Ben | |
| 11Mebc04 | Mexico | Garabatal River | *H. minckleyi* | Ben | |
| 11Mebc06 | Mexico | Garabatal River | *H. minckleyi* | Ben | |
| 11Mebc09 | Mexico | Garabatal River | *H. minckleyi* | Lim | Mol |
| 11Mebd09 | Mexico | Juan Santos | *H. minckleyi* | Ben | Mol |
| 11Mebd12 | Mexico | Juan Santos | *H. minckleyi* | Ben | Mol |
| 11Mebd16 | Mexico | Juan Santos | *H. minckleyi* | Ben | Mol |
| 11Mebd17 | Mexico | Juan Santos | *H. minckleyi* | Ben | Mol |
| 12Medc63 | Mexico | Juan Santos | *H. minckleyi* | Ben | Mol |
| 12Medc65 | Mexico | Juan Santos | *H. minckleyi* | Ben | Mol |
| 12Medc66 | Mexico | Juan Santos | *H. minckleyi* | Ben | Mol |
| 12Medc69 | Mexico | Juan Santos | *H. minckleyi* | Ben | Mol |
| 12Medc71 | Mexico | Juan Santos | *H. minckleyi* | Ben | Mol |
| 11Mebd13 | Mexico | Juan Santos | *H. minckleyi* | Ben | Pap |
| 12Medc61 | Mexico | Juan Santos | *H. minckleyi* | Ben | Pap |
| 12Medc64 | Mexico | Juan Santos | *H. minckleyi* | Ben | Pap |
| 12Medc67 | Mexico | Juan Santos | *H. minckleyi* | Ben | Pap |
| 12Medc70 | Mexico | Juan Santos | *H. minckleyi* | Ben | Pap |
| 11Mebd15 | Mexico | Juan Santos | *H. minckleyi* | Ben | |
| 11Mebd19 | Mexico | Juan Santos | *H. minckleyi* | Ben | |
| 11Medb11 | Mexico | Juan Santos | *H. minckleyi* | Ben | |
| 11Medb18 | Mexico | Juan Santos | *H. minckleyi* | Ben | |
| 12Medc60 | Mexico | Juan Santos | *H. minckleyi* | Ben | |
| 12Medc62 | Mexico | Juan Santos | *H. minckleyi* | Ben | |
| 11Mebd10 | Mexico | Juan Santos | *H. minckleyi* | | Mol |
| 11Mebd14 | Mexico | Juan Santos | *H. minckleyi* | | Mol |
| 11Mebd18 | Mexico | Juan Santos | *H. minckleyi* | | Mol |
| 11Mebd26 | Mexico | Juan Santos | *H. minckleyi* | | Mol |
| 11Mebd27 | Mexico | Juan Santos | *H. minckleyi* | | Mol |
| 11Mebd33 | Mexico | Juan Santos | *H. minckleyi* | | Mol |
| 11Mebd34 | Mexico | Juan Santos | *H. minckleyi* | | Mol |
| 11Mebd36 | Mexico | Juan Santos | *H. minckleyi* | | Mol |
| 11Mebd37 | Mexico | Juan Santos | *H. minckleyi* | | Mol |
| 11Mebd38 | Mexico | Juan Santos | *H. minckleyi* | | Mol |
| 11Mebd39 | Mexico | Juan Santos | *H. minckleyi* | | Mol |
| 11Mebd40 | Mexico | Juan Santos | *H. minckleyi* | | Mol |
| 11Mebd41 | Mexico | Juan Santos | *H. minckleyi* | | Mol |
| 11Mebd42 | Mexico | Juan Santos | *H. minckleyi* | | Mol |
| 12Medc68 | Mexico | Juan Santos | *H. minckleyi* | | Mol |
| 11Mebd11 | Mexico | Juan Santos | *H. minckleyi* | | Pap |
| 11Mebd20 | Mexico | Juan Santos | *H. minckleyi* | | Pap |
| 11Mebd22 | Mexico | Juan Santos | *H. minckleyi* | | Pap |
| 11Mebd29 | Mexico | Juan Santos | *H. minckleyi* | | Pap |
| 11Mebd31 | Mexico | Juan Santos | *H. minckleyi* | | Pap |
| 11Mebd44 | Mexico | Juan Santos | *H. minckleyi* | | Pap |
| 11Mebc56 | Mexico | Mesquítez River | *H. minckleyi* | Ben | Mol |
| 11Mebc58 | Mexico | Mesquítez River | *H. minckleyi* | Ben | Mol |
| 11Mebc62 | Mexico | Mesquítez River | *H. minckleyi* | Ben | Mol |
| 11Mebc63 | Mexico | Mesquítez River | *H. minckleyi* | Ben | Mol |

| | | | | | |
|---|---|---|---|---|---|
| 11Mebc64 | Mexico | Mesquítez River | *H. minckleyi* | Ben | Mol |
| 11Mebc66 | Mexico | Mesquítez River | *H. minckleyi* | Ben | Mol |
| 11Mebc68 | Mexico | Mesquítez River | *H. minckleyi* | Ben | Mol |
| 11Mebc69 | Mexico | Mesquítez River | *H. minckleyi* | Ben | Mol |
| 11Mebc71 | Mexico | Mesquítez River | *H. minckleyi* | Ben | Mol |
| 11Mebc72 | Mexico | Mesquítez River | *H. minckleyi* | Ben | Mol |
| 11Mebc80 | Mexico | Mesquítez River | *H. minckleyi* | Ben | Mol |
| 11Mebd03 | Mexico | Mesquítez River | *H. minckleyi* | Ben | Mol |
| 11Mebc65 | Mexico | Mesquítez River | *H. minckleyi* | Ben | Pap |
| 11Mebc67 | Mexico | Mesquítez River | *H. minckleyi* | Ben | Pap |
| 11Mebc73 | Mexico | Mesquítez River | *H. minckleyi* | Ben | Pap |
| 11Mebc74 | Mexico | Mesquítez River | *H. minckleyi* | Ben | Pap |
| 11Mebc75 | Mexico | Mesquítez River | *H. minckleyi* | Ben | Pap |
| 11Mebc77 | Mexico | Mesquítez River | *H. minckleyi* | Ben | Pap |
| 11Mebc79 | Mexico | Mesquítez River | *H. minckleyi* | Ben | Pap |
| 11Mebc81 | Mexico | Mesquítez River | *H. minckleyi* | Ben | Pap |
| 11Mebd04 | Mexico | Mesquítez River | *H. minckleyi* | Ben | Pap |
| 11Mebd05 | Mexico | Mesquítez River | *H. minckleyi* | Ben | Pap |
| 11Mebc55 | Mexico | Mesquítez River | *H. minckleyi* | Ben | |
| 11Mebc59 | Mexico | Mesquítez River | *H. minckleyi* | Ben | |
| 11Mebc70 | Mexico | Mesquítez River | *H. minckleyi* | Ben | |
| 11Mebc76 | Mexico | Mesquítez River | *H. minckleyi* | Ben | |
| 11Mebc78 | Mexico | Mesquítez River | *H. minckleyi* | Ben | |
| 11Mebd01 | Mexico | Mesquítez River | *H. minckleyi* | Ben | |
| 11Mebd02 | Mexico | Mesquítez River | *H. minckleyi* | Ben | |
| 12Medd17 | Mexico | Mesquítez River | *H. minckleyi* | | Mol |
| 12Medd18 | Mexico | Mesquítez River | *H. minckleyi* | | Mol |
| 12Medd20 | Mexico | Mesquítez River | *H. minckleyi* | | Mol |
| 12Medd21 | Mexico | Mesquítez River | *H. minckleyi* | | Mol |
| 12Medd22 | Mexico | Mesquítez River | *H. minckleyi* | | Mol |
| 12Medd24 | Mexico | Mesquítez River | *H. minckleyi* | | Mol |
| 12Medd27 | Mexico | Mesquítez River | *H. minckleyi* | | Mol |
| 12Medd28 | Mexico | Mesquítez River | *H. minckleyi* | | Mol |
| 12Medd34 | Mexico | Mesquítez River | *H. minckleyi* | | Mol |
| 12Medd36 | Mexico | Mesquítez River | *H. minckleyi* | | Mol |
| 12Medd16 | Mexico | Mesquítez River | *H. minckleyi* | | Pap |
| 12Medd23 | Mexico | Mesquítez River | *H. minckleyi* | | Pap |
| 12Medd25 | Mexico | Mesquítez River | *H. minckleyi* | | Pap |
| 12Medd30 | Mexico | Mesquítez River | *H. minckleyi* | | Pap |
| 12Medd31 | Mexico | Mesquítez River | *H. minckleyi* | | Pap |
| 12Medd33 | Mexico | Mesquítez River | *H. minckleyi* | | Pap |
| 12Medd37 | Mexico | Mesquítez River | *H. minckleyi* | | Pap |
| 12Medc06 | Mexico | Mojarral Este | *H. minckleyi* | Ben | Mol |
| 12Medc07 | Mexico | Mojarral Este | *H. minckleyi* | Ben | Mol |
| 12Medc08 | Mexico | Mojarral Este | *H. minckleyi* | Ben | Mol |
| 12Medc09 | Mexico | Mojarral Este | *H. minckleyi* | Ben | Mol |
| 12Medc12 | Mexico | Mojarral Este | *H. minckleyi* | Ben | Mol |
| 12Medc13 | Mexico | Mojarral Este | *H. minckleyi* | Ben | Mol |
| 12Medc14 | Mexico | Mojarral Este | *H. minckleyi* | Ben | Mol |
| 12Medc18 | Mexico | Mojarral Este | *H. minckleyi* | Ben | Mol |
| 12Medc22 | Mexico | Mojarral Este | *H. minckleyi* | Ben | Mol |
| 12Medc23 | Mexico | Mojarral Este | *H. minckleyi* | Ben | Mol |
| 12Medc26 | Mexico | Mojarral Este | *H. minckleyi* | Ben | Mol |
| 12Medc33 | Mexico | Mojarral Este | *H. minckleyi* | Ben | Mol |
| 12Medc37 | Mexico | Mojarral Este | *H. minckleyi* | Ben | Mol |
| 12Medc41 | Mexico | Mojarral Este | *H. minckleyi* | Ben | Mol |
| 12Medc15 | Mexico | Mojarral Este | *H. minckleyi* | Ben | Pap |
| 12Medc16 | Mexico | Mojarral Este | *H. minckleyi* | Ben | Pap |

| | | | | | |
|---|---|---|---|---|---|
| 12Medc27 | Mexico | Mojarral Este | *H. minckleyi* | Ben | Pap |
| 12Medc29 | Mexico | Mojarral Este | *H. minckleyi* | Ben | Pap |
| 12Medc35 | Mexico | Mojarral Este | *H. minckleyi* | Ben | Pap |
| 12Medc11 | Mexico | Mojarral Este | *H. minckleyi* | Ben | |
| 12Medc19 | Mexico | Mojarral Este | *H. minckleyi* | Ben | |
| 12Medc24 | Mexico | Mojarral Este | *H. minckleyi* | Ben | |
| 12Medc34 | Mexico | Mojarral Este | *H. minckleyi* | Ben | |
| 12Medc39 | Mexico | Mojarral Este | *H. minckleyi* | Ben | |
| 12Medc40 | Mexico | Mojarral Este | *H. minckleyi* | Ben | |
| 12Medc10 | Mexico | Mojarral Este | *H. minckleyi* | Lim | Mol |
| 12Medc20 | Mexico | Mojarral Este | *H. minckleyi* | | Mol |
| 12Medc28 | Mexico | Mojarral Este | *H. minckleyi* | | Mol |
| 12Medc21 | Mexico | Mojarral Este | *H. minckleyi* | | Pap |
| 12Medc31 | Mexico | Mojarral Este | *H. minckleyi* | | Pap |
| 12Medc32 | Mexico | Mojarral Este | *H. minckleyi* | | Pap |
| 12Medc36 | Mexico | Mojarral Este | *H. minckleyi* | | Pap |
| 12Medc38 | Mexico | Mojarral Este | *H. minckleyi* | | Pap |
| 12Medc76 | Mexico | Mojarral Oeste | *H. minckleyi* | Ben | Mol |
| 12Medb71 | Mexico | Mojarral Oeste | *H. minckleyi* | Ben | |
| 12Medb73 | Mexico | Mojarral Oeste | *H. minckleyi* | Ben | |
| 12Medd02 | Mexico | Mojarral Oeste | *H. minckleyi* | Ben | |
| 12Medd04 | Mexico | Mojarral Oeste | *H. minckleyi* | Lim | Mol |
| 12Medd05 | Mexico | Mojarral Oeste | *H. minckleyi* | Lim | Mol |
| 12Medd08 | Mexico | Mojarral Oeste | *H. minckleyi* | Lim | Mol |
| 12Medd14 | Mexico | Mojarral Oeste | *H. minckleyi* | Lim | Mol |
| 12Medc72 | Mexico | Mojarral Oeste | *H. minckleyi* | Lim | Pap |
| 12Medc74 | Mexico | Mojarral Oeste | *H. minckleyi* | Lim | Pap |
| 12Medc81 | Mexico | Mojarral Oeste | *H. minckleyi* | Lim | Pap |
| 12Medc80 | Mexico | Mojarral Oeste | *H. minckleyi* | Lim | |
| 12Medd10 | Mexico | Mojarral Oeste | *H. minckleyi* | Lim | |
| 12Medc75 | Mexico | Mojarral Oeste | *H. minckleyi* | | Mol |
| 12Medc77 | Mexico | Mojarral Oeste | *H. minckleyi* | | Mol |
| 12Medd09 | Mexico | Mojarral Oeste | *H. minckleyi* | | Mol |
| 12Medc78 | Mexico | Mojarral Oeste | *H. minckleyi* | | Pap |
| 12Medc79 | Mexico | Mojarral Oeste | *H. minckleyi* | | Pap |
| 12Medd07 | Mexico | Mojarral Oeste | *H. minckleyi* | | Pap |
| 12Medd11 | Mexico | Mojarral Oeste | *H. minckleyi* | | Pap |
| 12Medd12 | Mexico | Mojarral Oeste | *H. minckleyi* | | Pap |
| 12Medd13 | Mexico | Mojarral Oeste | *H. minckleyi* | | Pap |
| 12Medd15 | Mexico | Mojarral Oeste | *H. minckleyi* | | Pap |
| 12Medb77 | Mexico | Pozas Azules | *H. minckleyi* | Ben | Mol |
| 12Medb79 | Mexico | Pozas Azules | *H. minckleyi* | Ben | Mol |
| 12Medc42 | Mexico | Pozas Azules | *H. minckleyi* | Ben | Mol |
| 12Medc43 | Mexico | Pozas Azules | *H. minckleyi* | Ben | Mol |
| 12Medc46 | Mexico | Pozas Azules | *H. minckleyi* | Ben | Mol |
| 12Medc59 | Mexico | Pozas Azules | *H. minckleyi* | Ben | Mol |
| 11Mebb38 | Mexico | Pozas Azules | *H. minckleyi* | Ben | Pap |
| 11Mebb39 | Mexico | Pozas Azules | *H. minckleyi* | Ben | Pap |
| 11Mebb42 | Mexico | Pozas Azules | *H. minckleyi* | Ben | Pap |
| 11Mebb43 | Mexico | Pozas Azules | *H. minckleyi* | Ben | Pap |
| 12Medb70 | Mexico | Pozas Azules | *H. minckleyi* | Ben | Pap |
| 12Medb75 | Mexico | Pozas Azules | *H. minckleyi* | Ben | Pap |
| 12Medc50 | Mexico | Pozas Azules | *H. minckleyi* | Ben | Pap |
| 12Medc52 | Mexico | Pozas Azules | *H. minckleyi* | Ben | Pap |
| 12Medc54 | Mexico | Pozas Azules | *H. minckleyi* | Ben | Pap |
| 11Mebb40 | Mexico | Pozas Azules | *H. minckleyi* | Ben | |
| 12Medb66 | Mexico | Pozas Azules | *H. minckleyi* | Ben | |
| 12Medb68 | Mexico | Pozas Azules | *H. minckleyi* | Ben | |

| | | | | | |
|---|---|---|---|---|---|
| 12Medb74 | Mexico | Pozas Azules | *H. minckleyi* | Ben | |
| 12Medb80 | Mexico | Pozas Azules | *H. minckleyi* | Ben | |
| 12Medc01 | Mexico | Pozas Azules | *H. minckleyi* | Ben | |
| 12Medc44 | Mexico | Pozas Azules | *H. minckleyi* | Ben | |
| 11Mebb41 | Mexico | Pozas Azules | *H. minckleyi* | Lim | Mol |
| 12Medc48 | Mexico | Pozas Azules | *H. minckleyi* | Lim | Mol |
| 12Medc03 | Mexico | Pozas Azules | *H. minckleyi* | Lim | Pap |
| 12Medc55 | Mexico | Pozas Azules | *H. minckleyi* | Lim | Pap |
| 12Medc56 | Mexico | Pozas Azules | *H. minckleyi* | Lim | Pap |
| 12Medc58 | Mexico | Pozas Azules | *H. minckleyi* | Lim | |
| 12Medb67 | Mexico | Pozas Azules | *H. minckleyi* | | Mol |
| 12Medb69 | Mexico | Pozas Azules | *H. minckleyi* | | Mol |
| 12Medc45 | Mexico | Pozas Azules | *H. minckleyi* | | Mol |
| 12Medc47 | Mexico | Pozas Azules | *H. minckleyi* | | Mol |
| 12Medc51 | Mexico | Pozas Azules | *H. minckleyi* | | Mol |
| 12Medc53 | Mexico | Pozas Azules | *H. minckleyi* | | Mol |
| 12Medc02 | Mexico | Pozas Azules | *H. minckleyi* | | Pap |
| 12Medc04 | Mexico | Pozas Azules | *H. minckleyi* | | Pap |
| 12Medc05 | Mexico | Pozas Azules | *H. minckleyi* | | Pap |
| 12Medc57 | Mexico | Pozas Azules | *H. minckleyi* | | Pap |
| 11Mebc30 | Mexico | Río Garabatal | *H. minckleyi* | Ben | |
| 11Mebc25 | Mexico | Río Garabatal | *H. minckleyi* | Lim | Mol |
| 11Mebc26 | Mexico | Río Garabatal | *H. minckleyi* | | Mol |
| 11Mebc27 | Mexico | Río Garabatal | *H. minckleyi* | | Pap |
| 11Meba69 | Mexico | Santa Tecla | *H. minckleyi* | | Pap |
| 11Meba70 | Mexico | Santa Tecla | *H. minckleyi* | | Pap |
| 11Mebc12 | Mexico | Tierras Blancas | *H. minckleyi* | Ben | Mol |
| 11Mebc10 | Mexico | Tierras Blancas | *H. minckleyi* | Ben | Pap |
| 11Mebc13 | Mexico | Tierras Blancas | *H. minckleyi* | Ben | Pap |
| 11Mebc23 | Mexico | Tierras Blancas | *H. minckleyi* | Ben | Pap |
| 11Mebc18 | Mexico | Tierras Blancas | *H. minckleyi* | Ben | |
| 11Mebc19 | Mexico | Tierras Blancas | *H. minckleyi* | Ben | |
| 11Mebc24 | Mexico | Tierras Blancas | *H. minckleyi* | Ben | |
| 11Mebc16 | Mexico | Tierras Blancas | *H. minckleyi* | Lim | Mol |
| 11Mebc11 | Mexico | Tierras Blancas | *H. minckleyi* | | Pap |
| 11Mebb63 | Mexico | Tío Cándido | *H. minckleyi* | Ben | Mol |
| 11Mebb67 | Mexico | Tío Cándido | *H. minckleyi* | Ben | Mol |
| 11Mebb69 | Mexico | Tío Cándido | *H. minckleyi* | Ben | Mol |
| 11Mebb79 | Mexico | Tío Cándido | *H. minckleyi* | Ben | Mol |
| 11Mebb81 | Mexico | Tío Cándido | *H. minckleyi* | Ben | Mol |
| 12Medb47 | Mexico | Tío Cándido | *H. minckleyi* | Ben | Mol |
| 12Medb55 | Mexico | Tío Cándido | *H. minckleyi* | Ben | Mol |
| 12Medb57 | Mexico | Tío Cándido | *H. minckleyi* | Ben | Mol |
| 12Medb65 | Mexico | Tío Cándido | *H. minckleyi* | Ben | Mol |
| 11Mebb70 | Mexico | Tío Cándido | *H. minckleyi* | Ben | Pap |
| 11Mebb71 | Mexico | Tío Cándido | *H. minckleyi* | Ben | Pap |
| 11Mebb73 | Mexico | Tío Cándido | *H. minckleyi* | Ben | Pap |
| 11Mebb77 | Mexico | Tío Cándido | *H. minckleyi* | Ben | Pap |
| 12Medb48 | Mexico | Tío Cándido | *H. minckleyi* | Ben | Pap |
| 12Medb52 | Mexico | Tío Cándido | *H. minckleyi* | Ben | Pap |
| 12Medb60 | Mexico | Tío Cándido | *H. minckleyi* | Ben | Pap |
| 11Mebb68 | Mexico | Tío Cándido | *H. minckleyi* | Ben | |
| 11Mebb76 | Mexico | Tío Cándido | *H. minckleyi* | Ben | |
| 12Medb50 | Mexico | Tío Cándido | *H. minckleyi* | Ben | |
| 12Medb58 | Mexico | Tío Cándido | *H. minckleyi* | Ben | |
| 11Mebb80 | Mexico | Tío Cándido | *H. minckleyi* | Lim | Mol |
| 11Mebb66 | Mexico | Tío Cándido | *H. minckleyi* | Lim | Pap |
| 11Mebb65 | Mexico | Tío Cándido | *H. minckleyi* | Lim | |

| | | | | | |
|---|---|---|---|---|---|
| 11Mebb78 | Mexico | Tío Cándido | *H. minckleyi* | Lim | |
| 11Mebb75 | Mexico | Tío Cándido | *H. minckleyi* | | Mol |
| 12Medb49 | Mexico | Tío Cándido | *H. minckleyi* | | Mol |
| 12Medb51 | Mexico | Tío Cándido | *H. minckleyi* | | Mol |
| 12Medb61 | Mexico | Tío Cándido | *H. minckleyi* | | Mol |
| 12Medb63 | Mexico | Tío Cándido | *H. minckleyi* | | Mol |
| 11Mebb64 | Mexico | Tío Cándido | *H. minckleyi* | | Pap |
| 12Medb53 | Mexico | Tío Cándido | *H. minckleyi* | | Pap |
| 12Medb59 | Mexico | Tío Cándido | *H. minckleyi* | | Pap |
| 12Medb62 | Mexico | Tío Cándido | *H. minckleyi* | | Pap |
| 10NIag02 | Nicaragua | Apoyo | *A. astorquii* | Ben | Mol |
| 10NIag37 | Nicaragua | Apoyo | *A. astorquii* | Ben | Mol |
| 10NIak05 | Nicaragua | Apoyo | *A. astorquii* | Ben | Mol |
| 11NIce66 | Nicaragua | Apoyo | *A. astorquii* | Ben | Mol |
| 11NIce72 | Nicaragua | Apoyo | *A. astorquii* | Ben | Mol |
| 11NIce73 | Nicaragua | Apoyo | *A. astorquii* | Ben | Mol |
| 11NIce74 | Nicaragua | Apoyo | *A. astorquii* | Ben | Mol |
| 11NIcg03 | Nicaragua | Apoyo | *A. astorquii* | Ben | Mol |
| 14NIgc68 | Nicaragua | Apoyo | *A. astorquii* | Ben | Mol |
| 14NIgd58 | Nicaragua | Apoyo | *A. astorquii* | Ben | Mol |
| 14NIgd59 | Nicaragua | Apoyo | *A. astorquii* | Ben | Mol |
| 14NIgd60 | Nicaragua | Apoyo | *A. astorquii* | Ben | Mol |
| 14NIgd62 | Nicaragua | Apoyo | *A. astorquii* | Ben | Mol |
| 14NIgd63 | Nicaragua | Apoyo | *A. astorquii* | Ben | Mol |
| 16NIhc05 | Nicaragua | Apoyo | *A. astorquii* | Ben | Mol |
| 16NIhc07 | Nicaragua | Apoyo | *A. astorquii* | Ben | Mol |
| 16NIhc09 | Nicaragua | Apoyo | *A. astorquii* | Ben | Mol |
| 16NIhc11 | Nicaragua | Apoyo | *A. astorquii* | Ben | Mol |
| 16NIhc13 | Nicaragua | Apoyo | *A. astorquii* | Ben | Mol |
| 16NIhc15 | Nicaragua | Apoyo | *A. astorquii* | Ben | Mol |
| 16NIhc17 | Nicaragua | Apoyo | *A. astorquii* | Ben | Mol |
| 16NIhc19 | Nicaragua | Apoyo | *A. astorquii* | Ben | Mol |
| 16NIhc21 | Nicaragua | Apoyo | *A. astorquii* | Ben | Mol |
| 16NIhc23 | Nicaragua | Apoyo | *A. astorquii* | Ben | Mol |
| 16NIhc25 | Nicaragua | Apoyo | *A. astorquii* | Ben | Mol |
| 16NIhc27 | Nicaragua | Apoyo | *A. astorquii* | Ben | Mol |
| 16NIhc29 | Nicaragua | Apoyo | *A. astorquii* | Ben | Mol |
| 16NIhc31 | Nicaragua | Apoyo | *A. astorquii* | Ben | Mol |
| 16NIhc33 | Nicaragua | Apoyo | *A. astorquii* | Ben | Mol |
| 11NIcf80 | Nicaragua | Apoyo | *A. astorquii* | Ben | Pap |
| 09NI25f4 | Nicaragua | Apoyo | *A. astorquii* | Ben | |
| 09NI26e4 | Nicaragua | Apoyo | *A. astorquii* | Ben | |
| 09NI26f9 | Nicaragua | Apoyo | *A. astorquii* | Ben | |
| 10NIag16 | Nicaragua | Apoyo | *A. astorquii* | Ben | |
| 10NIag29 | Nicaragua | Apoyo | *A. astorquii* | Ben | |
| 10NIag30 | Nicaragua | Apoyo | *A. astorquii* | Ben | |
| 10NIag35 | Nicaragua | Apoyo | *A. astorquii* | Ben | |
| 10NIag38 | Nicaragua | Apoyo | *A. astorquii* | Ben | |
| 10NIag39 | Nicaragua | Apoyo | *A. astorquii* | Ben | |
| 10NIag40 | Nicaragua | Apoyo | *A. astorquii* | Ben | |
| 10NIag51 | Nicaragua | Apoyo | *A. astorquii* | Ben | |
| 11NIce67 | Nicaragua | Apoyo | *A. astorquii* | Ben | |
| 11NIce71 | Nicaragua | Apoyo | *A. astorquii* | Ben | |
| 11NIcg10 | Nicaragua | Apoyo | *A. astorquii* | Ben | |
| 11NIcg12 | Nicaragua | Apoyo | *A. astorquii* | Ben | |
| 11NIcg13 | Nicaragua | Apoyo | *A. astorquii* | Ben | |
| 11NIcg15 | Nicaragua | Apoyo | *A. astorquii* | Ben | |
| 11NIcg24 | Nicaragua | Apoyo | *A. astorquii* | Ben | |

| | | | | | |
|---|---|---|---|---|---|
| 13NIec47 | Nicaragua | Apoyo | *A. astorquii* | Ben | |
| 13NIec71 | Nicaragua | Apoyo | *A. astorquii* | Ben | |
| 13NIed02 | Nicaragua | Apoyo | *A. astorquii* | Ben | |
| 14NIgd66 | Nicaragua | Apoyo | *A. astorquii* | Ben | |
| 14NIgd67 | Nicaragua | Apoyo | *A. astorquii* | Ben | |
| 14NIgd68 | Nicaragua | Apoyo | *A. astorquii* | Ben | |
| 10NIag31 | Nicaragua | Apoyo | *A. astorquii* | | Mol |
| 10NIag42 | Nicaragua | Apoyo | *A. astorquii* | | Mol |
| 10NIag56 | Nicaragua | Apoyo | *A. astorquii* | | Mol |
| 10NIag60 | Nicaragua | Apoyo | *A. astorquii* | | Mol |
| 10NIag62 | Nicaragua | Apoyo | *A. astorquii* | | Mol |
| 10NIag64 | Nicaragua | Apoyo | *A. astorquii* | | Mol |
| 10NIag67 | Nicaragua | Apoyo | *A. astorquii* | | Mol |
| 10NIag78 | Nicaragua | Apoyo | *A. astorquii* | | Mol |
| 10NIak01 | Nicaragua | Apoyo | *A. astorquii* | | Mol |
| 10NIak04 | Nicaragua | Apoyo | *A. astorquii* | | Mol |
| 10NIak07 | Nicaragua | Apoyo | *A. astorquii* | | Mol |
| 10NIak08 | Nicaragua | Apoyo | *A. astorquii* | | Mol |
| 10NIak10 | Nicaragua | Apoyo | *A. astorquii* | | Mol |
| 10NIak14 | Nicaragua | Apoyo | *A. astorquii* | | Mol |
| 11NIce76 | Nicaragua | Apoyo | *A. astorquii* | | Mol |
| 11NIcg09 | Nicaragua | Apoyo | *A. astorquii* | | Mol |
| 13NIec77 | Nicaragua | Apoyo | *A. astorquii* | | Mol |
| 13NIec78 | Nicaragua | Apoyo | *A. astorquii* | | Mol |
| 13NIed25 | Nicaragua | Apoyo | *A. astorquii* | | Mol |
| 13NIed29 | Nicaragua | Apoyo | *A. astorquii* | | Mol |
| 13NIed33 | Nicaragua | Apoyo | *A. astorquii* | | Mol |
| 13NIed35 | Nicaragua | Apoyo | *A. astorquii* | | Mol |
| 14NIgc65 | Nicaragua | Apoyo | *A. astorquii* | | Mol |
| 14NIgc66 | Nicaragua | Apoyo | *A. astorquii* | | Mol |
| 14NIgd28 | Nicaragua | Apoyo | *A. astorquii* | | Mol |
| 14NIgd57 | Nicaragua | Apoyo | *A. astorquii* | | Mol |
| 14NIgd61 | Nicaragua | Apoyo | *A. astorquii* | | Mol |
| 16NIhc43 | Nicaragua | Apoyo | *A. astorquii* | | Mol |
| 10NIag18 | Nicaragua | Apoyo | *A. chancho* | Ben | Pap |
| 10NIag20 | Nicaragua | Apoyo | *A. chancho* | Ben | Pap |
| 10NIag45 | Nicaragua | Apoyo | *A. chancho* | Ben | Pap |
| 10NIag46 | Nicaragua | Apoyo | *A. chancho* | Ben | Pap |
| 10NIag47 | Nicaragua | Apoyo | *A. chancho* | Ben | Pap |
| 10NIak02 | Nicaragua | Apoyo | *A. chancho* | Ben | Pap |
| 10NIak31 | Nicaragua | Apoyo | *A. chancho* | Ben | Pap |
| 11NIce65 | Nicaragua | Apoyo | *A. chancho* | Ben | Pap |
| 11NIce75 | Nicaragua | Apoyo | *A. chancho* | Ben | Pap |
| 11NIcf54 | Nicaragua | Apoyo | *A. chancho* | Ben | Pap |
| 11NIcf78 | Nicaragua | Apoyo | *A. chancho* | Ben | Pap |
| 11NIcg05 | Nicaragua | Apoyo | *A. chancho* | Ben | Pap |
| 11NIcg07 | Nicaragua | Apoyo | *A. chancho* | Ben | Pap |
| 14NIgc75 | Nicaragua | Apoyo | *A. chancho* | Ben | Pap |
| 14NIgc76 | Nicaragua | Apoyo | *A. chancho* | Ben | Pap |
| 14NIgc77 | Nicaragua | Apoyo | *A. chancho* | Ben | Pap |
| 14NIgc78 | Nicaragua | Apoyo | *A. chancho* | Ben | Pap |
| 14NIgc79 | Nicaragua | Apoyo | *A. chancho* | Ben | Pap |
| 14NIgc80 | Nicaragua | Apoyo | *A. chancho* | Ben | Pap |
| 14NIgc81 | Nicaragua | Apoyo | *A. chancho* | Ben | Pap |
| 14NIgd01 | Nicaragua | Apoyo | *A. chancho* | Ben | Pap |
| 14NIgd25 | Nicaragua | Apoyo | *A. chancho* | Ben | Pap |
| 14NIgd31 | Nicaragua | Apoyo | *A. chancho* | Ben | Pap |
| 16NIhc73 | Nicaragua | Apoyo | *A. chancho* | Ben | Pap |

| | | | | | |
|---|---|---|---|---|---|
| 16NIhd37 | Nicaragua | Apoyo | *A. chancho* | Ben | Pap |
| 09NI23d3 | Nicaragua | Apoyo | *A. chancho* | Ben | |
| 09NI23d4 | Nicaragua | Apoyo | *A. chancho* | Ben | |
| 10NIag32 | Nicaragua | Apoyo | *A. chancho* | Ben | |
| 10NIag48 | Nicaragua | Apoyo | *A. chancho* | Ben | |
| 14NIgd02 | Nicaragua | Apoyo | *A. chancho* | Ben | |
| 14NIgd23 | Nicaragua | Apoyo | *A. chancho* | Ben | |
| 14NIgd30 | Nicaragua | Apoyo | *A. chancho* | Ben | |
| 14NIgd32 | Nicaragua | Apoyo | *A. chancho* | Ben | |
| 14NIgd34 | Nicaragua | Apoyo | *A. chancho* | Ben | |
| 14NIgd35 | Nicaragua | Apoyo | *A. chancho* | Ben | |
| 16NIhc55 | Nicaragua | Apoyo | *A. chancho* | Ben | |
| 16NIhc63 | Nicaragua | Apoyo | *A. chancho* | Ben | |
| 16NIhd35 | Nicaragua | Apoyo | *A. chancho* | Ben | |
| 10NIag06 | Nicaragua | Apoyo | *A. chancho* | | Pap |
| 10NIag07 | Nicaragua | Apoyo | *A. chancho* | | Pap |
| 10NIag12 | Nicaragua | Apoyo | *A. chancho* | | Pap |
| 10NIag15 | Nicaragua | Apoyo | *A. chancho* | | Pap |
| 10NIag50 | Nicaragua | Apoyo | *A. chancho* | | Pap |
| 10NIag63 | Nicaragua | Apoyo | *A. chancho* | | Pap |
| 10NIag81 | Nicaragua | Apoyo | *A. chancho* | | Pap |
| 10NIaj81 | Nicaragua | Apoyo | *A. chancho* | | Pap |
| 10NIak12 | Nicaragua | Apoyo | *A. chancho* | | Pap |
| 10NIak13 | Nicaragua | Apoyo | *A. chancho* | | Pap |
| 10NIak24 | Nicaragua | Apoyo | *A. chancho* | | Pap |
| 11NIce77 | Nicaragua | Apoyo | *A. chancho* | | Pap |
| 11NIce78 | Nicaragua | Apoyo | *A. chancho* | | Pap |
| 11NIcf81 | Nicaragua | Apoyo | *A. chancho* | | Pap |
| 11NIcg16 | Nicaragua | Apoyo | *A. chancho* | | Pap |
| 13NIec15 | Nicaragua | Apoyo | *A. chancho* | | Pap |
| 13NIec76 | Nicaragua | Apoyo | *A. chancho* | | Pap |
| 13NIed26 | Nicaragua | Apoyo | *A. chancho* | | Pap |
| 13NIed32 | Nicaragua | Apoyo | *A. chancho* | | Pap |
| 13NIed38 | Nicaragua | Apoyo | *A. chancho* | | Pap |
| 14NIgc67 | Nicaragua | Apoyo | *A. chancho* | | Pap |
| 14NIgd03 | Nicaragua | Apoyo | *A. chancho* | | Pap |
| 14NIgd24 | Nicaragua | Apoyo | *A. chancho* | | Pap |
| 14NIgd33 | Nicaragua | Apoyo | *A. chancho* | | Pap |
| 14NIgd64 | Nicaragua | Apoyo | *A. chancho* | | Pap |
| 16NIhd41 | Nicaragua | Apoyo | *A. chancho* | | Pap |
| 10NIag45 | Nicaragua | Apoyo | *A. chancho* | | |
| 11NIce68 | Nicaragua | Apoyo | *A. zaliosus* | Lim | Pap |
| 11NIcf52 | Nicaragua | Apoyo | *A. zaliosus* | Lim | Pap |
| 11NIcg23 | Nicaragua | Apoyo | *A. zaliosus* | Lim | Pap |
| 14NIgc58 | Nicaragua | Apoyo | *A. zaliosus* | Lim | Pap |
| 14NIgc59 | Nicaragua | Apoyo | *A. zaliosus* | Lim | Pap |
| 14NIgc61 | Nicaragua | Apoyo | *A. zaliosus* | Lim | Pap |
| 14NIgc62 | Nicaragua | Apoyo | *A. zaliosus* | Lim | Pap |
| 14NIgc63 | Nicaragua | Apoyo | *A. zaliosus* | Lim | Pap |
| 14NIgc64 | Nicaragua | Apoyo | *A. zaliosus* | Lim | Pap |
| 14NIgd26 | Nicaragua | Apoyo | *A. zaliosus* | Lim | Pap |
| 14NIgd29 | Nicaragua | Apoyo | *A. zaliosus* | Lim | Pap |
| 14NIgd41 | Nicaragua | Apoyo | *A. zaliosus* | Lim | Pap |
| 14NIgd43 | Nicaragua | Apoyo | *A. zaliosus* | Lim | Pap |
| 14NIgd45 | Nicaragua | Apoyo | *A. zaliosus* | Lim | Pap |
| 14NIgd47 | Nicaragua | Apoyo | *A. zaliosus* | Lim | Pap |
| 14NIgd49 | Nicaragua | Apoyo | *A. zaliosus* | Lim | Pap |
| 16NIhc37 | Nicaragua | Apoyo | *A. zaliosus* | Lim | Pap |

| | | | | | |
|---|---|---|---|---|---|
| 16NIhc39 | Nicaragua | Apoyo | *A. zaliosus* | Lim | Pap |
| 16NIhc41 | Nicaragua | Apoyo | *A. zaliosus* | Lim | Pap |
| 16NIhc45 | Nicaragua | Apoyo | *A. zaliosus* | Lim | Pap |
| 16NIhc47 | Nicaragua | Apoyo | *A. zaliosus* | Lim | Pap |
| 16NIhc49 | Nicaragua | Apoyo | *A. zaliosus* | Lim | Pap |
| 16NIhc53 | Nicaragua | Apoyo | *A. zaliosus* | Lim | Pap |
| 16NIhd43 | Nicaragua | Apoyo | *A. zaliosus* | Lim | Pap |
| 09NI23d8 | Nicaragua | Apoyo | *A. zaliosus* | Lim | |
| 09NI23d9 | Nicaragua | Apoyo | *A. zaliosus* | Lim | |
| 09NI23e1 | Nicaragua | Apoyo | *A. zaliosus* | Lim | |
| 09NI23e2 | Nicaragua | Apoyo | *A. zaliosus* | Lim | |
| 09NI24b2 | Nicaragua | Apoyo | *A. zaliosus* | Lim | |
| 09NI24b5 | Nicaragua | Apoyo | *A. zaliosus* | Lim | |
| 09NI25f8 | Nicaragua | Apoyo | *A. zaliosus* | Lim | |
| 09NI25g7 | Nicaragua | Apoyo | *A. zaliosus* | Lim | |
| 09NI25g8 | Nicaragua | Apoyo | *A. zaliosus* | Lim | |
| 09NI25h3 | Nicaragua | Apoyo | *A. zaliosus* | Lim | |
| 10NIag21 | Nicaragua | Apoyo | *A. zaliosus* | Lim | |
| 11NIcf53 | Nicaragua | Apoyo | *A. zaliosus* | Lim | |
| 11NIcg14 | Nicaragua | Apoyo | *A. zaliosus* | Lim | |
| 13NIec74 | Nicaragua | Apoyo | *A. zaliosus* | Lim | |
| 14NIgc53 | Nicaragua | Apoyo | *A. zaliosus* | Lim | |
| 14NIgc54 | Nicaragua | Apoyo | *A. zaliosus* | Lim | |
| 14NIgd33 | Nicaragua | Apoyo | *A. zaliosus* | Lim | |
| 14NIgd42 | Nicaragua | Apoyo | *A. zaliosus* | Lim | |
| 14NIgd44 | Nicaragua | Apoyo | *A. zaliosus* | Lim | |
| 14NIgd48 | Nicaragua | Apoyo | *A. zaliosus* | Lim | |
| 14NIgd50 | Nicaragua | Apoyo | *A. zaliosus* | Lim | |
| 16NIhc51 | Nicaragua | Apoyo | *A. zaliosus* | Lim | |
| 16NIhc59 | Nicaragua | Apoyo | *A. zaliosus* | Lim | |
| 16NIhc61 | Nicaragua | Apoyo | *A. zaliosus* | Lim | |
| 16NIhd45 | Nicaragua | Apoyo | *A. zaliosus* | Lim | |
| 16NIhd47 | Nicaragua | Apoyo | *A. zaliosus* | Lim | |
| 10NIag09 | Nicaragua | Apoyo | *A. zaliosus* | | Pap |
| 10NIag11 | Nicaragua | Apoyo | *A. zaliosus* | | Pap |
| 10NIag55 | Nicaragua | Apoyo | *A. zaliosus* | | Pap |
| 10NIag71 | Nicaragua | Apoyo | *A. zaliosus* | | Pap |
| 10NIag75 | Nicaragua | Apoyo | *A. zaliosus* | | Pap |
| 10NIaj80 | Nicaragua | Apoyo | *A. zaliosus* | | Pap |
| 10NIak15 | Nicaragua | Apoyo | *A. zaliosus* | | Pap |
| 10NIak19 | Nicaragua | Apoyo | *A. zaliosus* | | Pap |
| 10NIak21 | Nicaragua | Apoyo | *A. zaliosus* | | Pap |
| 10NIak22 | Nicaragua | Apoyo | *A. zaliosus* | | Pap |
| 10NIak23 | Nicaragua | Apoyo | *A. zaliosus* | | Pap |
| 11NIcf79 | Nicaragua | Apoyo | *A. zaliosus* | | Pap |
| 13NIec80 | Nicaragua | Apoyo | *A. zaliosus* | | Pap |
| 14NIgc55 | Nicaragua | Apoyo | *A. zaliosus* | | Pap |
| 14NIgc56 | Nicaragua | Apoyo | *A. zaliosus* | | Pap |
| 14NIgc57 | Nicaragua | Apoyo | *A. zaliosus* | | Pap |
| 14NIgc60 | Nicaragua | Apoyo | *A. zaliosus* | | Pap |
| 14NIgc69 | Nicaragua | Apoyo | *A. zaliosus* | | Pap |
| 14NIgd04 | Nicaragua | Apoyo | *A. zaliosus* | | Pap |
| 14NIgd05 | Nicaragua | Apoyo | *A. zaliosus* | | Pap |
| 14NIgd46 | Nicaragua | Apoyo | *A. zaliosus* | | Pap |
| 05NI5014 | Nicaragua | Xiloá | *A. amarillo* | Ben | Mol |
| 05NI5043 | Nicaragua | Xiloá | *A. amarillo* | Ben | Mol |
| 05NI5068 | Nicaragua | Xiloá | *A. amarillo* | Ben | Mol |
| 05NI5083 | Nicaragua | Xiloá | *A. amarillo* | Ben | Mol |

| | | | | | |
|---|---|---|---|---|---|
| 05NI5240 | Nicaragua | Xiloá | *A. amarillo* | Ben | Mol |
| 05NI5241 | Nicaragua | Xiloá | *A. amarillo* | Ben | Mol |
| 05NI5245 | Nicaragua | Xiloá | *A. amarillo* | Ben | Mol |
| 05NI5291 | Nicaragua | Xiloá | *A. amarillo* | Ben | Mol |
| 05NI5459 | Nicaragua | Xiloá | *A. amarillo* | Ben | Mol |
| 11NIcc04 | Nicaragua | Xiloá | *A. amarillo* | Ben | Mol |
| 11NIcc05 | Nicaragua | Xiloá | *A. amarillo* | Ben | Mol |
| 11NIcc11 | Nicaragua | Xiloá | *A. amarillo* | Ben | Mol |
| 11NIcc12 | Nicaragua | Xiloá | *A. amarillo* | Ben | Mol |
| 11NIcc13 | Nicaragua | Xiloá | *A. amarillo* | Ben | Mol |
| 14NIga07 | Nicaragua | Xiloá | *A. amarillo* | Ben | Mol |
| 14NIga08 | Nicaragua | Xiloá | *A. amarillo* | Ben | Mol |
| 14NIgf42 | Nicaragua | Xiloá | *A. amarillo* | Ben | Mol |
| 14NIgf45 | Nicaragua | Xiloá | *A. amarillo* | Ben | Mol |
| 14NIgf62 | Nicaragua | Xiloá | *A. amarillo* | Ben | Mol |
| 14NIgf63 | Nicaragua | Xiloá | *A. amarillo* | Ben | Mol |
| 16NIha09 | Nicaragua | Xiloá | *A. amarillo* | Ben | Mol |
| 16NIha19 | Nicaragua | Xiloá | *A. amarillo* | Ben | Mol |
| 16NIha29 | Nicaragua | Xiloá | *A. amarillo* | Ben | Mol |
| 16NIha31 | Nicaragua | Xiloá | *A. amarillo* | Ben | Mol |
| 16NIha51 | Nicaragua | Xiloá | *A. amarillo* | Ben | Mol |
| 13NIeg14 | Nicaragua | Xiloá | *A. amarillo* | Ben | |
| 13NIeg18 | Nicaragua | Xiloá | *A. amarillo* | Ben | |
| 13NIeg19 | Nicaragua | Xiloá | *A. amarillo* | Ben | |
| 13NIeg21 | Nicaragua | Xiloá | *A. amarillo* | Ben | |
| 05NI5015 | Nicaragua | Xiloá | *A. amarillo* | | Mol |
| 05NI5021 | Nicaragua | Xiloá | *A. amarillo* | | Mol |
| 05NI5037 | Nicaragua | Xiloá | *A. amarillo* | | Mol |
| 05NI5039 | Nicaragua | Xiloá | *A. amarillo* | | Mol |
| 05NI5040 | Nicaragua | Xiloá | *A. amarillo* | | Mol |
| 05NI5041 | Nicaragua | Xiloá | *A. amarillo* | | Mol |
| 05NI5042 | Nicaragua | Xiloá | *A. amarillo* | | Mol |
| 05NI5044 | Nicaragua | Xiloá | *A. amarillo* | | Mol |
| 05NI5059 | Nicaragua | Xiloá | *A. amarillo* | | Mol |
| 05NI5060 | Nicaragua | Xiloá | *A. amarillo* | | Mol |
| 05NI5061 | Nicaragua | Xiloá | *A. amarillo* | | Mol |
| 05NI5063 | Nicaragua | Xiloá | *A. amarillo* | | Mol |
| 05NI5064 | Nicaragua | Xiloá | *A. amarillo* | | Mol |
| 05NI5065 | Nicaragua | Xiloá | *A. amarillo* | | Mol |
| 05NI5069 | Nicaragua | Xiloá | *A. amarillo* | | Mol |
| 05NI5070 | Nicaragua | Xiloá | *A. amarillo* | | Mol |
| 05NI5072 | Nicaragua | Xiloá | *A. amarillo* | | Mol |
| 05NI5073 | Nicaragua | Xiloá | *A. amarillo* | | Mol |
| 05NI5074 | Nicaragua | Xiloá | *A. amarillo* | | Mol |
| 05NI5075 | Nicaragua | Xiloá | *A. amarillo* | | Mol |
| 05NI5078 | Nicaragua | Xiloá | *A. amarillo* | | Mol |
| 05NI5079 | Nicaragua | Xiloá | *A. amarillo* | | Mol |
| 05NI5080 | Nicaragua | Xiloá | *A. amarillo* | | Mol |
| 05NI5081 | Nicaragua | Xiloá | *A. amarillo* | | Mol |
| 05NI5084 | Nicaragua | Xiloá | *A. amarillo* | | Mol |
| 05NI5085 | Nicaragua | Xiloá | *A. amarillo* | | Mol |
| 05NI5086 | Nicaragua | Xiloá | *A. amarillo* | | Mol |
| 05NI5087 | Nicaragua | Xiloá | *A. amarillo* | | Mol |
| 05NI5088 | Nicaragua | Xiloá | *A. amarillo* | | Mol |
| 05NI5089 | Nicaragua | Xiloá | *A. amarillo* | | Mol |
| 05NI5090 | Nicaragua | Xiloá | *A. amarillo* | | Mol |
| 05NI5091 | Nicaragua | Xiloá | *A. amarillo* | | Mol |
| 05NI5092 | Nicaragua | Xiloá | *A. amarillo* | | Mol |

| | | | | |
|---|---|---|---|---|
| 05NI5093 | Nicaragua | Xiloá | *A. amarillo* | Mol |
| 05NI5094 | Nicaragua | Xiloá | *A. amarillo* | Mol |
| 05NI5096 | Nicaragua | Xiloá | *A. amarillo* | Mol |
| 05NI5098 | Nicaragua | Xiloá | *A. amarillo* | Mol |
| 05NI5099 | Nicaragua | Xiloá | *A. amarillo* | Mol |
| 05NI5100 | Nicaragua | Xiloá | *A. amarillo* | Mol |
| 05NI5101 | Nicaragua | Xiloá | *A. amarillo* | Mol |
| 05NI5103 | Nicaragua | Xiloá | *A. amarillo* | Mol |
| 05NI5112 | Nicaragua | Xiloá | *A. amarillo* | Mol |
| 05NI5224 | Nicaragua | Xiloá | *A. amarillo* | Mol |
| 05NI5229 | Nicaragua | Xiloá | *A. amarillo* | Mol |
| 05NI5230 | Nicaragua | Xiloá | *A. amarillo* | Mol |
| 05NI5234 | Nicaragua | Xiloá | *A. amarillo* | Mol |
| 05NI5235 | Nicaragua | Xiloá | *A. amarillo* | Mol |
| 05NI5236 | Nicaragua | Xiloá | *A. amarillo* | Mol |
| 05NI5237 | Nicaragua | Xiloá | *A. amarillo* | Mol |
| 05NI5238 | Nicaragua | Xiloá | *A. amarillo* | Mol |
| 05NI5239 | Nicaragua | Xiloá | *A. amarillo* | Mol |
| 05NI5242 | Nicaragua | Xiloá | *A. amarillo* | Mol |
| 05NI5244 | Nicaragua | Xiloá | *A. amarillo* | Mol |
| 05NI5247 | Nicaragua | Xiloá | *A. amarillo* | Mol |
| 05NI5248 | Nicaragua | Xiloá | *A. amarillo* | Mol |
| 05NI5307 | Nicaragua | Xiloá | *A. amarillo* | Mol |
| 05NI5309 | Nicaragua | Xiloá | *A. amarillo* | Mol |
| 05NI5312 | Nicaragua | Xiloá | *A. amarillo* | Mol |
| 05NI5313 | Nicaragua | Xiloá | *A. amarillo* | Mol |
| 05NI5314 | Nicaragua | Xiloá | *A. amarillo* | Mol |
| 05NI5315 | Nicaragua | Xiloá | *A. amarillo* | Mol |
| 05NI5460 | Nicaragua | Xiloá | *A. amarillo* | Mol |
| 05NI5461 | Nicaragua | Xiloá | *A. amarillo* | Mol |
| 05NI5462 | Nicaragua | Xiloá | *A. amarillo* | Mol |
| 05NI5464 | Nicaragua | Xiloá | *A. amarillo* | Mol |
| 05NI5470 | Nicaragua | Xiloá | *A. amarillo* | Mol |
| 05NI5745 | Nicaragua | Xiloá | *A. amarillo* | Mol |
| 05NI5746 | Nicaragua | Xiloá | *A. amarillo* | Mol |
| 05NI5747 | Nicaragua | Xiloá | *A. amarillo* | Mol |
| 05NI5748 | Nicaragua | Xiloá | *A. amarillo* | Mol |
| 05NI5749 | Nicaragua | Xiloá | *A. amarillo* | Mol |
| 10NIal01 | Nicaragua | Xiloá | *A. amarillo* | Mol |
| 10NIal02 | Nicaragua | Xiloá | *A. amarillo* | Mol |
| 10NIal04 | Nicaragua | Xiloá | *A. amarillo* | Mol |
| 10NIal09 | Nicaragua | Xiloá | *A. amarillo* | Mol |
| 10NIal11 | Nicaragua | Xiloá | *A. amarillo* | Mol |
| 10NIal19 | Nicaragua | Xiloá | *A. amarillo* | Mol |
| 10NIal20 | Nicaragua | Xiloá | *A. amarillo* | Mol |
| 10NIal21 | Nicaragua | Xiloá | *A. amarillo* | Mol |
| 10NIal24 | Nicaragua | Xiloá | *A. amarillo* | Mol |
| 10NIal27 | Nicaragua | Xiloá | *A. amarillo* | Mol |
| 10NIal28 | Nicaragua | Xiloá | *A. amarillo* | Mol |
| 10NIal35 | Nicaragua | Xiloá | *A. amarillo* | Mol |
| 10NIal69 | Nicaragua | Xiloá | *A. amarillo* | Mol |
| 10NIan26 | Nicaragua | Xiloá | *A. amarillo* | Mol |
| 10NIan61 | Nicaragua | Xiloá | *A. amarillo* | Mol |
| 10NIap81 | Nicaragua | Xiloá | *A. amarillo* | Mol |
| 10NIaq14 | Nicaragua | Xiloá | *A. amarillo* | Mol |
| 10NIaq15 | Nicaragua | Xiloá | *A. amarillo* | Mol |
| 10NIaq17 | Nicaragua | Xiloá | *A. amarillo* | Mol |
| 10NIaq20 | Nicaragua | Xiloá | *A. amarillo* | Mol |

| | | | | | |
|---|---|---|---|---|---|
| 10NIaq21 | Nicaragua | Xiloá | *A. amarillo* | | Mol |
| 10NIaq22 | Nicaragua | Xiloá | *A. amarillo* | | Mol |
| 10NIaq23 | Nicaragua | Xiloá | *A. amarillo* | | Mol |
| 10NIaq26 | Nicaragua | Xiloá | *A. amarillo* | | Mol |
| 10NIaq27 | Nicaragua | Xiloá | *A. amarillo* | | Mol |
| 10NIaq30 | Nicaragua | Xiloá | *A. amarillo* | | Mol |
| 13NIea16 | Nicaragua | Xiloá | *A. amarillo* | | Mol |
| 13NIea17 | Nicaragua | Xiloá | *A. amarillo* | | Mol |
| 13NIea19 | Nicaragua | Xiloá | *A. amarillo* | | Mol |
| 13NIea20 | Nicaragua | Xiloá | *A. amarillo* | | Mol |
| 14NIga10 | Nicaragua | Xiloá | *A. amarillo* | | Mol |
| 14NIga11 | Nicaragua | Xiloá | *A. amarillo* | | Mol |
| 16NIha11 | Nicaragua | Xiloá | *A. amarillo* | | Mol |
| 16NIha13 | Nicaragua | Xiloá | *A. amarillo* | | Mol |
| 16NIha15 | Nicaragua | Xiloá | *A. amarillo* | | Mol |
| 16NIha17 | Nicaragua | Xiloá | *A. amarillo* | | Mol |
| 16NIha21 | Nicaragua | Xiloá | *A. amarillo* | | Mol |
| 16NIha23 | Nicaragua | Xiloá | *A. amarillo* | | Mol |
| 16NIha25 | Nicaragua | Xiloá | *A. amarillo* | | Mol |
| 16NIha27 | Nicaragua | Xiloá | *A. amarillo* | | Mol |
| 16NIha33 | Nicaragua | Xiloá | *A. amarillo* | | Mol |
| 16NIha35 | Nicaragua | Xiloá | *A. amarillo* | | Mol |
| 16NIha37 | Nicaragua | Xiloá | *A. amarillo* | | Mol |
| 05NI5019 | Nicaragua | Xiloá | *A. sagittae* | Lim | Pap |
| 05NI5055 | Nicaragua | Xiloá | *A. sagittae* | Lim | Pap |
| 05NI5274 | Nicaragua | Xiloá | *A. sagittae* | Lim | Pap |
| 05NI5283 | Nicaragua | Xiloá | *A. sagittae* | Lim | Pap |
| 05NI5287 | Nicaragua | Xiloá | *A. sagittae* | Lim | Pap |
| 05NI5538 | Nicaragua | Xiloá | *A. sagittae* | Lim | Pap |
| 05NI5539 | Nicaragua | Xiloá | *A. sagittae* | Lim | Pap |
| 05NI5540 | Nicaragua | Xiloá | *A. sagittae* | Lim | Pap |
| 05NI5542 | Nicaragua | Xiloá | *A. sagittae* | Lim | Pap |
| 05NI5543 | Nicaragua | Xiloá | *A. sagittae* | Lim | Pap |
| 05NI5544 | Nicaragua | Xiloá | *A. sagittae* | Lim | Pap |
| 05NI5546 | Nicaragua | Xiloá | *A. sagittae* | Lim | Pap |
| 05NI5547 | Nicaragua | Xiloá | *A. sagittae* | Lim | Pap |
| 14NIga02 | Nicaragua | Xiloá | *A. sagittae* | Lim | Pap |
| 14NIga34 | Nicaragua | Xiloá | *A. sagittae* | Lim | Pap |
| 14NIga35 | Nicaragua | Xiloá | *A. sagittae* | Lim | Pap |
| 14NIga37 | Nicaragua | Xiloá | *A. sagittae* | Lim | Pap |
| 14NIgf53 | Nicaragua | Xiloá | *A. sagittae* | Lim | Pap |
| 14NIgf64 | Nicaragua | Xiloá | *A. sagittae* | Lim | Pap |
| 14NIgf66 | Nicaragua | Xiloá | *A. sagittae* | Lim | Pap |
| 16NIha05 | Nicaragua | Xiloá | *A. sagittae* | Lim | Pap |
| 16NIha07 | Nicaragua | Xiloá | *A. sagittae* | Lim | Pap |
| 16NIhb01 | Nicaragua | Xiloá | *A. sagittae* | Lim | Pap |
| 16NIhb05 | Nicaragua | Xiloá | *A. sagittae* | Lim | Pap |
| 05NI5035 | Nicaragua | Xiloá | *A. sagittae* | Lim | |
| 05NI5548 | Nicaragua | Xiloá | *A. sagittae* | Lim | |
| 11NIcc06 | Nicaragua | Xiloá | *A. sagittae* | Lim | |
| 13NIeg09 | Nicaragua | Xiloá | *A. sagittae* | Lim | |
| 05NI5016 | Nicaragua | Xiloá | *A. sagittae* | | Pap |
| 05NI5017 | Nicaragua | Xiloá | *A. sagittae* | | Pap |
| 05NI5020 | Nicaragua | Xiloá | *A. sagittae* | | Pap |
| 05NI5038 | Nicaragua | Xiloá | *A. sagittae* | | Pap |
| 05NI5056 | Nicaragua | Xiloá | *A. sagittae* | | Pap |
| 05NI5226 | Nicaragua | Xiloá | *A. sagittae* | | Pap |
| 05NI5266 | Nicaragua | Xiloá | *A. sagittae* | | Pap |

| | | | | | |
|---|---|---|---|---|---|
| 05NI5267 | Nicaragua | Xiloá | *A. sagittae* | | Pap |
| 05NI5268 | Nicaragua | Xiloá | *A. sagittae* | | Pap |
| 05NI5269 | Nicaragua | Xiloá | *A. sagittae* | | Pap |
| 05NI5270 | Nicaragua | Xiloá | *A. sagittae* | | Pap |
| 05NI5271 | Nicaragua | Xiloá | *A. sagittae* | | Pap |
| 05NI5272 | Nicaragua | Xiloá | *A. sagittae* | | Pap |
| 05NI5273 | Nicaragua | Xiloá | *A. sagittae* | | Pap |
| 05NI5282 | Nicaragua | Xiloá | *A. sagittae* | | Pap |
| 05NI5284 | Nicaragua | Xiloá | *A. sagittae* | | Pap |
| 05NI5285 | Nicaragua | Xiloá | *A. sagittae* | | Pap |
| 05NI5286 | Nicaragua | Xiloá | *A. sagittae* | | Pap |
| 05NI5288 | Nicaragua | Xiloá | *A. sagittae* | | Pap |
| 05NI5289 | Nicaragua | Xiloá | *A. sagittae* | | Pap |
| 05NI5290 | Nicaragua | Xiloá | *A. sagittae* | | Pap |
| 05NI5292 | Nicaragua | Xiloá | *A. sagittae* | | Pap |
| 05NI5293 | Nicaragua | Xiloá | *A. sagittae* | | Pap |
| 05NI5295 | Nicaragua | Xiloá | *A. sagittae* | | Pap |
| 05NI5296 | Nicaragua | Xiloá | *A. sagittae* | | Pap |
| 05NI5297 | Nicaragua | Xiloá | *A. sagittae* | | Pap |
| 05NI5299 | Nicaragua | Xiloá | *A. sagittae* | | Pap |
| 05NI5302 | Nicaragua | Xiloá | *A. sagittae* | | Pap |
| 05NI5304 | Nicaragua | Xiloá | *A. sagittae* | | Pap |
| 05NI5324 | Nicaragua | Xiloá | *A. sagittae* | | Pap |
| 05NI5450 | Nicaragua | Xiloá | *A. sagittae* | | Pap |
| 05NI5472 | Nicaragua | Xiloá | *A. sagittae* | | Pap |
| 05NI5473 | Nicaragua | Xiloá | *A. sagittae* | | Pap |
| 05NI5541 | Nicaragua | Xiloá | *A. sagittae* | | Pap |
| 05NI5729 | Nicaragua | Xiloá | *A. sagittae* | | Pap |
| 05NI5730 | Nicaragua | Xiloá | *A. sagittae* | | Pap |
| 05NI5731 | Nicaragua | Xiloá | *A. sagittae* | | Pap |
| 05NI5737 | Nicaragua | Xiloá | *A. sagittae* | | Pap |
| 05NI5741 | Nicaragua | Xiloá | *A. sagittae* | | Pap |
| 05NI5743 | Nicaragua | Xiloá | *A. sagittae* | | Pap |
| 05NI5744 | Nicaragua | Xiloá | *A. sagittae* | | Pap |
| 05NI5750 | Nicaragua | Xiloá | *A. sagittae* | | Pap |
| 05NI5751 | Nicaragua | Xiloá | *A. sagittae* | | Pap |
| 05NI5752 | Nicaragua | Xiloá | *A. sagittae* | | Pap |
| 10NIal03 | Nicaragua | Xiloá | *A. sagittae* | | Pap |
| 10NIal12 | Nicaragua | Xiloá | *A. sagittae* | | Pap |
| 10NIal16 | Nicaragua | Xiloá | *A. sagittae* | | Pap |
| 10NIal57 | Nicaragua | Xiloá | *A. sagittae* | | Pap |
| 10NIal70 | Nicaragua | Xiloá | *A. sagittae* | | Pap |
| 10NIal72 | Nicaragua | Xiloá | *A. sagittae* | | Pap |
| 10NIan05 | Nicaragua | Xiloá | *A. sagittae* | | Pap |
| 10NIan06 | Nicaragua | Xiloá | *A. sagittae* | | Pap |
| 10NIan08 | Nicaragua | Xiloá | *A. sagittae* | | Pap |
| 10NIan10 | Nicaragua | Xiloá | *A. sagittae* | | Pap |
| 10NIan12 | Nicaragua | Xiloá | *A. sagittae* | | Pap |
| 10NIan14 | Nicaragua | Xiloá | *A. sagittae* | | Pap |
| 10NIan29 | Nicaragua | Xiloá | *A. sagittae* | | Pap |
| 14NIgf68 | Nicaragua | Xiloá | *A. sagittae* | | Pap |
| 14NIgf69 | Nicaragua | Xiloá | *A. sagittae* | | Pap |
| 16NIha01 | Nicaragua | Xiloá | *A. sagittae* | | Pap |
| 16NIhb03 | Nicaragua | Xiloá | *A. sagittae* | | Pap |
| 16NIhb09 | Nicaragua | Xiloá | *A. sagittae* | | Pap |
| 16NIhb15 | Nicaragua | Xiloá | *A. sagittae* | | Pap |
| 16NIhb17 | Nicaragua | Xiloá | *A. sagittae* | | Pap |
| 05NI5018 | Nicaragua | Xiloá | *A. xiloaensis* | Ben | Pap |

| | | | | | |
|---|---|---|---|---|---|
| 14NIga25 | Nicaragua | Xiloá | *A. xiloaensis* | Ben | Pap |
| 14NIga26 | Nicaragua | Xiloá | *A. xiloaensis* | Ben | Pap |
| 14NIga27 | Nicaragua | Xiloá | *A. xiloaensis* | Ben | Pap |
| 14NIga28 | Nicaragua | Xiloá | *A. xiloaensis* | Ben | Pap |
| 14NIga29 | Nicaragua | Xiloá | *A. xiloaensis* | Ben | Pap |
| 14NIga30 | Nicaragua | Xiloá | *A. xiloaensis* | Ben | Pap |
| 14NIga31 | Nicaragua | Xiloá | *A. xiloaensis* | Ben | Pap |
| 14NIga32 | Nicaragua | Xiloá | *A. xiloaensis* | Ben | Pap |
| 14NIga33 | Nicaragua | Xiloá | *A. xiloaensis* | Ben | Pap |
| 14NIga36 | Nicaragua | Xiloá | *A. xiloaensis* | Ben | Pap |
| 14NIgf35 | Nicaragua | Xiloá | *A. xiloaensis* | Ben | Pap |
| 14NIgf36 | Nicaragua | Xiloá | *A. xiloaensis* | Ben | Pap |
| 14NIgf38 | Nicaragua | Xiloá | *A. xiloaensis* | Ben | Pap |
| 14NIgf43 | Nicaragua | Xiloá | *A. xiloaensis* | Ben | Pap |
| 14NIgf44 | Nicaragua | Xiloá | *A. xiloaensis* | Ben | Pap |
| 14NIgf46 | Nicaragua | Xiloá | *A. xiloaensis* | Ben | Pap |
| 14NIgf50 | Nicaragua | Xiloá | *A. xiloaensis* | Ben | Pap |
| 14NIgf55 | Nicaragua | Xiloá | *A. xiloaensis* | Ben | Pap |
| 16NIha45 | Nicaragua | Xiloá | *A. xiloaensis* | Ben | Pap |
| 16NIha47 | Nicaragua | Xiloá | *A. xiloaensis* | Ben | Pap |
| 16NIha49 | Nicaragua | Xiloá | *A. xiloaensis* | Ben | Pap |
| 16NIha55 | Nicaragua | Xiloá | *A. xiloaensis* | Ben | Pap |
| 16NIha57 | Nicaragua | Xiloá | *A. xiloaensis* | Ben | Pap |
| 16NIha59 | Nicaragua | Xiloá | *A. xiloaensis* | Ben | Pap |
| 16NIha61 | Nicaragua | Xiloá | *A. xiloaensis* | Ben | Pap |
| 16NIha63 | Nicaragua | Xiloá | *A. xiloaensis* | Ben | Pap |
| 16NIha71 | Nicaragua | Xiloá | *A. xiloaensis* | Ben | Pap |
| 16NIha73 | Nicaragua | Xiloá | *A. xiloaensis* | Ben | Pap |
| 05NI5026 | Nicaragua | Xiloá | *A. xiloaensis* | Ben | |
| 05NI5032 | Nicaragua | Xiloá | *A. xiloaensis* | Ben | |
| 13NIeg10 | Nicaragua | Xiloá | *A. xiloaensis* | Ben | |
| 14NIgf49 | Nicaragua | Xiloá | *A. xiloaensis* | Ben | |
| 14NIgf65 | Nicaragua | Xiloá | *A. xiloaensis* | Ben | |
| 16NIhb24 | Nicaragua | Xiloá | *A. xiloaensis* | Ben | |
| 05NI5054 | Nicaragua | Xiloá | *A. xiloaensis* | | Pap |
| 05NI5057 | Nicaragua | Xiloá | *A. xiloaensis* | | Pap |
| 05NI5058 | Nicaragua | Xiloá | *A. xiloaensis* | | Pap |
| 05NI5076 | Nicaragua | Xiloá | *A. xiloaensis* | | Pap |
| 05NI5077 | Nicaragua | Xiloá | *A. xiloaensis* | | Pap |
| 05NI5104 | Nicaragua | Xiloá | *A. xiloaensis* | | Pap |
| 05NI5105 | Nicaragua | Xiloá | *A. xiloaensis* | | Pap |
| 05NI5108 | Nicaragua | Xiloá | *A. xiloaensis* | | Pap |
| 05NI5109 | Nicaragua | Xiloá | *A. xiloaensis* | | Pap |
| 05NI5110 | Nicaragua | Xiloá | *A. xiloaensis* | | Pap |
| 05NI5275 | Nicaragua | Xiloá | *A. xiloaensis* | | Pap |
| 05NI5276 | Nicaragua | Xiloá | *A. xiloaensis* | | Pap |
| 05NI5277 | Nicaragua | Xiloá | *A. xiloaensis* | | Pap |
| 05NI5278 | Nicaragua | Xiloá | *A. xiloaensis* | | Pap |
| 05NI5279 | Nicaragua | Xiloá | *A. xiloaensis* | | Pap |
| 05NI5280 | Nicaragua | Xiloá | *A. xiloaensis* | | Pap |
| 05NI5281 | Nicaragua | Xiloá | *A. xiloaensis* | | Pap |
| 05NI5294 | Nicaragua | Xiloá | *A. xiloaensis* | | Pap |
| 05NI5300 | Nicaragua | Xiloá | *A. xiloaensis* | | Pap |
| 05NI5327 | Nicaragua | Xiloá | *A. xiloaensis* | | Pap |
| 05NI5448 | Nicaragua | Xiloá | *A. xiloaensis* | | Pap |
| 05NI5449 | Nicaragua | Xiloá | *A. xiloaensis* | | Pap |
| 05NI5454 | Nicaragua | Xiloá | *A. xiloaensis* | | Pap |
| 05NI5468 | Nicaragua | Xiloá | *A. xiloaensis* | | Pap |

| | | | | |
|---|---|---|---|---|
| 05NI5469 | Nicaragua | Xiloá | *A. xiloaensis* | Pap |
| 05NI5471 | Nicaragua | Xiloá | *A. xiloaensis* | Pap |
| 05NI5474 | Nicaragua | Xiloá | *A. xiloaensis* | Pap |
| 05NI5477 | Nicaragua | Xiloá | *A. xiloaensis* | Pap |
| 05NI5753 | Nicaragua | Xiloá | *A. xiloaensis* | Pap |
| 05NI5754 | Nicaragua | Xiloá | *A. xiloaensis* | Pap |
| 05NI5755 | Nicaragua | Xiloá | *A. xiloaensis* | Pap |
| 05NI5756 | Nicaragua | Xiloá | *A. xiloaensis* | Pap |
| 10NIal50 | Nicaragua | Xiloá | *A. xiloaensis* | Pap |
| 10NIal54 | Nicaragua | Xiloá | *A. xiloaensis* | Pap |
| 10NIal64 | Nicaragua | Xiloá | *A. xiloaensis* | Pap |
| 10NIal65 | Nicaragua | Xiloá | *A. xiloaensis* | Pap |
| 10NIal66 | Nicaragua | Xiloá | *A. xiloaensis* | Pap |
| 10NIal73 | Nicaragua | Xiloá | *A. xiloaensis* | Pap |
| 10NIan03 | Nicaragua | Xiloá | *A. xiloaensis* | Pap |
| 10NIan32 | Nicaragua | Xiloá | *A. xiloaensis* | Pap |
| 10NIan60 | Nicaragua | Xiloá | *A. xiloaensis* | Pap |
| 10NIan69 | Nicaragua | Xiloá | *A. xiloaensis* | Pap |
| 10NIaq16 | Nicaragua | Xiloá | *A. xiloaensis* | Pap |
| 14NIgf37 | Nicaragua | Xiloá | *A. xiloaensis* | Pap |
| 14NIgf39 | Nicaragua | Xiloá | *A. xiloaensis* | Pap |
| 14NIgf40 | Nicaragua | Xiloá | *A. xiloaensis* | Pap |
| 14NIgf41 | Nicaragua | Xiloá | *A. xiloaensis* | Pap |
| 14NIgf51 | Nicaragua | Xiloá | *A. xiloaensis* | Pap |
| 14NIgf52 | Nicaragua | Xiloá | *A. xiloaensis* | Pap |
| 14NIgf54 | Nicaragua | Xiloá | *A. xiloaensis* | Pap |
| 14NIgf58 | Nicaragua | Xiloá | *A. xiloaensis* | Pap |
| 14NIgf59 | Nicaragua | Xiloá | *A. xiloaensis* | Pap |
| 14NIgf61 | Nicaragua | Xiloá | *A. xiloaensis* | Pap |
| 16NIha53 | Nicaragua | Xiloá | *A. xiloaensis* | Pap |
| 16NIha65 | Nicaragua | Xiloá | *A. xiloaensis* | Pap |
| 16NIha67 | Nicaragua | Xiloá | *A. xiloaensis* | Pap |
| 16NIha69 | Nicaragua | Xiloá | *A. xiloaensis* | Pap |
