## Supplementary Material for "Exploring phenotypic diversity: a comparative analysis of cichlid convergence"

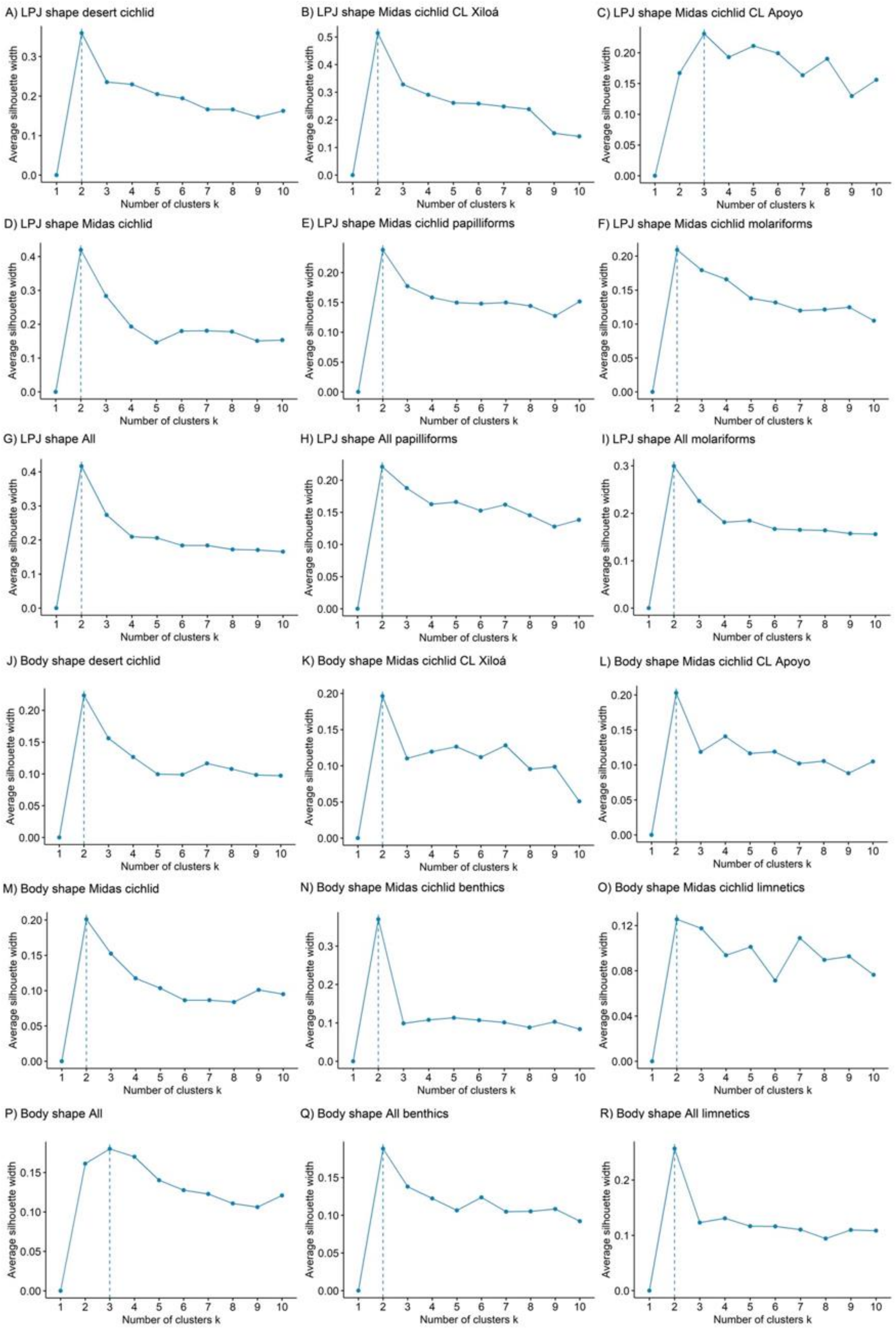


**Figure S1.** Optimal number of clusters obtained by the Silhouette method of the k-means clustering analysis. All analyses recovered and optimal number of clusters of two, except for the analysis LPJ shape within CL Apoyo in Nicaragua (C), and the analysis of body shape including both geographical regions (P), which recovered three clusters in both cases.

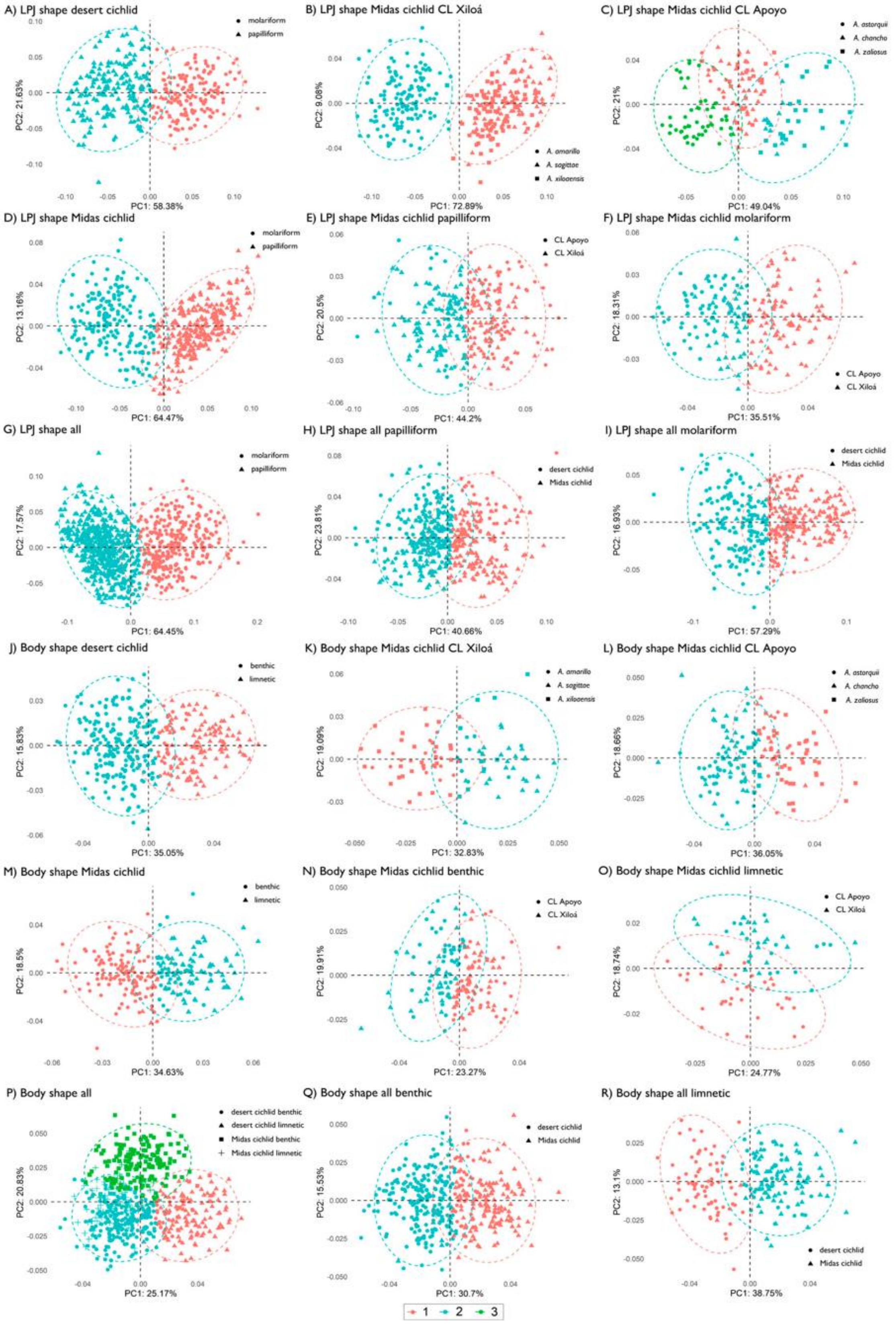


**Figure S2**. PCAs showing the results of the clustering analysis with the K-means algorithm. Colors represent the different clusters resolved by the analyses. Symbols represent the species or morphotypes that are being compared.

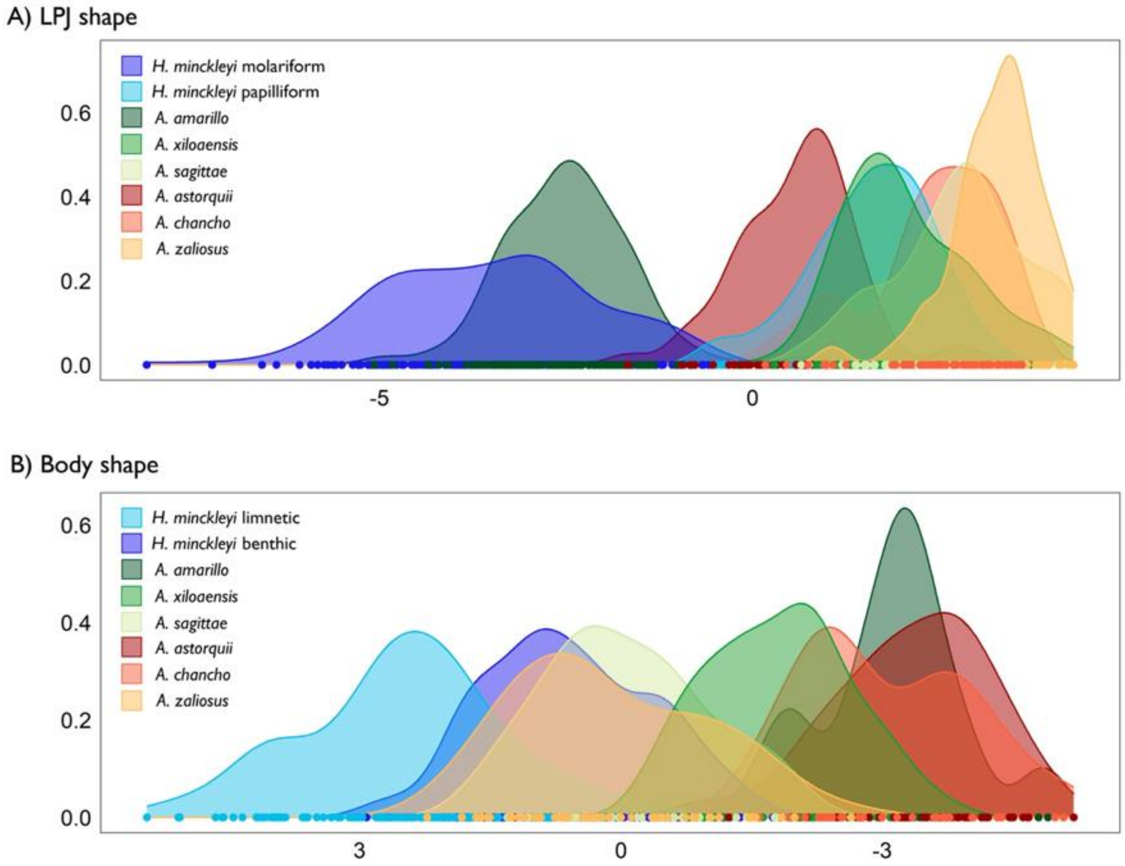


**Figure S3**. Linear discriminant function analyses differentiating among all species within the two cichlid radiations in jaw (A) and body (B) shape. LPJ density plots show that molarization is stronger in the desert cichlid, followed by CL Xiloá. Molariform jaws in CL Apoyo show considerable overlap with papilliform jaws in the desert cichlid and even in CL Xiloá. Body density plots evidence dissimilar strategies in limnetic fish in both cichlid radiations.

**Table S1.** Detailed list of the samples included in gene expression analyses. Pap and Mol refer to papilliform and molariform LPJ morphotypes respectively. Ben and Lim refer to benthic and limnetic body shape morphotypes respectively.

| ID | SYSTEM | LAKE | SPECIES | BODY | JAW |
|---|---|---|---|---|---|
| **BA14** | Mexico | Anteojo | *H. minckleyi* | Lim | Pap |
| **BA79** | Mexico | Becerra | *H. minckleyi* | Lim | Pap |
| **BB03** | Mexico | Becerra | *H. minckleyi* | Lim | Pap |
| **BB28** | Mexico | Becerra | *H. minckleyi* | Lim | Mol |
| **BB23** | Mexico | Becerra | *H. minckleyi* | Lim | Mol |
| **BA02** | Mexico | Churince | *H. minckleyi* | Ben | Mol |
| **BB59** | Mexico | Escobedo | *H. minckleyi* | Lim | Mol |
| **BB60** | Mexico | Escobedo | *H. minckleyi* | Lim | Mol |
| **BB40** | Mexico | Pozas Azules | *H. minckleyi* | Ben | Pap |
| **BB66** | Mexico | Tío Cándido | *H. minckleyi* | Lim | Pap |
| **23F3** | Nicaragua | Apoyo | *A. astorquii* | Ben | Mol |
| **23A4** | Nicaragua | Apoyo | *A. chancho* | Ben | Pap |
| **24A6** | Nicaragua | Apoyo | *A. chancho* | Ben | Pap |
| **24A7** | Nicaragua | Apoyo | *A. chancho* | Ben | Pap |
| **25H2** | Nicaragua | Apoyo | *A. zaliosus* | Lim | Pap |
| **24B2** | Nicaragua | Apoyo | *A. zaliosus* | Lim | Pap |
| **24B6** | Nicaragua | Apoyo | *A. zaliosus* | Lim | Pap |
| **24A8** | Nicaragua | Apoyo | *A. zaliosus* | Lim | Pap |
| **27A7** | Nicaragua | Xiloá | *A. amarillo* | Ben | Mol |
| **27A8** | Nicaragua | Xiloá | *A. amarillo* | Ben | Mol |
| **24G2** | Nicaragua | Xiloá | *A. amarillo* | Ben | Mol |
| **27A6** | Nicaragua | Xiloá | *A. sagittae* | Lim | Pap |
| **27G8** | Nicaragua | Xiloá | *A. sagittae* | Lim | Pap |
| **27A9** | Nicaragua | Xiloá | *A. xiloensis* | Ben | Pap |
| **23D1** | Nicaragua | Xiloá | *A. xiloensis* | Ben | Pap |

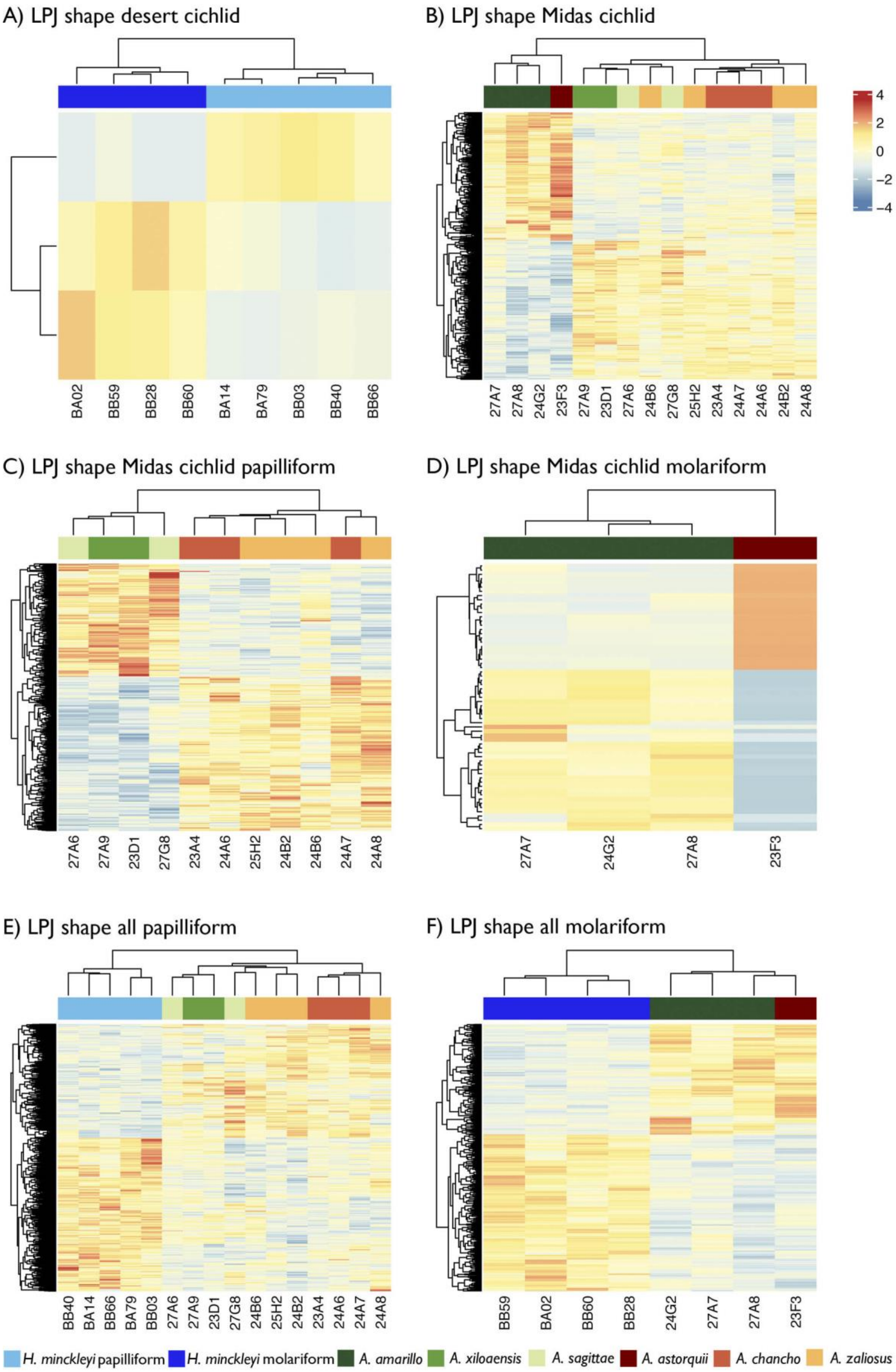


**Figure S4**. Heatmaps of the general patterns of gene regulation of the differentially expressed genes in the analyses by LPJ morphotypes within and between geographical regions.

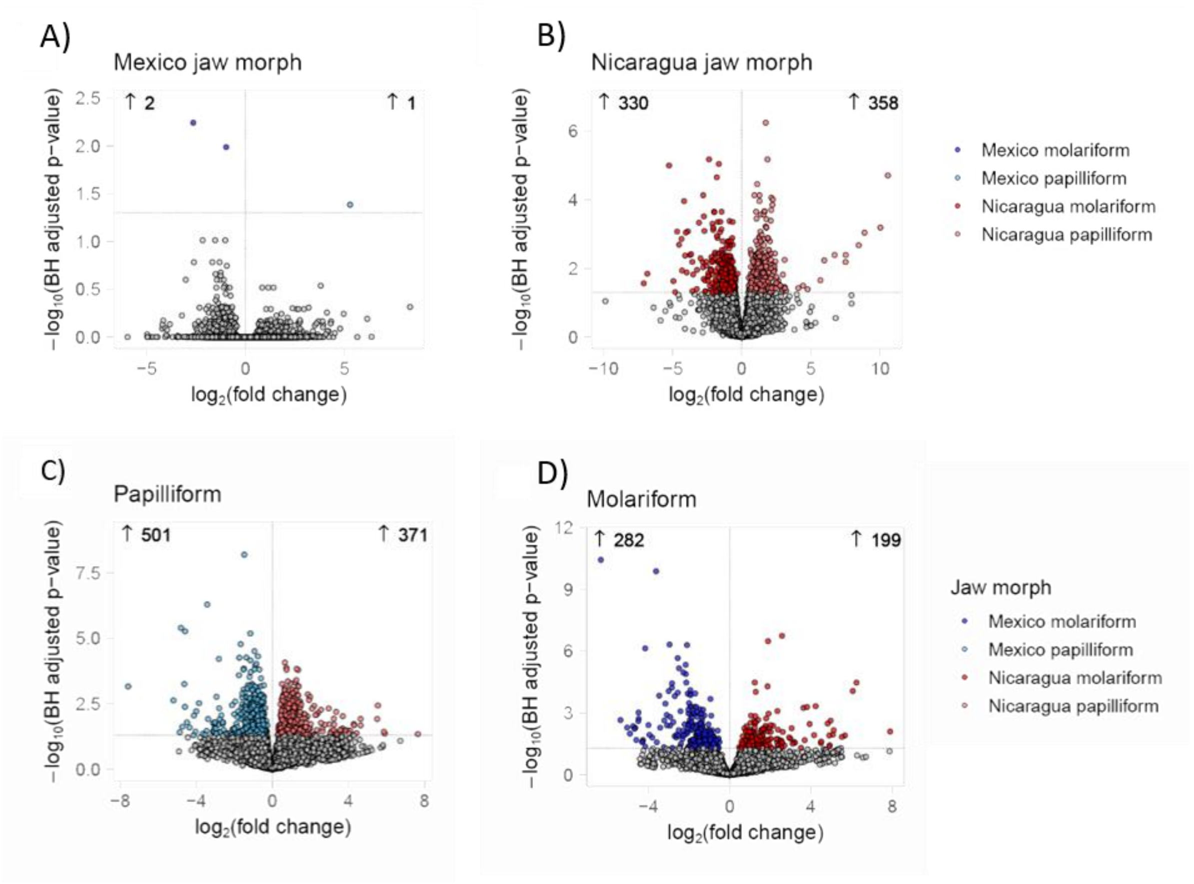


**Figure S5**. Volcano plots of the different comparisons of DEG between jaws. Each dot represents a gene, with grey dots below the threshold horizontal line indicating genes with statistically insignificant change in DEG.

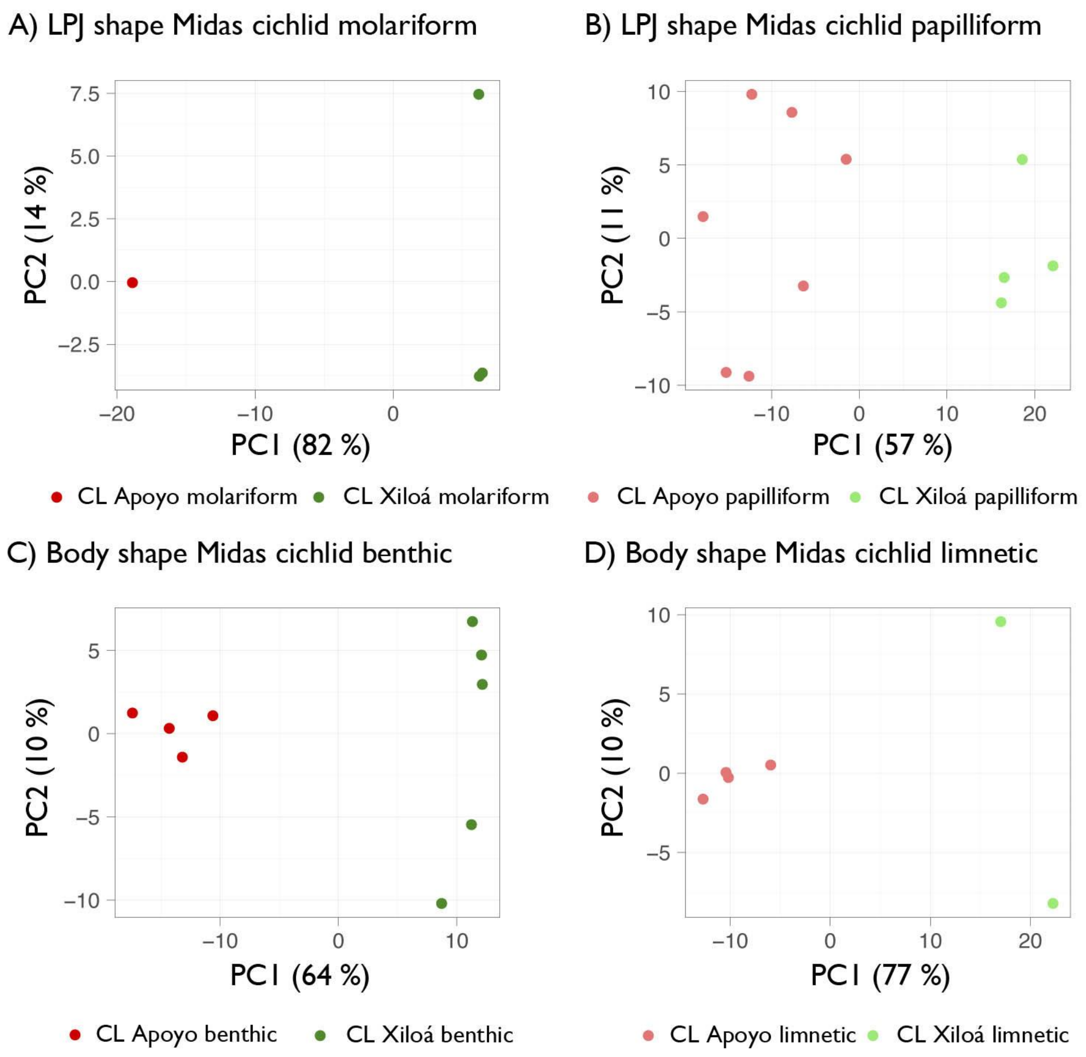


**Figure S6.** PCA of gene expression levels in the comparisons between morphotypes within the Midas cichlid complex. Dots represent individuals. Axes represent the first two PCs, with the percentage of the variance explained by each of them indicated. (A) Molariform LPJ. (B) Papilliform LPJ. (C) Benthic body shape. (D) Limnetic body shape.

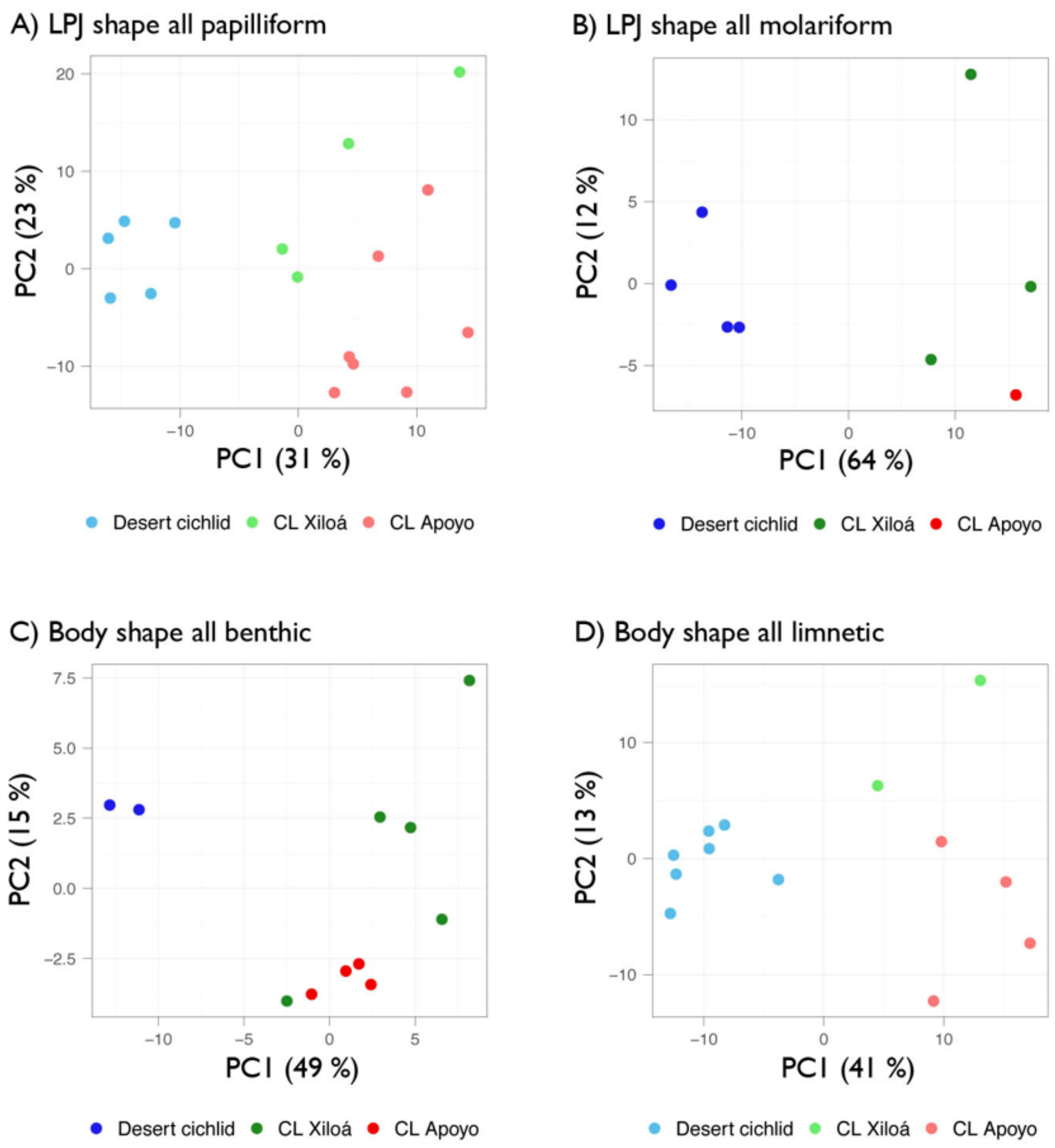


**Figure S7**. PCAs of gene expression levels in the comparisons between papilliform (A) and molariform (B) LPJs and benthic (C) and limnetic (D) body morphotypes including both geographical regions. Dots represent individuals. Axes represent the first two PCs, with the percentage of the variance explained by each of them indicated.

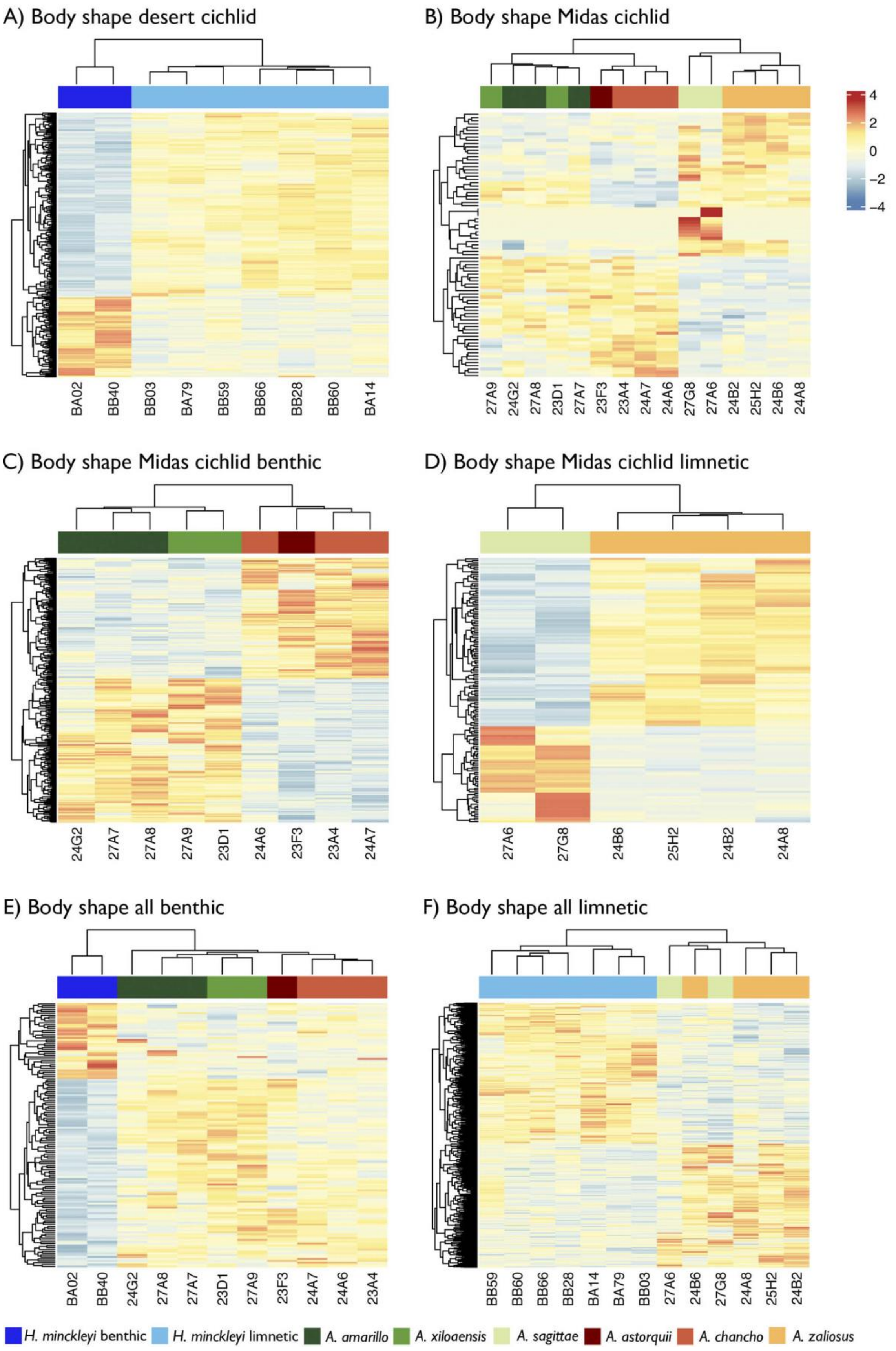


**Figure S8**. Heatmaps of the general patterns of gene regulation of the differentially expressed genes in the analyses by body shape morphotypes within and between geographical regions.

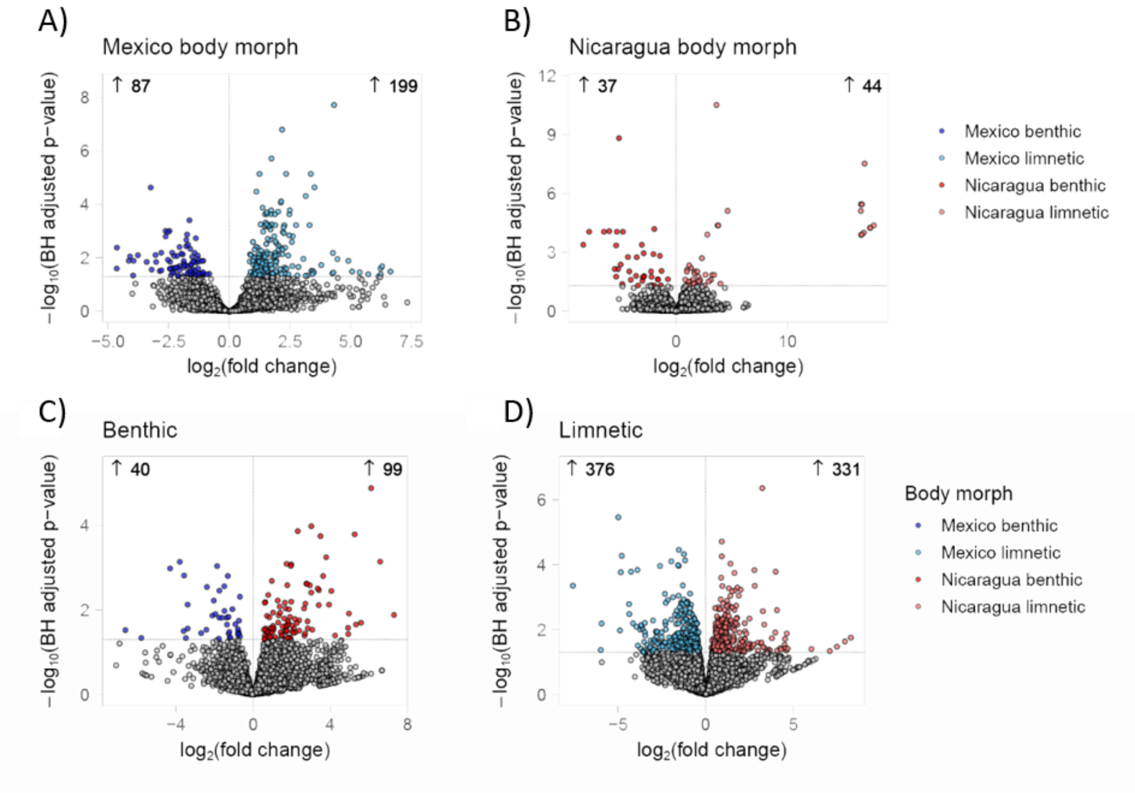


**Figure S9**. Volcano plots of the different comparisons of DEG between bodies. Each dot represents a gene, with grey dots below the threshold horizontal line indicating genes with statistically insignificant change in DEG.
